# DrugMap: A quantitative pan-cancer analysis of cysteine ligandability

**DOI:** 10.1101/2023.10.20.563287

**Authors:** Mariko Takahashi, Harrison B. Chong, Siwen Zhang, Matthew J. Lazarov, Stefan Harry, Michelle Maynard, Ryan White, Heather E. Murrey, Brendan Hilbert, Jason R. Neil, Magdy Gohar, Maolin Ge, Junbing Zhang, Benedikt R. Durr, Gregory Kryukov, Chih-Chiang Tsou, Natasja Brooijmans, Aliyu Sidi Omar Alghali, Karla Rubio, Antonio Vilanueva, Drew Harrison, Ann-Sophie Koglin, Samuel Ojeda, Barbara Karakyriakou, Alexander Healy, Jonathan Assaad, Farah Makram, Inbal Rachman, Neha Khandelwal, Pei-Chieh Tien, George Popoola, Nicholas Chen, Kira Vordermark, Marianne Richter, Himani Patel, Tzu-yi Yang, Hanna Griesshaber, Tobias Hosp, Sanne van den Ouweland, Toshiro Hara, Lily Bussema, Rui Dong, Lei Shi, Martin Q. Rasmussen, Ana Carolina Domingues, Aleigha Lawless, Jacy Fang, Satoshi Yoda, Linh Phuong Nguyen, Sarah Marie Reeves, Farrah Nicole Wakefield, Adam Acker, Sarah Elizabeth Clark, Taronish Dubash, David E. Fisher, Shyamala Maheswaran, Daniel A. Haber, Genevieve Boland, Moshe Sade-Feldman, Russel Jenkins, Aaron Hata, Nabeel Bardeesy, Mario L. Suva, Brent Martin, Brian Liau, Christopher Ott, Miguel N. Rivera, Michael S. Lawrence, Liron Bar-Peled

**Author notes:** These authors contributed equally (Mariko Takahashi, Harrison B. Chong, Siwen Zhang). Correspondence should be addressed to M.L. and L.B-P.

## Abstract

Cysteine-focused chemical proteomic platforms have accelerated the clinical development of covalent inhibitors of a wide-range of targets in cancer. However, how different oncogenic contexts influence cysteine targeting remains unknown. To address this question, we have developed *DrugMap*, an atlas of cysteine ligandability compiled across 416 cancer cell lines. We unexpectedly find that cysteine ligandability varies across cancer cell lines, and we attribute this to differences in cellular redox states, protein conformational changes, and genetic mutations. Leveraging these findings, we identify actionable cysteines in NFκB1 and SOX10 and develop corresponding covalent ligands that block the activity of these transcription factors. We demonstrate that the NFκB1 probe blocks DNA binding, whereas the SOX10 ligand increases SOX10-SOX10 interactions and disrupts melanoma transcriptional signaling. Our findings reveal heterogeneity in cysteine ligandability across cancers, pinpoint cell-intrinsic features driving cysteine targeting, and illustrate the use of covalent probes to disrupt oncogenic transcription factor activity.

## Introduction

The molecular and genetic characterization of cancer, and more recently high-throughput genomic and functional interrogations of malignancies, have revealed the pathways that cancers exploit to survive^1–6^. Findings from these investigations have been leveraged to develop blockbuster medicines that target oncogenes such as EGFR^7, 8^, ALK^9, 10^, BRAF^11, 12^ and BTK^13–15^. These same approaches have also begun to identify the mechanisms by which cancers rewire or mutate oncogenic pathways to evade targeted agents. However, for the more than 400 oncogenic drivers discovered to date, less than 10% have been drugged, and this small subset is almost entirely composed of protein kinases or enzymes^16^. It remains unclear whether small-molecule targeting can be systematically extended to other classes of oncogenic drivers.

In the pantheon of anti-cancer drugs, covalent inhibitors targeting the amino acid cysteine have recently gained traction, and drugs bearing irreversible cysteine-reactive warheads have met with clinical success^17, 18^. Prominent examples include osimertinib^8, 19^ for the treatment of EGFR-mutant non-small cell lung cancers (NSCLCs), ibrutinib^14, 15^ for the targeting of BTK in B cell lymphomas, and sotorasib^20, 21^ for inhibiting mutant KRAS•G12C in NSCLCs. Despite their efficacy in some patients, there is wide variability in responses, thought to stem from inter- and intra-tumoral heterogeneity^22–26^. However, how this heterogeneity determines the ability of drugs to engage their corresponding target is not well understood.

Concomitant with the rise of covalent inhibitors in the clinic, chemical proteomic methods have now become a popular approach to quantitatively and globally measure cysteine reactivity changes that are dependent on particular metabolic states or physiologies, or are influenced by covalent small-molecule inhibitors^27–35^. Advances in these cysteine-focused chemical proteomics approaches now allow for the routine quantification of 10,000+ cysteines per run, and multiplexing technologies have enabled the identification of corresponding targets for hundreds of covalent fragments^17, 36, 37^. Empowered by these technological advances, we set out to perform a series of chemical proteomic experiments to develop a quantitative portrait of cysteine ligandability across 400+ cancer cell lines, thereby establishing an initial framework for a cancer ligandability map. We refer to this dataset as *DrugMap*. By profiling changes in cysteine reactivity using broadly reactive electrophilic compounds and combining this information with comprehensive genomic characterization, we reveal widespread heterogeneity in cysteine targeting. This variability can be explained, in part, by variation in the intrinsic redox environments of cancer cells, as well as cysteine-proximal mutations in proteins. By integrating protein structural datasets with chemical proteomic data, we uncover ligand- and mutation-induced conformational changes that can be read out as changes in cysteine ligandability. Leveraging the findings from DrugMap, we develop a covalent ligand for the oncogenic transcription factor NFkB1 that functions through disruption of DNA interactions, as well as a covalent ligand for SOX10 that induces oligomerization by acting as a ‘molecular glue’, disrupting oncogenic transcriptional circuits and ultimately blocking proliferation of melanoma. The development of DrugMap, and these two novel anti-oncogenic agents it enabled, represents an opportunity to begin systematically uncovering the rules governing cysteine ligandability, providing a roadmap for oncology-focused ligand discovery and drug development.

## Results

### Development of a Pan-Cancer *DrugMap*

To develop a comprehensive portrait of cysteine ligandability across multiple cancers, we selected 416 cell lines representative of 25 cancer subtypes (“lineages”), each represented on average by ∼18 cell lines, with rarer cancers (e.g. Merkel cell carcinoma) covered by a minimum of two (**Figure 1A**, **Table S1**). We integrated the isoTOP-ABBP platform with tandem mass tag (TMT) - based mass spectrometry quantification (iso-TMT) to measure cysteine reactivity^28, 35, 37, 38^. In this approach, cell lysates are first treated with cysteine-reactive “scout” compounds or vehicle control, allowing reactive cysteines a chance to form covalent adducts, and then this is followed by a chase with a pan-cysteine-reactive probe (iodoacetamide-desthiobiotin, DBIA^37^) which reacts with all remaining free cysteine thiolate groups. Crucially, cysteines that reacted with the scout compound will escape being tagged by DBIA (**Figure 1A**). Throughout this work, we define ligandable cysteines as those which are engaged (ε-value) >60% by cysteine-reactive compounds (**Figures 1, S1A-B**, see Methods). Given the breadth of cell lines profiled, we wished to identify covalent scout fragments that broadly represent cysteine engagement by larger sets of covalent fragments. To select scouts, we first mapped cysteines targeted by a larger set of 152 covalent fragments, finding that three covalent scout probes, previously characterized in cysteine ligandability studies^39–41^, engage ∼74% of cysteines covered by the larger fragment library (**Figures S1C-E**). Leveraging these three scout probes and the iso-TMT method, we systematically measured changes in cysteine reactivity across our pan-cancer cohort of 416 cancer cell lines, which we divided across 420+ multiplex proteomics experiments, in total encompassing ∼1,100 hours of mass spectrometry analysis (**Table S2**). We achieved deep cysteine coverage, quantifying a median of 14,000+ unique cysteine-containing peptides per cell line. The abundances of these cysteines correlated well to corresponding RNA expression and DNA copy number (**Figures 1A, S1F, K**). In aggregate, we quantified 78,778 cysteines across 23,016 protein isoforms, finding that 5999 cysteines are ligandable (**Figure 1A, S1G-J**, see Methods).

**Figure 1:**
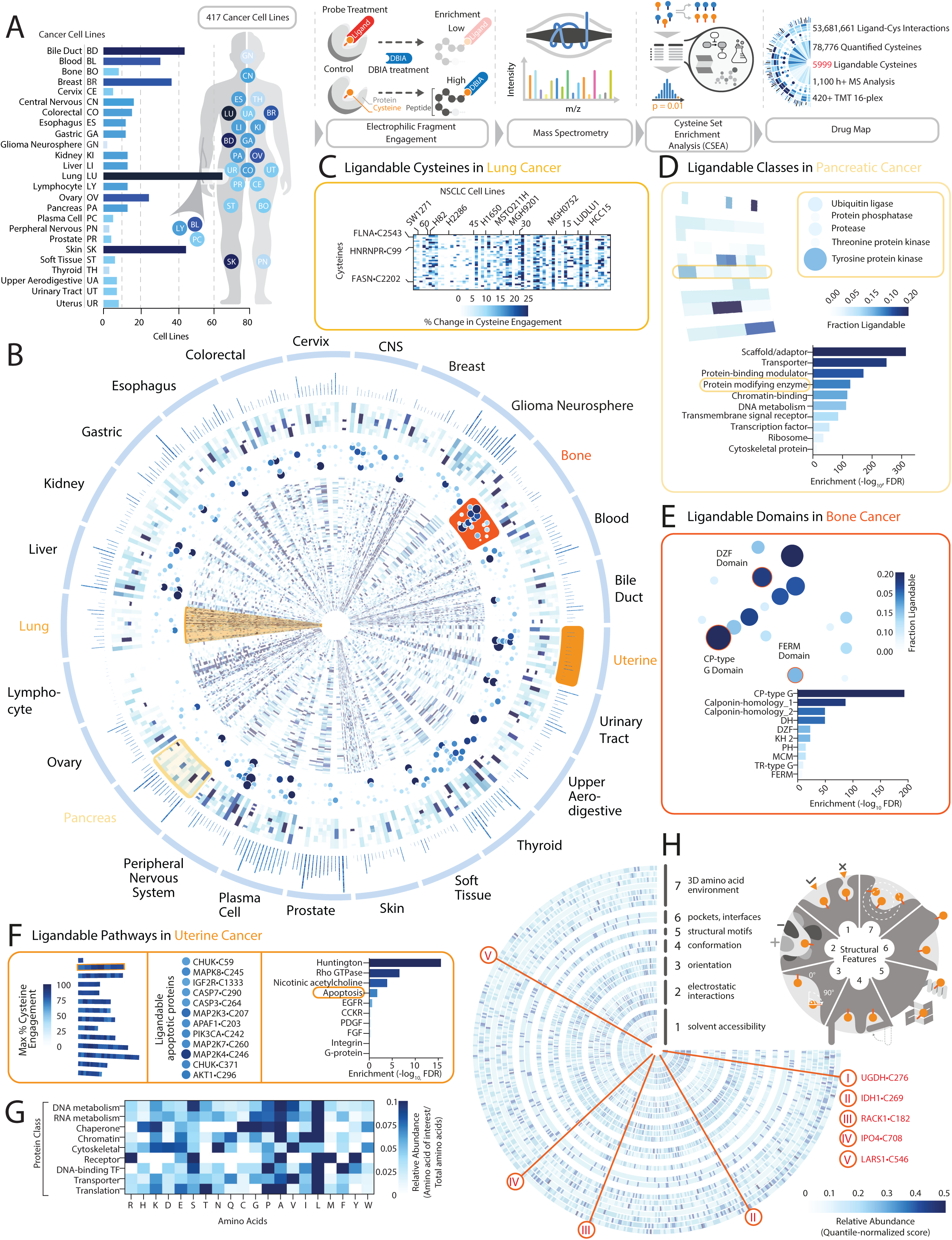
Defining a cysteine ligandability map across cancer. (A) Development of a Pan-Cancer ligandability map. Cancer cell lines analyzed in this study, and schematic of DrugMap development (see also **Table S1**). (B) Mapping ligandability across 25 cancer types. Circular heatmap depicting differences in cysteine, domain, class and pathway ligandability (moving from inner- to outermost layers) across 416 cell lines. (see also **Table S2** and Methods). (C) Examples of differences in cysteine ligandability among 64 lung cancer cell lines. (D) Cysteine set enrichment analysis (CSEA) plot of protein classes with ligandable cysteines identified in twelve pancreatic cancer cells (see **Figure S2A**, text, and Methods). (E) Protein domains with ligandable cysteines enriched in nine bone cancer cell lines. (F) Pathways with ligandable cysteines enriched in nine uterine cancer cell lines. (G) Amino acid neighbors of ligandable cysteines. Heatmap depicting relative abundance of residues within a 6 Å radius of ligandable cysteines for each protein class (see also **Figures S3F-G**, Methods). TF: transcription factors. (H) Structural characterization of ligandable cysteines. >20 structural features were calculated for each of the 632 ligandable cysteines identified in a corresponding protein structure (See also **Figures S3C-E, Table S13** and Methods).

To enable a systematic analysis of cysteine ligandability, we developed a computational analysis method called cysteine set enrichment analysis (CSEA), based on commonly used gene-centric enrichment score algorithms^42, 43^. However, rather than highlighting gene-level feature enrichment, CSEA determines enrichment signals at cysteine-level resolution (**Figure S2A**). This analytical pipeline leverages a repository of 6,000+ unique cysteine sets compiled from molecular features (e.g. domains, proteins and pathways^42, 44^), cysteine reactivity studies (e.g. cysteine reactivity changes in diverse physiological contexts, or triggered by covalent small-molecule profiling^37, 45^), and biochemical-structural features (e.g. surface accessibility, cysteine depth^46–51^) (**Figures S2A, Table S16**). When applied across all cancer lineages, CSEA revealed that microtubule-binding proteins are disproportionately ligandable. Looking at individual lineages, we found that proteins that function as intramolecular scaffolds are enriched for cysteine ligandability in pancreatic cancers **(Figures 1D, S2B).** Proteins with CP-type G domains^52^ are highly ligandable across multiple cancer cell lines, including bone cancers (**Figures 1E, S2B**). Among cellular pathways underlying tumorigenesis, CSEA revealed that the EGF-JAK-STAT pathway is enriched across all cancer cell lines profiled and the Rho GTPase pathway is enriched in uterine cancers (**Figure 1F, S2D**). PI3K signaling provides an example of how ligandabilty is altered across proteins that are members of this signaling pathway, across individual cysteines in those proteins, and among different cell lines, including lymphomas, liver cancers, and pancreatic cancers (**Figure S2E**). Transcription factors, which have historically been considered difficult to target^53^, include multiple family members with ligandable cysteines, most notably homeodomain-containing transcription factors (**Figure S2C**). DrugMap thus reveals a rich detailed landscape of cysteine ligandability across cancer.

Despite the relative structural simplicity and broad reactivity of the three covalent scout probes, we observed distinct liganding events for each, with KB03 showing the greatest unique ligandability (**Figure S2F**). The distinct reactivity of KB03 led us to ask whether we could infer protein-structural correlates of cysteine liganding. To this end, we trained a composite feed-forward convolutional neural network on structural parameters computed across 55,000+ human protein structures^48, 49^. Leveraging the >9,000 cysteines identified in DrugMap having corresponding structural annotation, this allowed us to predict KB03 ligandability with ∼70% accuracy (**Figure S3B**). The predictive ability of this neural network suggested that coherent structural principles of cysteine liganding can be gleaned from this dataset. Thus, we ran CSEA over our library of structural cysteine sets, to identify structural features determining cysteine ligandability. This analysis revealed strong enrichment of ligandable cysteines in alpha-helices (**Figure S3A-B, E**). Subsequent analyses revealed that cysteines that are highly solvent-accessible (those lying <2Å from bulk solvent and having a coordination number of <25) or deeply buried (>8Å from bulk solvent with a coordination number of >50) are both disfavored for liganding (**Figures 1H, S3D-E**). In contrast, cysteines displaying intermediate burial (on average 4-6Å from bulk solvent, with a coordination number of 30-40) are more ligandable (**Figures 1H, S3D**). We also quantified the nearest amino acid neighbors within a 6Å sphere around ligandable cysteines^32^, resulting in identification of amino acid neighbors associated with ligandability in specific protein families (**Figure 1G, S3F**). We found a general trend for enrichment of basic amino acids near ligandable cysteines, possibly stemming from modulation of cysteine pK_a_^54^ (**Figure S3G**). There was little correlation between structure-based and primary-sequence-based enrichment of residues near ligandable cysteines, emphasizing the importance of evaluating contributors to cysteine ligandability from a structural perspective (**Figure S3H**).

A critical determinant of protein druggability is the availability of a structural pocket that can accommodate small-molecule binding near a critical portion of a protein^55–59^. To identify potential protein pockets, we used P2Rank and DeepPocket, two artificial intelligence-based pocket prediction algorithms that deploy unique pocket segmentation strategies^60^, allowing us to identify >250,000 high-confidence pockets across 55,500 protein structures (**Figure S3H**). Among ligandable cysteines, ∼5% lay within a predicted pocket of >50 Å^3^ (**Figures S3I,K**). There is a higher abundance of ligandable cysteines in pockets localized to protein active sites and allosteric sites^61^, consistent with the known enrichment of binding pockets in these structural elements^39^ (**Figure S3L**). Reassuringly, cysteines residing within pockets were twice as likely to be liganded (**Figure S3J**). The amino acid content of pockets with ligandable cysteines favored aliphatic amino acids, notably leucine (**Figure S3M**). This accords with the enrichment found by CSEA of ligandable cysteines localized to alpha helices (**Figure S3C**).

In summary, by profiling reactivity changes in cysteines across 416 cell lines, we globally establish the identity of cysteines in cancers that can in principle be targeted by covalent drugs, and provide a complementary analytical platform to aid in the identification of molecular and structural features underlying cysteine ligandability.

### Heterogeneity in cysteine ligandability is in part defined by cancer cell redox states and protein conformational changes

Our analysis of ligandable cysteines in DrugMap revealed that a majority of ligandable cysteines are consistently engaged across cancers (**Figures 2A-B**). However, for ∼8% cysteines, we instead found heterogeneity in engagement, with these cysteines being ligandable in some cell lines but not others (**Figures 2A-B, S4A, Table S3**). To identify the molecular determinants governing heterogeneous cysteines, we employed CSEA. This analysis identified a strong tendency toward heterogeneous ligandability for cysteines regulated by redox processes (**Figures 2C, S4B**). This led us to directly examine the contribution of the cellular redox environment to ligandability. To do this, we pre-treated K562 cells with a cell-permeant reducing agent, tris(2-methoxycarboxyethyl)phosphine (TMCEP)^62^ prior to *in vitro* ligandabilty analysis. This revealed a substantial increase in liganding of cysteines that had shown heterogeneity of ligandability (**Figures 2D, S4C,F, Table S4A**). Further supporting the premise that intracellular redox environment impacts ligandability, we found that increasing the expression of NRF2, a master antioxidant regulator that promotes a reductive environment^63–66^, caused a concomitant increase in cysteine ligandability (**Figures 2D, S4D-F, Table S4B**). Importantly, there was strong overlap in effects on heterogeneous cysteine liganding following the two different perturbations (**Figures 2E, S4G**). NRF2 is commonly activated in cancers through mutation of its negative regulator KEAP1^67^. Consistent with this, we observed increased ligandability for a subset of heterogeneous cysteines in a panel of KEAP1-mutant cell lines compared to KEAP1-WT cells (**Figure S4H**). These results suggest a potential opportunity to differentially target proteins required for proliferation based on the metabolic state of a cancer cell.

**Figure 2:**
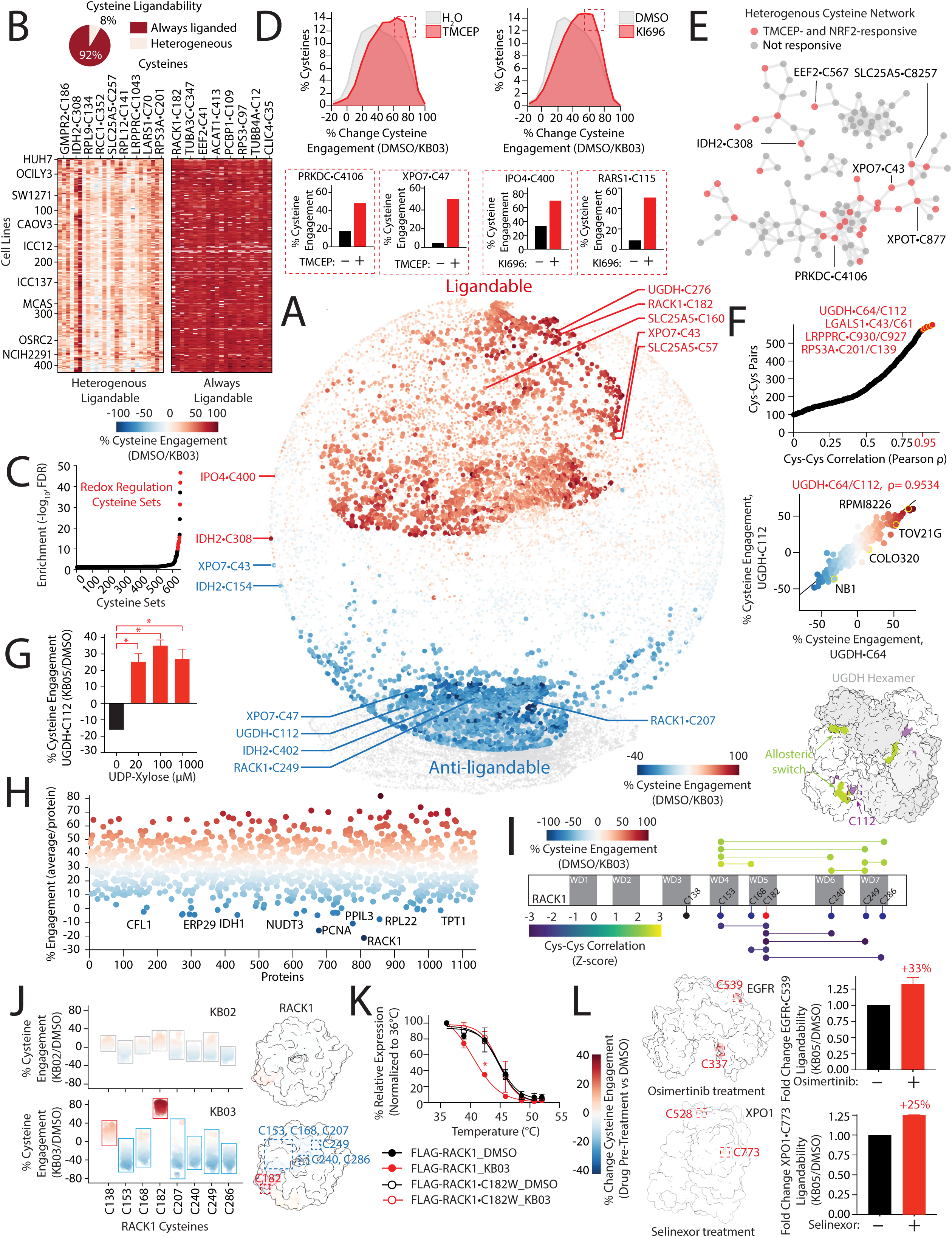
Heterogeneity in cysteine ligandability is driven by cellular redox state and protein conformational changes. (A) Characterizing distinct cysteine populations in DrugMap. Cysteine ligandability plot for ∼18,000 cysteines reveals ligandable (dark red), heterogeneous (light red) and anti-ligandable cysteines (blue) (see also **Figure S4A** and Methods). (B) Relative abundance of ligandable and heterogeneous cysteines. Top, proportion of ligandable cysteines that are homogeneous or heterogeneous in their engagement by scout probes. Bottom, notable examples of homogeneous or heterogeneous cysteine ligandability (see also **Table S3**). (C) Heterogeneous cysteines are enriched by CSEA for redox regulation. (D) Cellular redox state regulates heterogeneous cysteine engagement. Left, plot depicting changes in cysteine ligandability following tris(2-methoxycarbonylethyl)phosphine (TMCEP)^62^ treatment. Right, Plot depicting changes in cysteine engagement following activation of NRF2. Insets, examples of ligandability changes for heterogeneous cysteines following TMCEP treatment (bottom left) or NRF2 activation (bottom right). K562 cells were treated with TMCEP (1 mM) or vehicle (water) for 1 hr or were treated a NRF2 activator (1 µM of KI696^64, 66^) or vehicle (DMSO) for 48 hrs and changes in cysteine engagement by iso-TMT were determined following in lysate treatment with 500 µM KB03 (see also **Figure S4F** and **Table S4**). (E) Redox regulated heterogeneous cysteines. Network analysis of heterogeneous cysteines denoting cysteines whose ligandability changes following TMCEP and NRF2 activation. (F) UGDH•C64 and UGDH•C112 are highly correlated in their ligandability. Top, Plot of cys-cys engagement correlations across heterogeneous cysteines. Middle, UGDH•C64/C112 are tightly correlated in their heterogeneous ligandability across hundreds cell lines. Bottom, UGDH•C112 (purple) is proximal to UGDH allosteric switch (green) and protein-protein interface (PDB:5VR8^70^). (G) UGDH oligomerization following treatment with UDP-xylose increases UGDH•C112 ligandability. Engagement plot for UGDH•C112 following treatment with the indicated concentrations of UDP-xylose. Changes in cysteine ligandability were determined as described in (D) (see also **Table S5**). (H) Anti-ligandable cysteines identified across DrugMap. Average cysteine ligandability per protein is displayed, identifying RACK1 with the greatest degree of anti-ligandability following scout-probe treatment. (I) RACK1•C182 is anti-correlated in its ligandability with other cysteines in RACK1. Schematic depicting RACK1 cysteine engagement and cys-cys correlations. (J) RACK1 liganding results in increase in cysteine anti-ligandability. Left, RACK1 cysteine engagement following treatment of KB02 (which does not ligand RACK1•C182) and KB03 (which ligands RACK1•C182) across > 200 cell lines in DrugMap. Right, RACK1 structure with average cysteine ligandability displayed following treatment with KB02 or KB03 (PDB:4AOW^115^). (K) RACK1•C182 engagement decreases thermal stability. Thermal shift assay of RACK1-FLAG or RACK1•C182W-FLAG following treatment with 100 µM KB03. Expression of indicated proteins was determined by immunoblot (see also **Figure S5C**, Methods). (L) EGFR and XPO1 liganding alters cysteine accessibility, revealing newly ligandable sites. PC9 and K562 lysates were pre-treated with 1 µM of osimertinib and selinexor, respectively, and changes in cysteine engagement following treatment with KB05 is displayed on the structures of EGFR (PDB:3QWQ^116^) and XPO1 (PDB=3GB8^117^) (see also **Figure S5E**). Cysteine engagement in cell lysates was determined as described in (D). Data are represented as mean ± SD. *p < 0.05. Student’s *t*-test (two-tailed, unpaired) were used to determine statistical significance.

Next, we asked if a difference in the liganding of heterogeneous cysteines can function as a surrogate readout for cellular redox states. Accordingly, we computed a proteomic redox score for each cell line in DrugMap. Most cell lines had a proteomic redox score corresponding to higher levels of liganding, suggesting a more reduced cellular environment (**Figure S4I**). Focusing on a panel of ovarian cancer cell lines with diverse proteomic redox scores, we found that cells with high levels of heterogeneous cysteine liganding showed higher ROS induction following treatment with auranofin, a drug previously reported to increase ROS levels^32^ (**Figure S4J**). Taken together, these results support the hypothesis that a cancer cell’s redox state plays a substantial role in controlling cysteine ligandabilty, and that ligandability itself can be used as surrogate readout for measuring one dimension of the cellular redox environment.

Because cysteine liganding does not occur in isolation but rather in the context of a whole protein and all its interacting partners, we sought to identify structural features that drive heterogeneity in cysteine engagement. To do this, we computed pairwise cys-cys correlations across >4,000+ cysteines. This analysis revealed that cysteine pairs that are highly correlated in their ligandability tend to share common structural features, notably intermediate levels of residue burial (**Figure S4K**). Among the most highly correlated cysteines pairs were Cys64 and Cys112 of the protein UGDH, two cysteines that happen to belong to the core subset of heterogeneous cysteines identified above (**Table S3**). Because UGDH is known to undergo a conformational change as a mechanism controlling its catalytic activity^68–70^, and Cys112 in UGDH resides within an established allosteric regulatory switch (**Figure 2F, S4K**), we hypothesized that differences in ligandability may reflect different UGDH complex states. Consistent with prior studies, we found that treatment of UGDH with the allosteric modulator UDP-xylose^71^, led to a substantial increase in thermal stability, indicating an induction of conformational changes^72^ (**Figure S4L**). Addition of UDP-xylose to cell lysates prior to ligandabilty analysis resulted in a pronounced increase in UGDH•C112 liganding (**Figure 2G, Table S5**). These results strongly suggest that heterogeneity in cysteine ligandability is driven, in part, by protein conformational changes.

### Ligand-induced conformational changes promote greater cysteine accessibility

Chemical proteomic studies such as ours define cysteines as ligandable if their reactivity with pan-cysteine reactive probes (e.g. DBIA) *decreases* following treatment with scout probes (**Figure S4A**). Curiously, we found that for ∼5% of cysteines detected in >40 cell lines, treatment with scout probes paradoxically *increased* cysteine reactivity with DBIA, the opposite of the decreased reactivity seen for liganded cysteines (**Figures 2A, S4A**). We refer to these unusual cysteines as ‘anti-ligandable’ given their increase in accessibility following covalent inhibitor treatment (**Figure 2H, S4A**). We detected anti-ligandable cysteines across hundreds of cell lines in DrugMap, and we also found them to be pervasive in previously reported cysteine-focused chemical proteomic datasets^36, 37, 39, 41, 73^ **(Figures S5A-B**). Given that liganding of proteins by small molecules can result in substantial conformational rearrangements^74^, we hypothesized that covalent adduction of one cysteine within a protein might result in a conformational change that increases the accessibility to DBIA of another cysteine in the same protein. To test this hypothesis, we concentrated on proteins containing cysteines with the greatest anti-ligandability. This drew our attention to RACK1, a WD-40 containing protein that regulates protein translation^75, 76^ (**Figure 2I**). Cys-cys correlations of anti-ligandable cysteines within RACK1 revealed that RACK1•C182, a surface-exposed cysteine residing in a shallow groove, had the strongest negative correlations to other RACK1 cysteines across our entire dataset (**Figures 2I-J, S4K**). Nearly all other RACK1 cysteines were highly correlated and anti-ligandable (**Figure 2I**). Observing that RACK1•C182 is exclusively liganded by KB03 but not KB02 (**Figure S2F**), we compared DBIA accessibility for other anti-correlated RACK1 cysteines, and found a decrease in their engagement after KB03 but not KB02 treatment (**Figure 2J**). This decrease in engagement of RACK1 cysteines occurred across dozens of cell lines (**Figure 2J**). Given that these RACK1 cysteines are buried in the *apo* structure (**Figure 2J**), their decreased engagement suggests a potential conformational change. Using thermal shift assays, we found that KB03 decreases thermal stability of wildtype RACK1 but not of the RACK1•C182W mutant that blocks liganding of Cys182 (**Figure 2K**). Treatment with KB02, which did not bind RACK1•C182, had no impact on thermal stability in comparison to vehicle control (**Figure S5C**). These results provide strong evidence that RACK1 undergoes a conformational change upon Cys182 liganding. Interestingly, we uncovered several examples of paired ligandable and anti-ligandable cysteines in oncogenes identified in DrugMap, including XPO1 and EGFR (**Figure S5D**). To establish whether this finding can extend beyond broadly reactive covalent fragments, we treated lysates with osimertinib^77^ or selinexor^78^, drugs that target EGFR•C797 and XPO1•C528 respectively. We found corresponding alterations in DBIA accessibility for multiple cysteines in these proteins (**Figures 2L, S5E**). Treatment with these drugs also subsequently increased KB05 engagement of EGFR•C539 and XPO1•C773 (**Figures 2L, S5E, Table S6**), potentially suggesting the emergence of a cryptic pocket upon inhibitor binding. These results imply that protein conformational changes can be inferred at global scale in complex settings by monitoring differences in cysteine ligandability using chemical proteomics, thus outlining the beginnings of a new scalable approach to the study of protein structural dynamics.

### Genetic determinants of cysteine ligandability

To reveal how the landscape of cysteine ligandability may be influenced by a cancer’s genetic architecture, we leveraged the deep genomic and functional characterization that has been obtained for most of the cell lines profiled in DrugMap^79^. We applied hierarchical clustering to identify genetic features associated with cysteine ligandability and resolved three main clusters from this multi-omic analysis, segregating largely based on lineage (**Figure 3A**). We did not observe a bulk correlation between cysteine ligandability and RNA expression or gene essentiality (**Figures S6A-B**). However, a few cysteines showed strong correlations between ligandability and genomic or transcriptomic features (e.g. COL6A3•C775 and AGO•C328), potentially reflective of differences in protein complex formation that alter ligandability, and in accordance with protein complexes being tied to co-essentiality^80^ (**Figures S6A-B**). Strikingly, we observed a strong correlation between a cell line’s mutational burden and the prevalence of corresponding alterations in cysteine ligandability (**Figure S6C**), prompting us to investigate how single mutations associate with ligandability changes. We identified >50 *de novo* cysteine mutations (i.e. X–>Cys) that create novel ligandable cysteines, many in essential genes (**Figure S6D**). Far more prominent in our analysis were cysteine-proximal mutations that associated with changes in cysteine ligandability. Proximal mutations could be local (e.g. <5Å) or distal (e.g., >40Å) to a cysteine (**Figure 3B**). Some proximal mutations associated with increased cysteine ligandability occurred in key oncogenes such as MGMT and IDH1 (**Figure 3C**). Other mutations associated with cysteine anti-ligandability occurred in tumor suppressors such as MYH9 (**Figure 3D**). In HCC1395 cells, which harbor an allosteric F157L mutation in the antioxidant enzyme PRDX5, we saw an increase in ligandability of PRDX5•C100 relative to cell lines expressing WT PRDX5 (**Figure S6E**). Supporting the correlation between cysteine-proximal mutations and changes in ligandability, overexpression of the PRDX5^F157L^ mutant in HEK-293T cells increased Cys100 ligandability compared to its WT counterpart (**Figure S6E, Table S7**), demonstrating that mutations need not be local to alter cysteine liganding. To better understand associations between the spectrum of mutations in a protein and changes in cysteine ligandability, we performed logistic regression analysis, revealing that clusters of mutations in TP53 correlate to changes in TP53•C141 liganding (**Figures S6F-G**). Finally, we compared how pathway ligandability is impacted in cell lines driven by common oncogenic mutations. We identified, for example, that proteins involved in serine biosynthesis are particularly ligandable in cell lines with BRCA1 mutations (**Figure S6H**).

**Figure 3:**
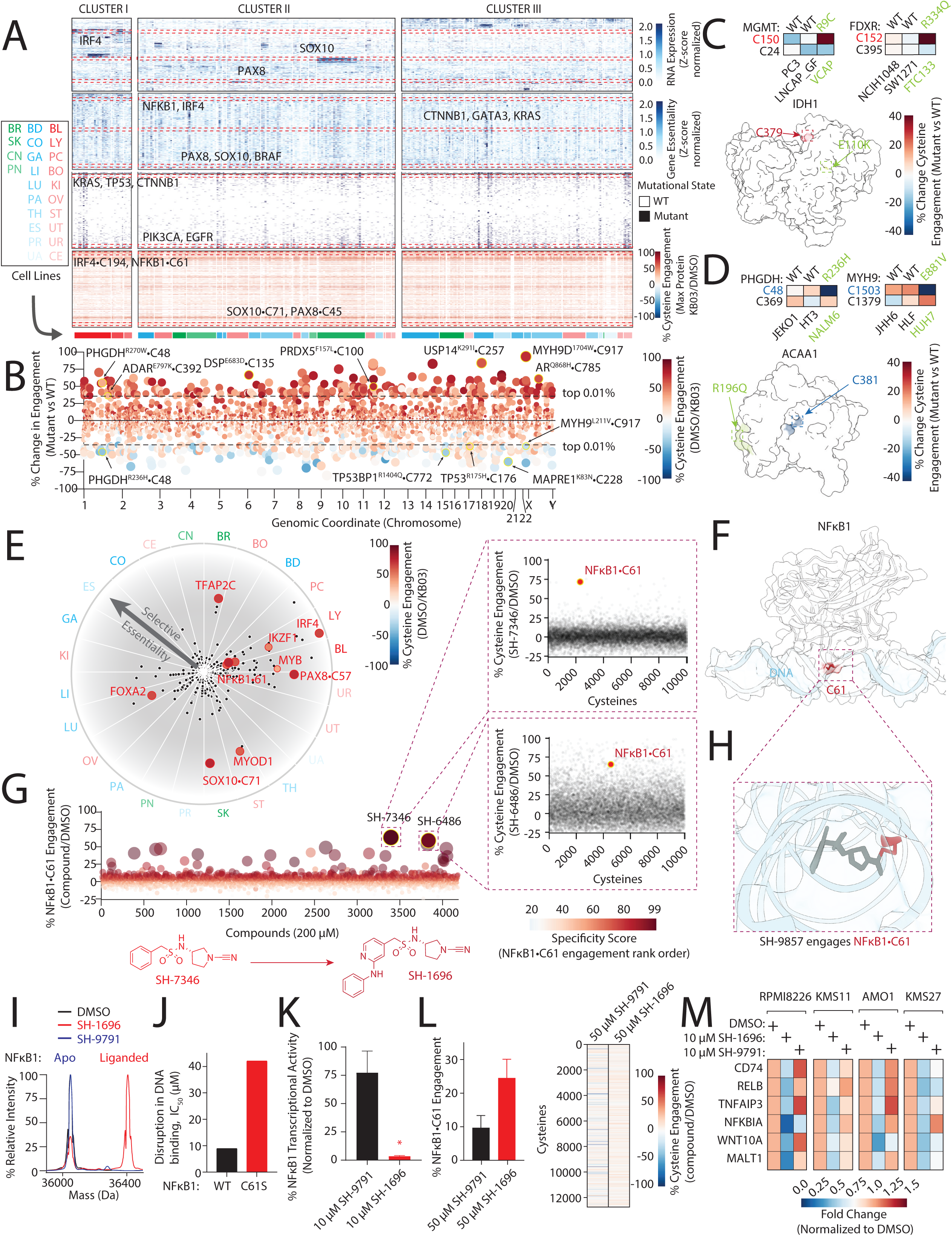
Genetic determinants of cysteine ligandabilty enable the development of a NFkB1 probe that disrupts DNA interaction. (A) Identifying genetic associates of cysteine ligandability across cancer. Multi-omic clustering based on mRNA expression, gene essentiality, mutational status and cysteine ligandability across 300 cell lines. (B) Engagement plot identifying intra-gene mutations that associate with corresponding changes in cysteine ligandability (see Methods). (C) Cysteine-proximal mutations associate with increased cysteine ligandability. Top, examples of increased cysteine ligandability following the proximal mutations (green) in the indicated cell lines compared to cells expressing the WT protein. Bottom, mutation of Glu110->Lys (green) in IDH1 is associated with an increase Cys379 (red) ligandability (PDB:1T09^118^). (D) Proximal mutations associating with increased cysteine anti-ligandability. Top, examples of increased cysteine anti-igandability following the proximal mutations (green) in the indicated cell lines compared to cells expressing the WT protein. Bottom, mutation of Arg->Gln (green) in ACAA1 increases Cys381 (blue) anti-ligandability (PDB:2IIK). (E) Defining ligandable transcription factor dependencies. Plot comparing differential essentiality, expression and cysteine ligandability in each lineage profiled in DrugMap. (F) The DNA binding domain of NFkB1 (white) from PDB:2O61^119^ bound to DNA (blue). The Cys61 alpha carbons (red) lie adjacent to the DNA backbone. (G) Identification of a covalent ligand for NFkB1•C61. iso-TMT screen of 4,000+ cysteine reactive compounds identified SH-7346, which ligands NFkB1•C61 with good specificity. Insets, iso-TMT engagement profile for SH-7346 and SH-6486. Bottom, further development of SH-7346 to SH-1696 was guided by NFkB1•C61 engagement as determined by intact mass spectrometry (see also Figure S7A-B and Methods). (H) Covalently bound SH-9857 blocks NFkB1 DNA binding. The SH-9857 (an SH-7346 analog)-bound structure resolved at 2.02Å was aligned. The proximity of the Cys61 alpha carbons results in SH-9857 sterically clashing with DNA (See also **Figure S7C, Table S15**). (I) SH-1696 engages NFkB1 in a Cys61-specific manner. Intact mass spectrometry analysis of SH-1696 and SH-9791 (negative control) binding to recombinant NFkB1. (J) SH-1696 disrupts NFkB1-DNA interactions. Cellular lysates isolated from HEK-293FT expressing HiBiT-tagged NFkB1•WT or NFkB1•C61S were treated with SH-1696 (3 nM-20 µM). IC_50_ was determined by measuring disruption in DNA binding through NFkB1 enrichment with a biotinylated oligonucleotide (see 8Methods). (K) SH-1696 blocks NFkB1 transcriptional activity. HEK-293 cells expressing a NFkB1 transcriptional reporter were treated with SH-1696 or SH-9791 (10 µM) and relative transcriptional activity was determined 3 hrs post-treatment. (L) SH-1696 engages NFkB1•C61. MM1S cell lysate was treated with 50 µM of SH-1696 or SH-9791, and NFkB1•C61 engagement was determined by iso-TMT (see also **Table S8**). (L) SH-1696 downregulates NFkB1 target genes in haematopoietic cancers. The indicated NFkB1-dependent cell lines were treated with vehicle (DMSO), SH-1696, or SH-9791 (10 µM) for 3 hrs, and relative gene expression was determined by qPCR (see also **Figure S7E**). Data are represented as mean ± SD. *p< 0.05. Student’s *t*-test (two-tailed, unpaired) were used to determine statistical significance.

Given these genetic contributions to cysteine ligandability, we leveraged gene essentiality data to identify differentially targetable and essential proteins. To prioritize ligand development, we focused on transcription factors, because this class of proteins, as a whole, displays the greatest lineage-restricted essentiality^79^ (**Figure S6G**). Interestingly, multiple lineage-restricted transcription factors required for proliferation were found to contain highly ligandable cysteines (**Figure 3E**). Prominent examples included SOX10•C71 and PAX8•C45/C57 in solid tumors, and IRF4•C194 and NFkB1•C61 in heme malignancies. Given their high degree of lineage-specificity, if these proteins could be targeted by covalent agents, they might achieve a large therapeutic index, due to their limited importance in non-essential tissues.

### Disruption of NFkB1-DNA interactions with a cysteine-directed chemical probe

One of the prominent examples identified above, NFkB1, is a compelling ligandable heme transcription factor, given the location of its ligandable cysteine in the DNA-binding interface and strong literature precedent indicating that NFkB1•C61 liganding disrupts the interaction of NFkB1 with DNA^81, 82^ and promotes target degradation^83^ (**Figures 3E-F**). To identify a covalent ligand for NFkB1•C61, we carried out an iso-TMT screen of a library of 4,000+ commercially available cysteine-reactive small-molecule inhibitors. This analysis led to the identification of SH-7346, a cyclic cyanamide that engages NFkB1•C61 with high specificity (**Figure 3G**). We established that SH-7346 engages NFkB1•C61 but not the NFkB1•C61S mutant (**Figure S7A**). We then proceeded to measure the occupancy of a set of pyrrolidine-cyanamides with the goal of improving occupancy through modification of the distal sulfonamide group, using intact mass spectrometry analysis (**Figure S7B**). Compared to the initial hit, we observed an increase in NFkB1•C61 occupancy from 29% to 74% with SH-1696, which has an aniline-substituted pyridyl-sulfonamide group (**Figure S7B**). In the process of hit expansion, we solved the structure of NFkB1 bound to a closely related analog SH-9857 at a resolution of 2.02Å (**Figures 3H, S7C**). Alignment of the co-structure of SH-9857-bound NFkB1 with a model of DNA-bound NFkB1 demonstrated that SH-9857 sterically clashes with DNA (**Figure 3H**). We established that SH-1696 disrupts NFkB1-DNA interactions with an IC_50_ of ∼5 µM in a C61-dependent manner (**Figure 3J**) and also blocks NFkB1 transcriptional activity with an IC_50_ of 4.9 µM in a reporter cell line (**Figures 3J, S7D**). SH-1696, but not the structurally related control analog SH-9791, efficiently engages NFkB1 (**Figures 3I, L**, **S7B, Table S8**), with SH-9791 showing little effect on DNA binding or on NFkB1 transcriptional activity (**Figures 3J, 3K-L**, **Table S8**). Finally, we examined SH-1696 activity in five hematopoietic cell lines dependent on NFkB1 (**Figure S7F**), finding that the compound decreases NFkB1 target gene expression as measured by qPCR (**Figures 3M, S7E**). These results suggest that liganding NFkB1•C61 may provide a mechanism to target NFkB1 signaling in hematopoietic cancers.

### Development of a chemical probe that disrupts SOX10 transcriptional activity

We next asked whether we could leverage DrugMap to help guide the development of a chemical probe for lineage-restricted transcription factors that do not have literature precedent. We prioritized the SOX10•C71 ligandability hotspot identified above for this analysis. The SOX10 transcription factor is a member of the SOX family of high mobility group box (HMG-box) transcription factors ^84–86^ and a major known dependency of melanoma cells^87, 88^. We found that SOX10•C71 was highly and consistently ligandable in the majority of melanoma models that we characterized (**Figure 4A**). Furthermore, in melanomas^89^ defined by a high SOX10 signature, including immunotherapy-resistant models, we found that depletion of SOX10 strongly blocked proliferation (**Figure S8A-C**). We observed that SOX10•C71 is localized to the protein’s conserved SOXE dimerization domain^90, 91^ and is predicted to be adjacent to a small-molecule binding groove (**Figure 4B**). In light of these facts, we reasoned that SOX10•C71 represents a highly compelling ligandable dependency to prioritize for inhibitor development.

**Figure 4:**
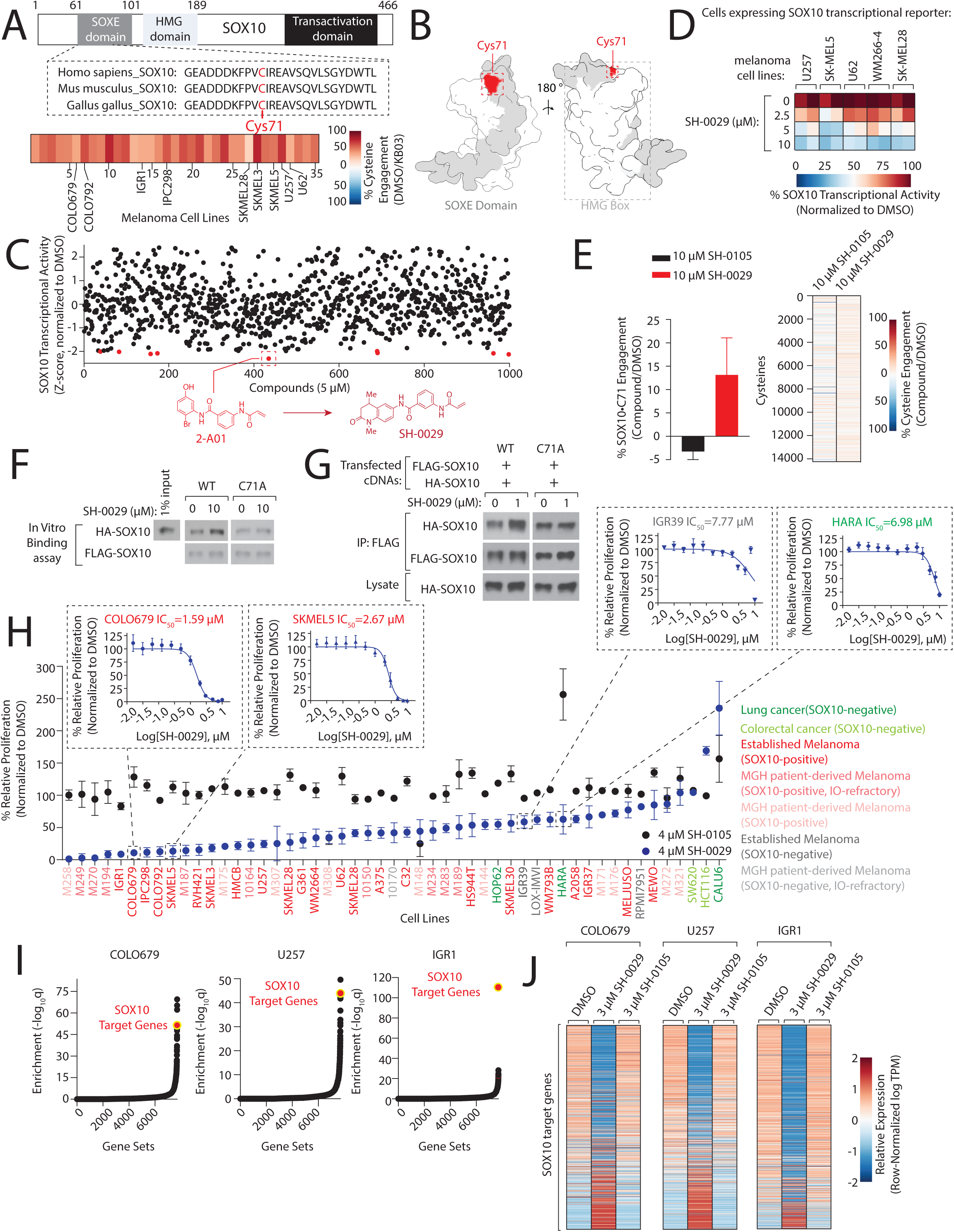
Development of a covalent molecular glue that disrupts SOX10 activity in melanoma. (A) SOX10•C71 is highly ligandable across melanoma cell lines. Top, SOX10•C71 is localized to the SOXE dimerization domain and conserved across vertebrates. Bottom, SOX10•C71 engagement across 35 melanoma cell lines was determined by iso-TMT. (B) SOX10•C71 (red) resides near a putative small-molecule binding groove (opaque red). SOX10 structure (AF-P56693-F1) encompassing amino acids 62-172. (C) Identification of cysteine-reactive compounds that disrupt SOX10 transcriptional activity in melanoma. Top, U257 cells expressing a SOX10 transcriptional reporter were treated with a library of 1000 cysteine-reactive compounds and SOX10 activity was assessed after 48 hrs. Bottom, advancement of hit compound 2-A01 to lead compound SH-0029 was assessed by SOX10 transcriptional activity (see also Figures S9A-B, methods). (D) SH-0029 disrupts SOX10 activity in multiple melanoma cell lines. Melanoma cell lines expressing the SOX10 transcriptional reporter were treated for 48 hrs at the indicated concentrations of SH-0029. (E) SH-0029 engages SOX10•C71 at low occupancy. Left, SKMEL5 lysates were treated with 10 µM SH-0029 or the control compound SH-0105, and SOX10•C71 engagement was determined by iso-TMT. Right, heatmap displaying cysteinome-wide reactivity changes (see also **Figure S9G**, **Table S9**). (F) SH-0029 increases SOX10-SOX10 interactions in a Cys71-dependent manner. Lysates from HEK-293T cells expressing HA-SOX10 or HA-SOX10•C71A were treated with vehicle control or 10 µM SH-0029 and following immunoprecipitation with immobilized FLAG-SOX10 or FLAG-SOX10•C71A, the interaction with HA-SOX10 was determined by immunoblot (see also **Figures S9H-J**). (G) SH-0029 functions as a SOX10 covalent molecular glue in cells. HEK-293T cells expressing the indicated proteins were treated with SH-0029 for 3 hrs and SOX10 dimerization was determined by immunoblot following immunoprecipitation from lysates with anti-FLAG M2 beads (see also **Figures S9K-L**). (H) SH-0029 differentially impacts SOX10-expressing melanoma proliferation. Cell lines were treated with 4 µM of SH-0029 or SH-0105 and proliferation was determined by crystal violet staining following four days of treatment and normalized to vehicle (DMSO). Inset, IC_50_ values were determined in COLO679, SKMEL5, IGR39, and HARA treated with SH-0029 (0.0156-10 µM) (see also **Figure S10B**). (I-J) SH-0029 disrupts SOX10 transcriptional signaling in melanomas. (I) GSEA analysis identifies the SOX10 regulon as highly enriched in melanomas following treatment with SH-0029. (J) Heatmaps displaying relative levels of SOX10-regulated genes following treatment with vehicle, SH-0029 or SH-0105 were determined by RNAseq (see also **Figure S10E**).

To develop a covalent probe for SOX10•C71, we established a SOX10 transcriptional reporter assay that was responsive to SOX10 depletion in the U257 melanoma cell line (**Figure S9A**). Using this assay, we screened a commercially available library of 1,000 cysteine-reactive compounds (**Figure 4C**). Validation of the hits from this screen focused our attention on compound 2-A01, a benzanilide acrylamide, and we subsequently identified additional analogs with increasing potency, including SH-0029, a benzamide-THQ acrylamide (**Figure S9B**). We found that SH-0029 disrupts SOX10 transcriptional readout in five melanoma cell lines, with an IC_50_ of ∼11 µM, but a related analog, SH-0105 had minimal activity (**Figures 4D, S9B-C**). Additionally, SH-0029 did not perturb the transcriptional activity of other lineage-restricted transcription factors (**Figure S9D**). Using a biotinylated version of the SOX10 ligand (SH-0029-DTB), we verified that it specifically engages SOX10•C71 but not a SOX10•C71A mutant, in HEK-293T cells (**Figure S9E**). SH-0029 outcompeted SH-0029-DTB labeling of SOX10 in two melanoma cell lines, providing further evidence of SOX10•C71 engagement (**Figure S9F**). Initially, we were perplexed by the strong disruption of transcriptional activity but low level of SOX10•C71 engagement in iso-TMT experiments (**Figures 4E, S9G, Table S9**). This surprising finding suggested an alternative mechanism of inhibition other than one that requires high-occupancy binding^92–95^. The localization of SOX10•C71 to the SOXE domain, which has been implicated in SOX10 dimerization^96^, led us to explore whether SH-0029 might modulate SOX10 multimerization. A SOX10 *in vitro* binding assay revealed that SH-0029, rather than disrupting SOX10-SOX10 dimerization, instead increases the interaction. This result was recapitulated in cells (**Figures 4F-G, S9H-I**). This ‘glue-like’ effect of SH-0029 was completely dependent on Cys71, and SH-0029 did not increase the binding in a SOX10•C71A mutant, either *in vitro* or in cells. This result was echoed by the negative control probe, SH-0105 (**Figures 4F-G, S9I-K**).

### A covalent SOX10 ligand disrupts melanoma transcriptional signaling and proliferation

Given the strong and consistent proliferation defect induced by depletion of SOX10 in melanoma cell lines (**Figures S8A-B**), we proceeded to examine the activity of SH-0029 in 40+ melanoma cell lines of diverse origins. This revealed a wide range of inhibition of proliferation (**Figures 4H, S10A)**. Sensitivity to SH-0029 correlated with an established SOX10 transcriptional signature and with sensitivity to SOX10 depletion (**Figure S10B-C)**. Expression of a SOX10•C71A mutant in U257 melanoma cells partially, albeit significantly, rescued the proliferation block of SH-0029 in comparison to cells expressing WT SOX10 (**Figures S10D**). In contrast, treatment with SH-0029 or depletion of SOX10 showed only modest activity in SOX10-negative melanomas and in lung or colon models deficient in SOX10 expression (**Figures 4H, S10A-B**).

To investigate how liganding of SOX10 impacts melanoma transcriptional networks, we defined an extended SOX10 transcriptional signature by depleting SOX10 in seven melanoma cell lines and analyzing the resulting transcriptional changes by RNAseq (**Figure S10E, Table S10**). SH-0029 treatment in three melanoma cell lines revealed a significant enrichment for the SOX10 target genes as determined by gene set enrichment analysis, which we confirmed in other melanomas by qPCR (**Figures 4I-J, S10F-G, Table S11**). In SK-MEL5 cells, there was a strong and consistent overlap between cellular pathways perturbed following SH-0029 treatment and SOX10 depletion, and we found that many of the genes downregulated by SH-0029 in this model were bound by SOX10 near active enhancers, including those of the genes ERBB3 and IL-16 (**Figures S10H-J, Table S12**). ERBB3 is a known melanoma-specific dependency^97^ and IL-16, an immunomodulatory regulator, is preferentially expressed in SOX10-high melanocytic cells^98^, suggesting that the SOX10 ligand causes down-regulation of the SOX10 transcriptional network. Collectively, our findings demonstrate the unexpected ligandability of SOX10, and show that its covalent adduction can disrupt both oncogenic transcriptional signaling as well as proliferation in melanomas. More generally, proven by examples like this, DrugMap provides a detailed roadmap for developing covalent ligands against challenging targets in oncology, by integrating high-throughput dependency and ligandability datasets.

## Discussion

Large-scale systematic analysis of cancer genomes, proteomes and metabolomes has been leveraged to nominate targets for drug development^33, 79, 99^. DrugMap represents the chemical-biology complement to these studies, providing a roadmap for systematically evaluating cysteine ligandability across cancers.

Perhaps one of the most surprising findings from this study is that cysteine ligandability is heterogeneous. Although heterogeneity is now established as the rule rather the exception in cancer^26^, this principle had yet to be extended to protein-ligand interactions. It is likely to have important ramifications for understanding differential responses to cysteine-reactive drugs in the clinic. Our finding that the cellular redox state is an important determinant of ligandability accords well with previous studies demonstrating that covalent inhibitors targeting EGFR can be modulated in their efficacy through active-site cysteine sulfenylation^100–103^. As multi-omic profiling of tumors continues to identify new cell states, our data raise the intriguing possibility that this information can be leveraged to target proteins within cancers defined by a specific cell state, in addition to targets characterized by mutation or differential expression. Future studies aimed at deciphering how different metabolic states can regulate cysteine ligandability will be critical both for designing new therapeutic strategies and for understanding which tumors may or may not be amenable to specific covalent drugging approaches.

Our investigations of the molecular features underlying heterogeneity in cysteine ligandability reveal a prominent role for protein conformational changes. When viewed through the lens of protein complexes, this heterogeneity may reflect differences in protein conformations or complexation across cancer cell lines^74^. While the systematic evaluation of protein conformations has historically been restricted to purified recombinant proteins, new mass-spectrometry based assessments of protein conformations are beginning to reveal differences in protein-metabolite interactions and protein-protein interactions in their native contexts^30, 104–108^. However, these studies have been restricted to a handful of cell lines^109^. In contrast, the comprehensive data obtained herein begins to define a picture in which protein conformations may be substantially altered across a large number of cancer lineages, thus providing a novel opportunity to understand how cell state governs protein dynamics.

Drug-discovery approaches are assisted profoundly by protein structure determination^57, 105, 110^. However, for many targets of interest, including transcription factors, structural solutions are not currently possible because of the intrinsically disordered regions within these proteins. Cysteine-focused chemical proteomics offers one potential solution to this problem, by systematically mapping sites of ligandability with covalent inhibitors. Our discovery that cysteine *anti-ligandability* can serve as a proxy readout of protein conformational changes suggests that this approach, when fully realized, may enable detection of cryptic or transient pockets in complex mixtures of protein states. In support of this hypothesis, we measured changes in cysteine accessibility following engagement with drugs directed against EGFR and XPO1 and proceeded to identify an increase in ligandabilty among these structurally-sensitive cysteines. Importantly, a change in protein conformation as determined by proteomics may also provide a first indication that liganding of a given cysteine has a measurable, functional impact on a protein, which may be a useful triaging mechanism for cysteines lacking annotated functions.

Cancer is a genetic disease, and tumors contain a multitude of mutations of unknown significance. While a loss-of-function mutation in a well-annotated protein domain provides important clues about its potential impact, for the vast majority of missense mutations, the functional consequences are unknown. Our study suggests that monitoring small-molecule interactions with proteins may help to reveal the functional impact of specific missense mutations. By leveraging the deep genomic characterization that has been performed for the majority of cell lines profiled in DrugMap, we begin to uncover how mutations can influence cysteine ligandability, providing not only opportunities for mutation-specific targeting but also a novel potential readout of protein conformational dynamics as read out by protein-ligand interactions in unique mutational contexts. Perhaps the biggest surprise from this analysis is the pervasiveness of cysteine-proximal mutations that associate with altered cysteine ligandability. This finding suggests that many more opportunities exist for cancer-specific targeting in addition to concentrating on *de novo* cysteine mutations like KRAS•G12C^18, 33, 99^. Further understanding how mutations in *trans* affect cysteine ligandability is an important future endeavor that can be aided by studies that globally assess protein-protein interactions in cancer using co-expression as an indirect readout of these connections^111, 112^.

Transcription factors represent ∼19% of all oncogenes and, as a class, show the greatest differential essentiality in pan-cancer functional genomic investigations^113^. Despite this, transcription factors have historically been thought to lie largely outside the reach of small-molecule manipulation. In a hopeful development, this study reveals that multiple transcription factors in fact contain highly ligandable cysteines, and this information can be usefully leveraged, which we prove by developing chemical probes to manipulate the activity of NFkB1 and SOX10. The development of each probe was entirely dependent on information about the target cysteine made available by the DrugMap atlas. Based on the localization of NFkB1•C61 to the DNA-binding site, this protein provides a prominent example of how cysteine liganding can disrupt this biological activity, especially when realized with a more advanced inhibitor compound. For SOX10, a functional screen was required which led to the unexpected identification of a covalent ‘glue-like’ mechanism like that previously reported for covalent ligands targeting TP53^114^. Given that numerous components of the transcriptional machinery function through multimerization, it would not be surprising if many other members of this class of proteins will be similarly amenable to this form of disruption, with covalent attachment proving to be the key to stabilizing transient interactions.

In summary, this study highlights critical insights provided by cysteine-focused chemical proteomics, and systematically interrogates the general principles governing cysteine ligandability across cancer. While understanding what is ligandable in cancer is an immediate output of this study, it is also important to emphasize that chemical proteomics is at its core a distinct measure of protein-ligand interactions, and hence protein conformational states. Going forward, a broader characterization of cysteine ligandability in patient tumors is likely to further reveal how cysteine targeting can be influenced by the tumor and biochemical microenvironments, and to further illuminate the vast heterogeneity emerging from genomic studies of these malignancies.

## Acknowledgements

We thank all members of the Bar-Peled and Lawrence Labs, David Sabatini and Keith Flaherty for helpful suggestions. This work was supported by the Damon Runyon Cancer Research Foundation (62-20), the American Association for Cancer Research (19-20-45-BARP), the American Cancer Society, the Melanoma Research Alliance, the Ludwig Cancer Center of Harvard Medical School, Lungevity, ALK Positive, V-Foundation, Mary Kay Foundation, Paula and Rodger Riney Foundation, the PEW-Stewart Trusts, Lisa and Mark Schwartz and the NIH/NCI (1R21CA226082-01, R37CA260062) DF/HCC SPORE in Gastrointestinal Cancer, NIH/NCI (P50CA127003).

## Author contributions

M.T., H.B.C, S.Z., M.L. and L.B-P. conceived and designed the study. M.T. performed most of the experiments with the assistance of S.Z., M.J.L., M.M., M.G., M.G., J.Z., J.A. F.M., K.V. M.R., H.P., H.G., T.H., S.O., I.R., and D.E.F. H.B.C. and M.L. performed bioinformatics analysis and protein structural analysis with the assistance of G.K., B.H., N.B., and G.P. S.Z. performed proteomics analysis and interpreted the data with assistance from C.-C.T., G.K., B.M., T.-y.Y., N.K., P.-C.T., and N.C. S.H., R.W., and B.L. synthesized compounds and analyzed with LC/MS and NMR. M.T. with assistance from T.H., L.B., and M.L.S. prepared samples for SmartSeq2. A.S.O.A., K.R., A.V., and M.J.R performed RNAseq analysis and ChIP-seq sample preparation and analysis. D.H., A.-S.K., S.O., B.K., L.S., M.Q.R., A.C.D., A.L., J.F., S.Y., L.P.N., S.M.R., F.N.W., A.A., S.E.C., T.D., S.M., D.A.H., G.B., M.S.-F., R.J. A.H., N.B., M.L.S., and C.O. generated the models underlying DrugMap. B.R.D. assisted with data visualization. M.T., H.B.C, S.Z., M.L. and L.B-P. wrote the manuscript with assistance from all the coauthors. M.L. and L.B-P. supervised the studies.

---

**Figure S1:**
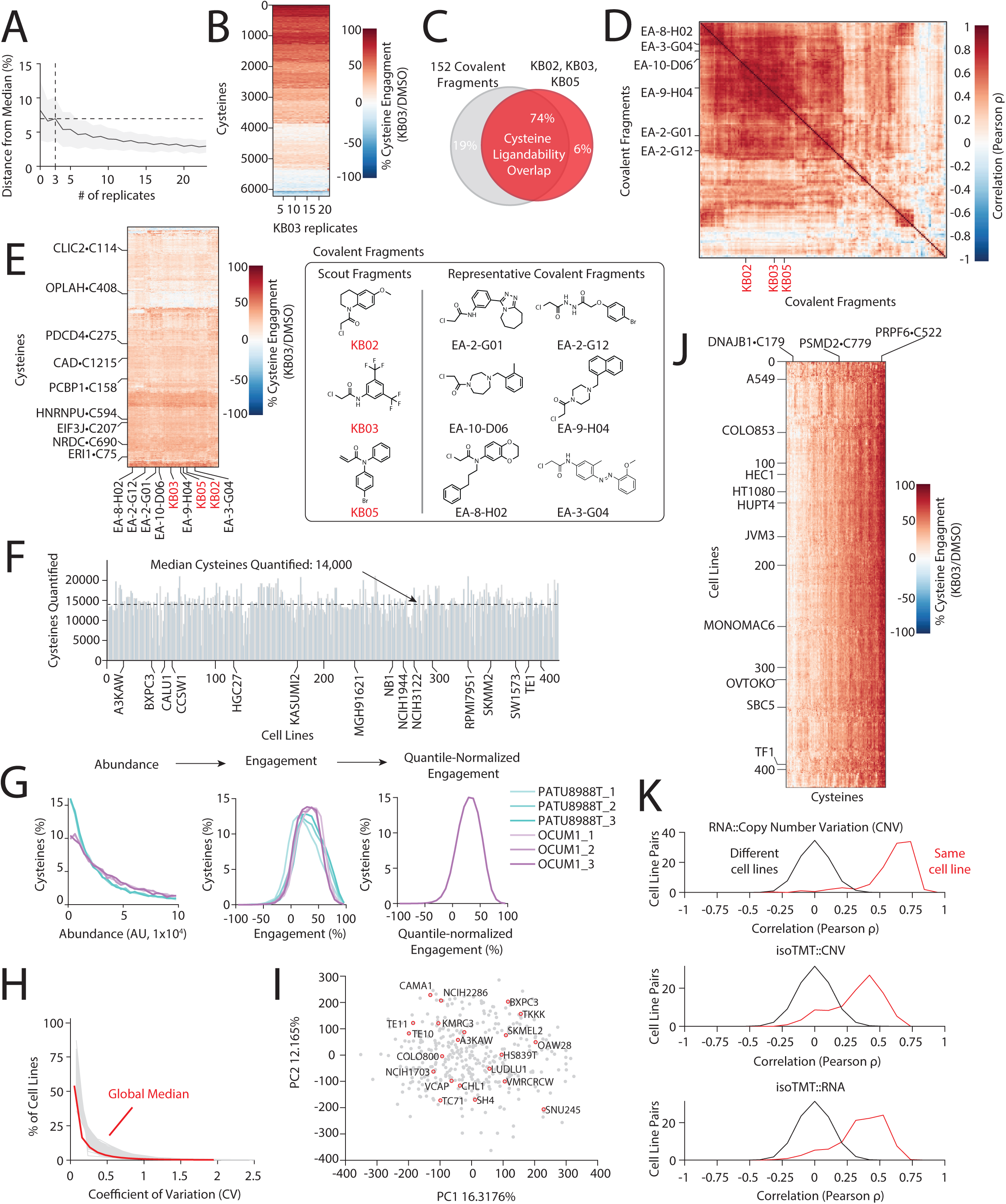
Development of a Pan-Cancer cysteine ligandability map, related to Figure 1. (A) Saturation analysis demonstrates that three replicates are sufficient to quantify engagement within 2% of the median cysteine engagement across 24 replicates of K562 cells. K562 cell lysates were treated with 500 µM KB03 or vehicle, and changes in cysteine ligandability were determined by iso-TMT (see Methods). (B) Heatmap of cysteine engagement across >6000 cysteines detected across 24 replicates. Cysteine ligandability was determined as described in (A). (C) The scout fragments KB02, KB03 and KB05 broadly recapitulate cysteine engagement of 152 cysteine-reactive fragments. K562 cell lysate was treated with the indicated compounds (500 µM) and cysteine engagement was determined by iso-TMT. Overlap in cysteine ligandability between scout fragments and 152 cysteine reactive compounds (n=16,026 cysteines analyzed). (D) Hierarchical clustering identifies a subset of compounds having high correlation with KB05, KB03, or KB02. (E) Left, heatmap depicting cysteine reactivity changes following treatment with scout probes or highly correlated fragments. Right, structures of fragments that correlate strongly with scout fragments. (F) Quantification of cysteine-containing peptides for each cell line profiled in DrugMap. (G) Schematic describing data transformation workflow (see also Methods). (H) Low coefficient of variation across replicates of cell lines. (I) Principal component analysis (PCA) of cysteine ligandability across DrugMap demonstrates the presence of modest sources of variation that structure the entire dataset. (J) A subset of cysteines display high reproducibility in engagement across 416 cell lines. (K) Mutual correlation of RNA expression level, DNA copy number variation (CNV), and iso-TMT protein abundance.

**Figure S2:**
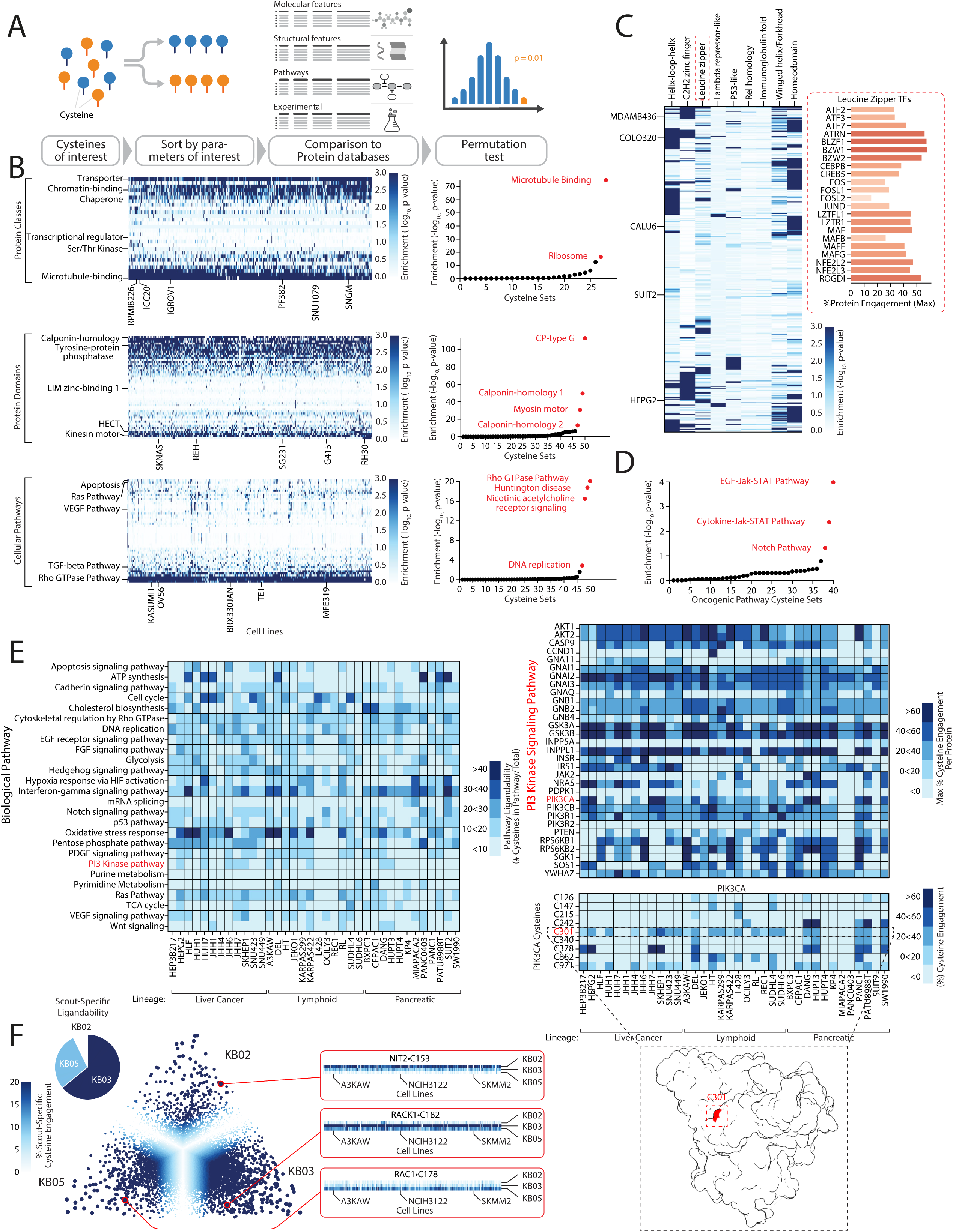
Identification of highly ligandable cysteines in protein domains, classes and pathways, related to Figure 1. (A) Schematic describing Cysteine Set Enrichment Analysis (CSEA), a bioinformatic pipeline to identify enriched cysteine sets defined by molecular, experimental, structural and pathway-level features (see also **Table S16**, Methods). (B) Heatmaps (left) and CSEA enrichment plots (right) displaying highly enriched protein classes, domains, and pathways in each cancer cell line profiled in DrugMap. (C) CSEA identification of ligandable transcription factor classes across DrugMap. Inset, engagement of leucine zipper transcription factors by scout fragments. (D) Ligandable cysteines localize to oncogenic pathways. (E) Pathway/protein/cysteine-level analysis of cysteine ligandability in liver cancer, lymphomas, and pancreatic cancer. Left, Heatmap of cysteine ligandability in each biological pathway. Right, Heatmap of ligandable proteins involved in the PI3K pathway. Bottom, Structure of PIK3CA with ligandable cysteine C301 highlighted in red (PDB:2RD0^120^). (F) Some cysteines display scout-specific liganding. Left, fraction of cysteines uniquely engaged by each scout probe across DrugMap. Right, ternary plot highlighting examples of cysteines preferentially liganded by one scout.

**Figure S3:**
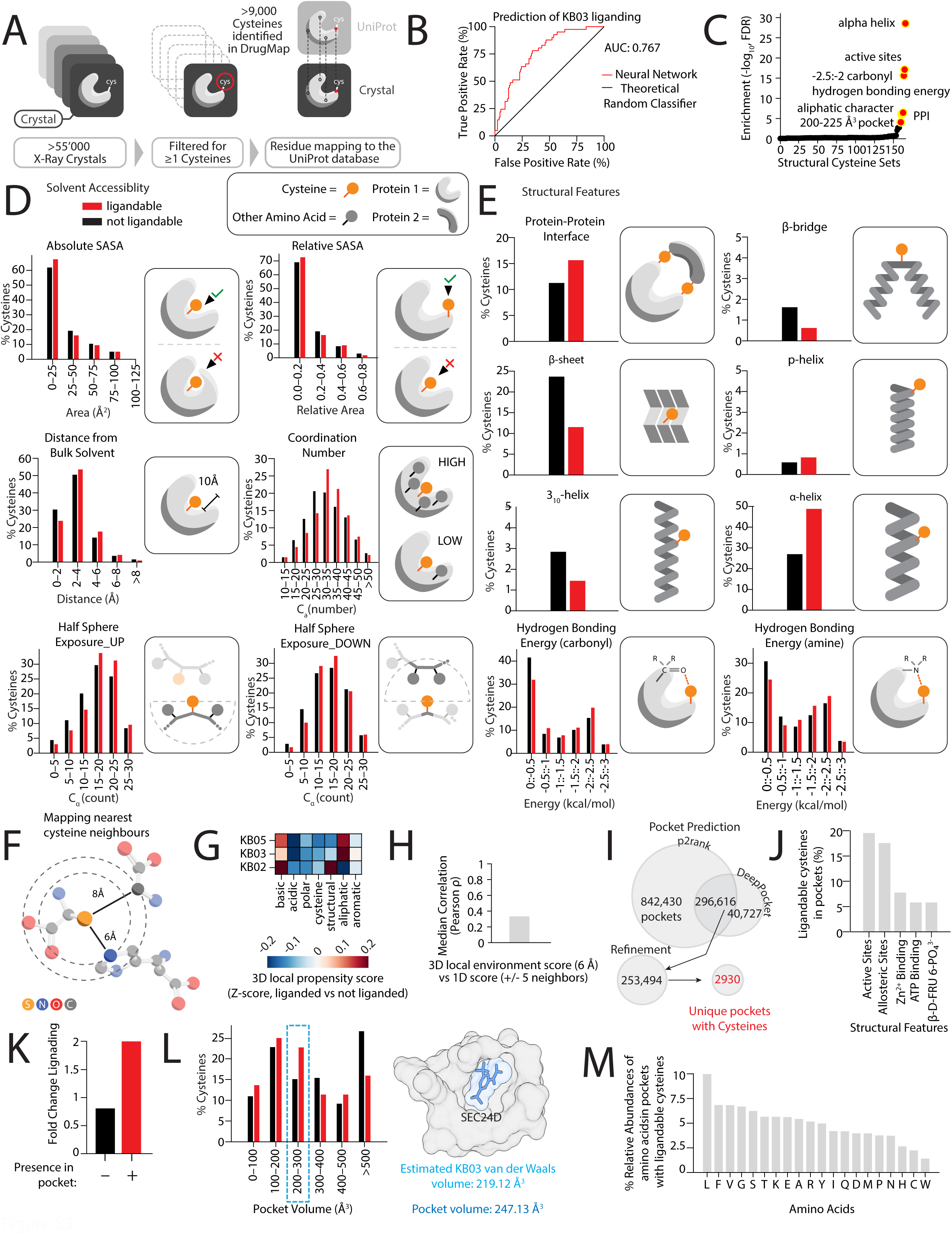
Structural characterization of ligandable cysteines, related to Figure 1. (A) Overview of the cysteine structural annotation that forms the basis for structure-informed CSEA. (B) Structure-informed deep learning enables ∼70% accuracy in predicting KB03 liganding (see Methods). (C) Structural cysteine sets enriched in KB03-liganded cysteines. (D-E) Structural features underlying cysteine ligandability. The indicated structural parameters were calculated for ligandable (red) and non-ligandable (black) cysteines by mapping cysteines identified in DrugMap (n=9,352) to corresponding human PDB structures (n>55,000) and extracting structural parameters (see Methods). (F) Schematic displaying nearest-neighbor identification. (G) Basic amino acid neighbors are enriched near ligandable cysteines, whereas acidic amino acids are depleted. (H) Bar chart showing disagreement between primary-sequence neighbors and 3D neighbors of ligandable cysteines. (I) Overlap in annotated pockets using p2rank^60^ and DeepPocket ^58^. (J) Ligandable cysteines (ε>60%) are more likely to reside in pockets than not. (K) KB03-engaged cysteines are enriched in pockets having volumes of 200-300 Å^3^. Right, distribution of ligandable cysteines across pockets of different volumes. Left, predicted KB03 van der Waals volume (blue) and docking within a pocket identified in protein SEC24D (PDB=3EFO^121^, see Methods). (L) Ligandable cysteines within pockets are localized to the indicated protein domains. (M) Amino acid content of protein pockets containing ligandable cysteines (see Methods).

**Figure S4:**
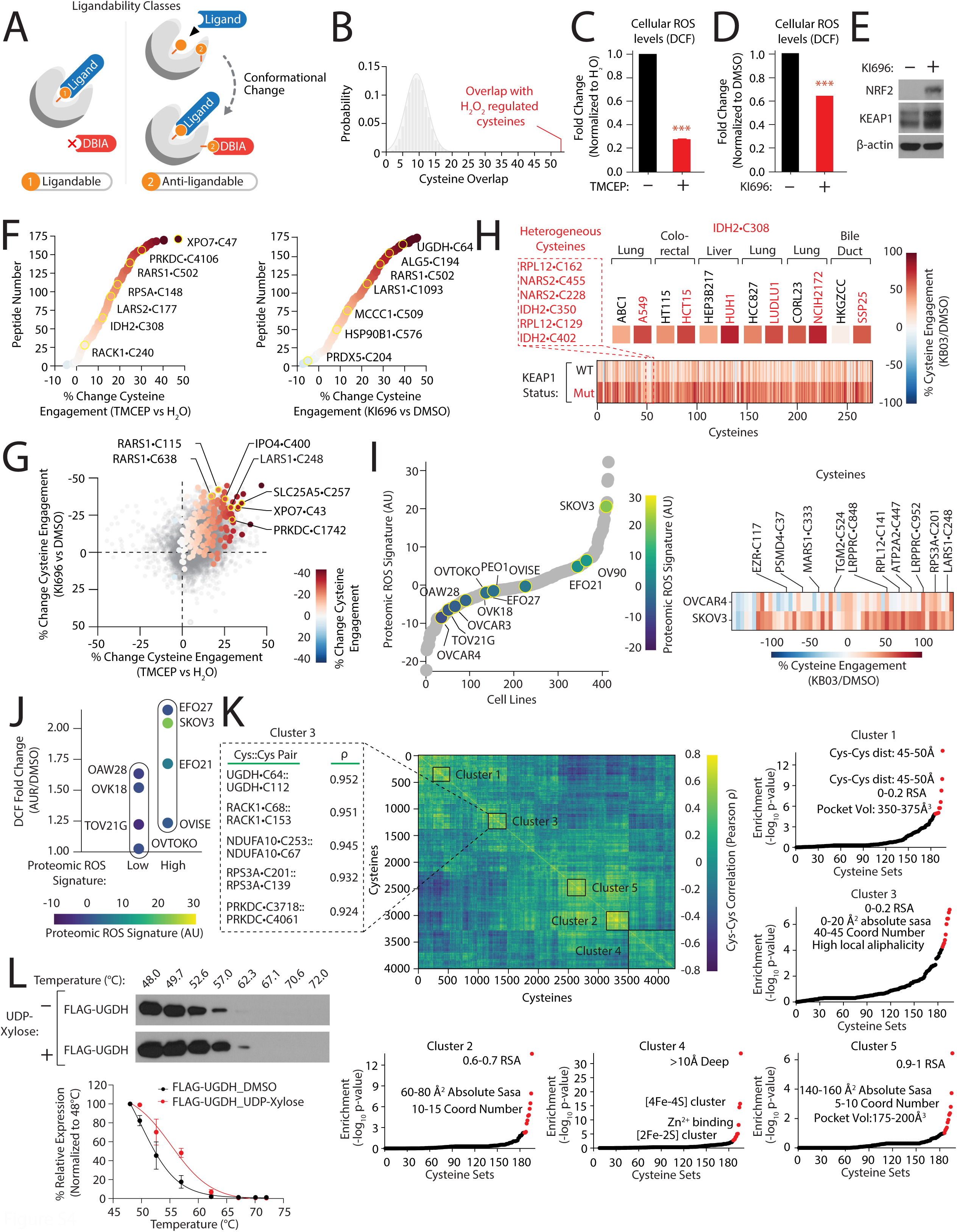
Cellular redox state and protein conformational changes regulate cysteine ligandability, related to Figure 2. (A) Schematic representation of different cysteine classes discovered in DrugMap. (B) Overlap of heterogeneous cysteines with H_2_O_2_-regulated cysteine set as determined by CSEA. (C) TMCEP induces a reductive state in K562 cells. Flow cytometry analysis was used to monitor changes in DCF fluorescence in K562 cells treated with 1 mM TMCEP for 1 hr. (D) NRF2 activation regulates cellular redox states. K562 cells were treated with 1 µM KI696 (NRF2 activator) for 48 hrs, and changes in DCF fluorescence were determined as described in (C). (E) Immunoblot of the indicated proteins in K562 cells following treatment with vehicle (DMSO) or 1 µM KI696 for 48 hrs. (F) Impact of redox environment on heterogeneous cysteine ligandability. Scatter plot denoting changes in the ligandability of heterogeneous cysteines following TMCEP treatment (left) or NRF2 activation (right). (G) TMCEP treatment and NRF2 activation increase a subset of heterogeneous cysteine ligandability. Scatter plot comparing changes in cysteine ligandability in K562 cell lysate following the indicated treatments (see also Figures 2D-E). Heterogeneous cysteines are highlighted. (H) A subset of heterogeneous cysteines are differentially ligandable in KEAP1-mutant cells. Bottom, heatmap depicting differential cysteine engagement between KEAP1-mutant and KEAP1-WT cancer cell lines profiled in DrugMap. Heterogeneous cysteines are indicated. Top, representative examples of differential ligandability for IDH2•C308 in a panel of KEAP1-mutant (red) and KEAP1-WT (black) cell lines of various lineages. (I) Cysteine ligandability is an approximate measure of cellular redox state. Left, plot of proteomic redox score for 416 cell lines based on the ligandability of redox-sensitive heterogeneous cysteines. Right, ligandability of heterogeneous redox-sensitive cysteines in OVCAR4 and SKOV3 ovarian cancer cell lines (see Methods). (J) Proteomic redox score correlates with cellular response to auranofin. Comparison of DCF fluorescence changes following auranofin treatment for eight ovarian cancer cell lines with differing proteomic redox scores. The indicated cell lines were treated with 1 µM auranofin (AUR) for 6 hrs, and changes in DCF fluorescence were determined as described in (C). (K) Cys-cys ligandability correlations are enriched for cysteines within close spatial proximity. Left, heatmap depicting correlations of 4,000+ cys-cys pairs. Surrounding insets, enrichment of structural elements by CSEA on the indicated clusters of highly-correlated cysteine pairs. Boxed inset, top-ranked cys-cys correlations identified in Cluster 3. (L) UDP-xylose increases UGDH thermal stability. Top, cell lysate isolated from HEK-293T cells overexpressing FLAG-UGDH was treated with 1 mM of UDP-xylose followed by incubation at the indicated temperatures for 3 min. Following centrifugation, the relative levels of UGDH were determined by immunoblot (see Methods). Bottom, quantification of UGDH thermal stability following treatment with UDP-xylose. Data are represented as mean ± SD. *p < 0.05. ***p < 0.0001. Student’s *t*-test (two-tailed, unpaired) was used to determine statistical significance.

**Figure S5:**
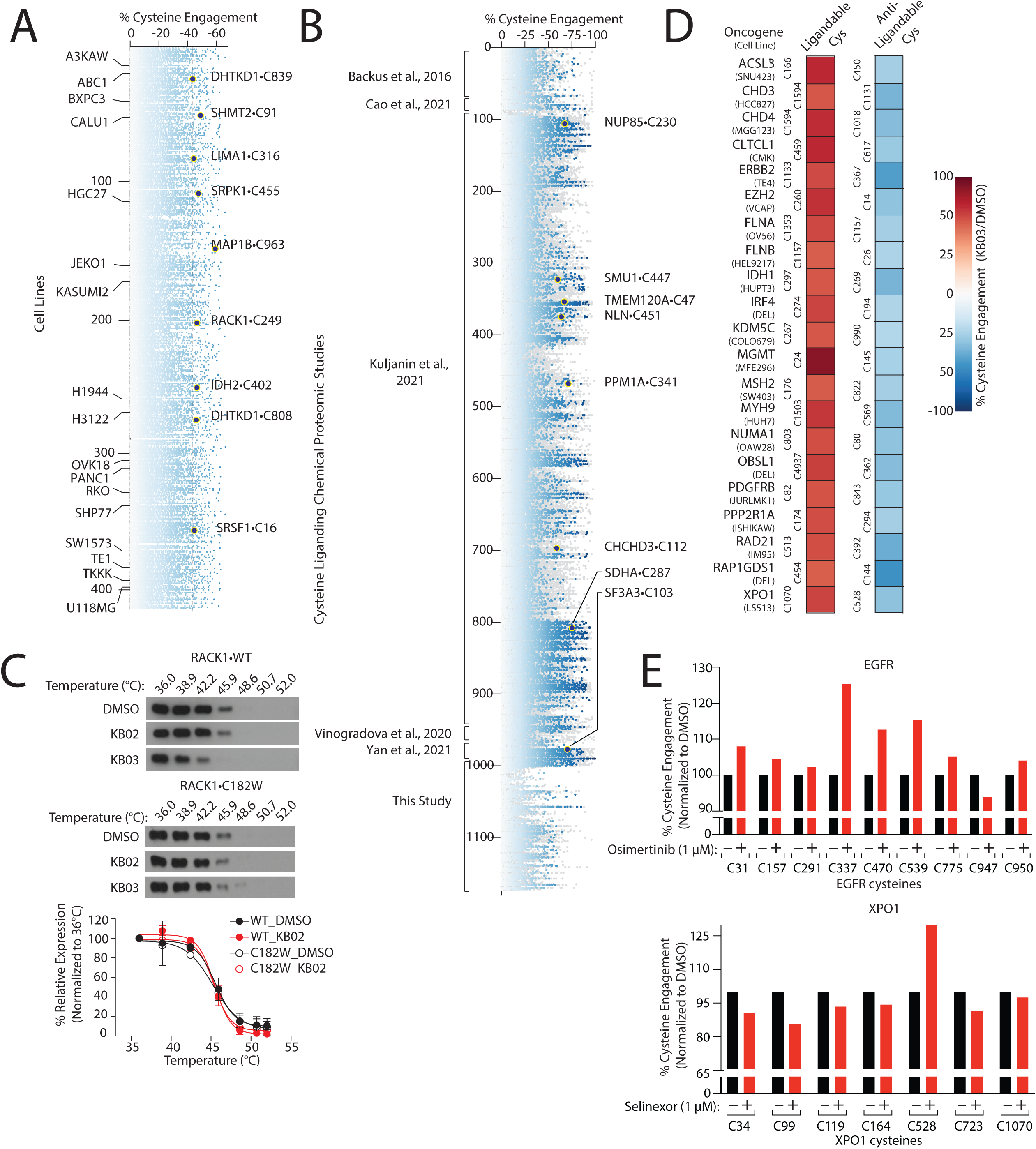
Ligand-induced conformation changes promote greater cysteine accessibility, related to Figure 2. (A) Anti-ligandable cysteines identified in cell lines analyzed in DrugMap. (B) Anti-ligandable cysteines are pervasive in covalent fragment treatment studies. Dot plot highlighting anti-ligandable cysteines in the indicated studies. (C) RACK1•C182 liganding by KB03 leads to thermal instability. Top, cell lysate isolated from HEK-293T cells expressing FLAG-RACK1 or FLAG-RACK1•C182W was treated with DMSO, 100 µM KB02 or 100 µM KB03 for 1 hr followed by incubation at the indicated temperatures for 3 min. Following centrifugation, the relative levels of RACK1 were assessed by immunoblot. Bottom, quantification of thermal stability of RACK1 following treatment with KB02 (see also Figure 2K, Methods). (D) Oncogenes with ligandable and anti-ligandable cysteine pairs identified in DrugMap. (E) Cysteine accessibility of EGFR (top) and XPO1 (bottom) following treatment with the corresponding covalent drug (see also Figure 2L).

**Figure S6:**
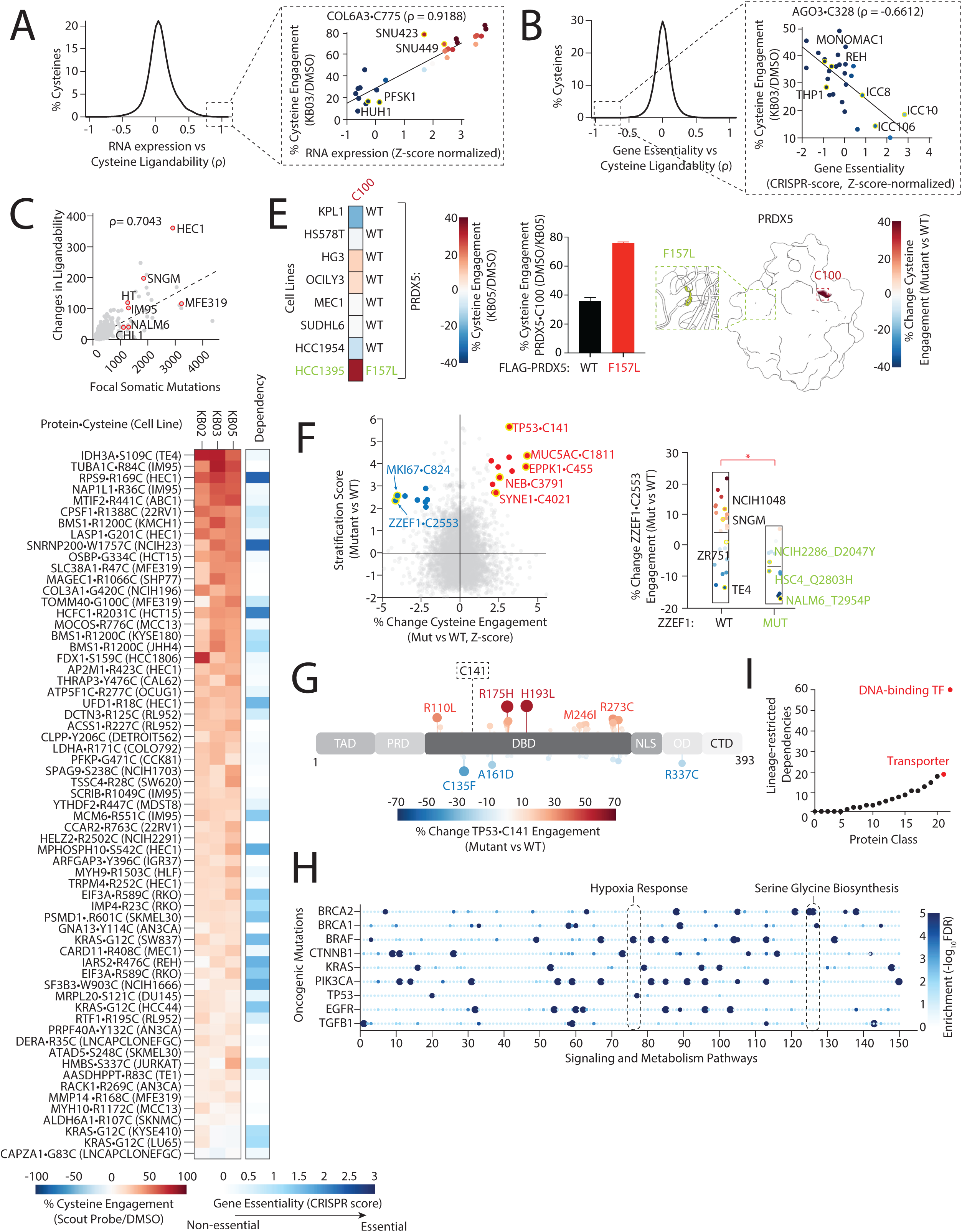
Genetic determinants of cysteine ligandability, related to Figure 3. (A) RNA-expression and cysteine ligandability are poorly correlated in general. Distribution of correlations between cysteine ligandability and RNA expression. Inset, ligandability of COL6A3•C775 is highly correlated with its corresponding RNA expression across DrugMap. (B). Gene essentiality and cysteine ligandability are poorly correlated in general. Distribution of correlations between cysteine ligandability and CRISPR-based genetic dependency. Inset, ligandability of AGO3•C328 is negatively correlated with its corresponding dependency across DrugMap. (C) Increased mutational burden correlates with increased changes in cysteine ligandability. Scatter plot comparing cysteine ligandability and mutational burden for cell lines analyzed in DrugMap. (D) Identification of newly encoded cysteines that are ligandable in DrugMap. Left, heatmap depicting ligandability of newly encoded cysteines in cell lines harboring the corresponding mutations. Right, dependency of the corresponding gene for each protein harboring a *de novo* ligandable cysteine. (E) Mutation of Phe157–>Leu increases PRDX5•C100 ligandability. Left, heatmap showing differential engagement of PRDX5•C100 across several cell lines in DrugMap, including HCC1395 (green), which encodes PRDX5•F157L. Middle, lysates isolated from HEK-293T cells expressing FLAG-PRDX5 or FLAG-PRDX5•F157L were treated with vehicle or KB05, and ligandability was determined by iso-TMT. Right, structure of PRDX5 highlighting location of F157 and change in ligandability of PRDX5•C100 (PDB:1HD2^122^). (F) Systematic identification of amino acid mutations that associate with changes in cysteine ligandability. Left, scatter plot comparing cysteine ligandability and amino acid mutations. Right, distribution of cysteine ligandability changes in ZZEF1-WT and ZZEF1-mutant cell lines. (G) Lollipop plot depicting changes in TP53•C141 engagement in cell lines harboring the indicated mutations. (H) Cellular pathways with ligandable cysteines enriched in cell lines with corresponding oncogenic mutations. (I) Transcription factors (TFs) represent the most lineage-restricted class of dependencies in DepMap. Data are represented as mean ± SD. *p < 0.05. Student’s *t*-test (two-tailed, unpaired) was used to determine statistical significance.

**Figure S7:**
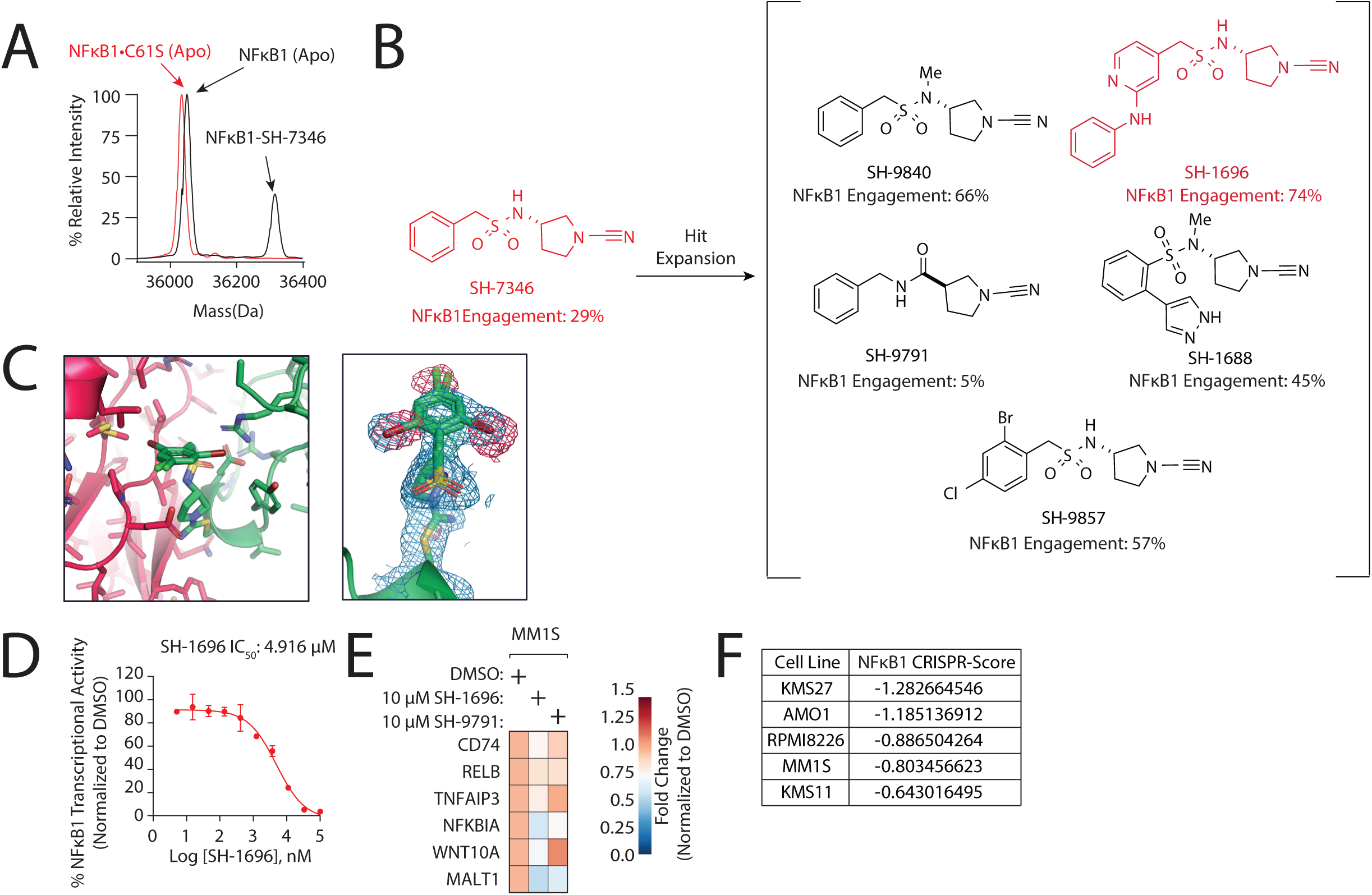
Development of a covalent probe targeting NFkB1•C61, related to Figure 3. (A) SH-7346 ligands NFkB1 in a Cys61-dependent manner. Recombinant NFkB1 or NFkB1•C61S was incubated with 10 µM SH-7346, and engagement was determined by intact mass spectrometry analysis (see Methods). (B) Structures and engagement of SH-7346 and related analogs. NFkB1•C61 engagement with the indicated analogs was measured at 10 µM as described in (A), identifying SH-1696 as an advanced ligand. (C) Conformation and modeling of SH-9857. Left, the SH-9857 bromochlorobenzene ring is modeled in two different conformations that flip the bromine position. Additionally, SH-9857 positions closely between the covalently attached protomer (green) and an adjacent protomer in the crystal lattice (red). Right, the 2Fo-Fc map (1 σ, blue mesh) and Fo-Fc map (*±*3 σ, green and red mesh) around SH-9857. The 2Fo-Fc density and Fo-Fc difference map indicate confident modelling of the compound from the covalent attachment through the sulfonamide. Although the bromochlorobenzene is already modeled in two conformations, compound flexibility or additional unmodeled conformations result in negative density around the halogens. Regardless of this apparent conformational flexibility distal to the covalent bond, DNA binding would still be inhibited (see also Figure 3H, **Table S15**). (D) IC_50_ determination for SH-1696 disruption of NFkB1 transcriptional activity. HEK-293 cells expressing a NFkB1 transcriptional reporter were treated with SH-1696, and relative transcriptional activity was determined 3 hrs post-treatment (see also Figure 3K). (E) SH-1696 downregulates NFkB1 target genes in haematopoietic cancers. The indicated NFkB1-dependent cell lines were treated with vehicle (DMSO), SH-1696, or SH-9791 (10 µM) for 3 hrs, and relative gene expression was determined by qPCR (see also Figure 3L). (F) Dependency of NFkB1 in haematopoietic cancer lines profiled in Figures 3L, **S7E**.

**Figure S8:**
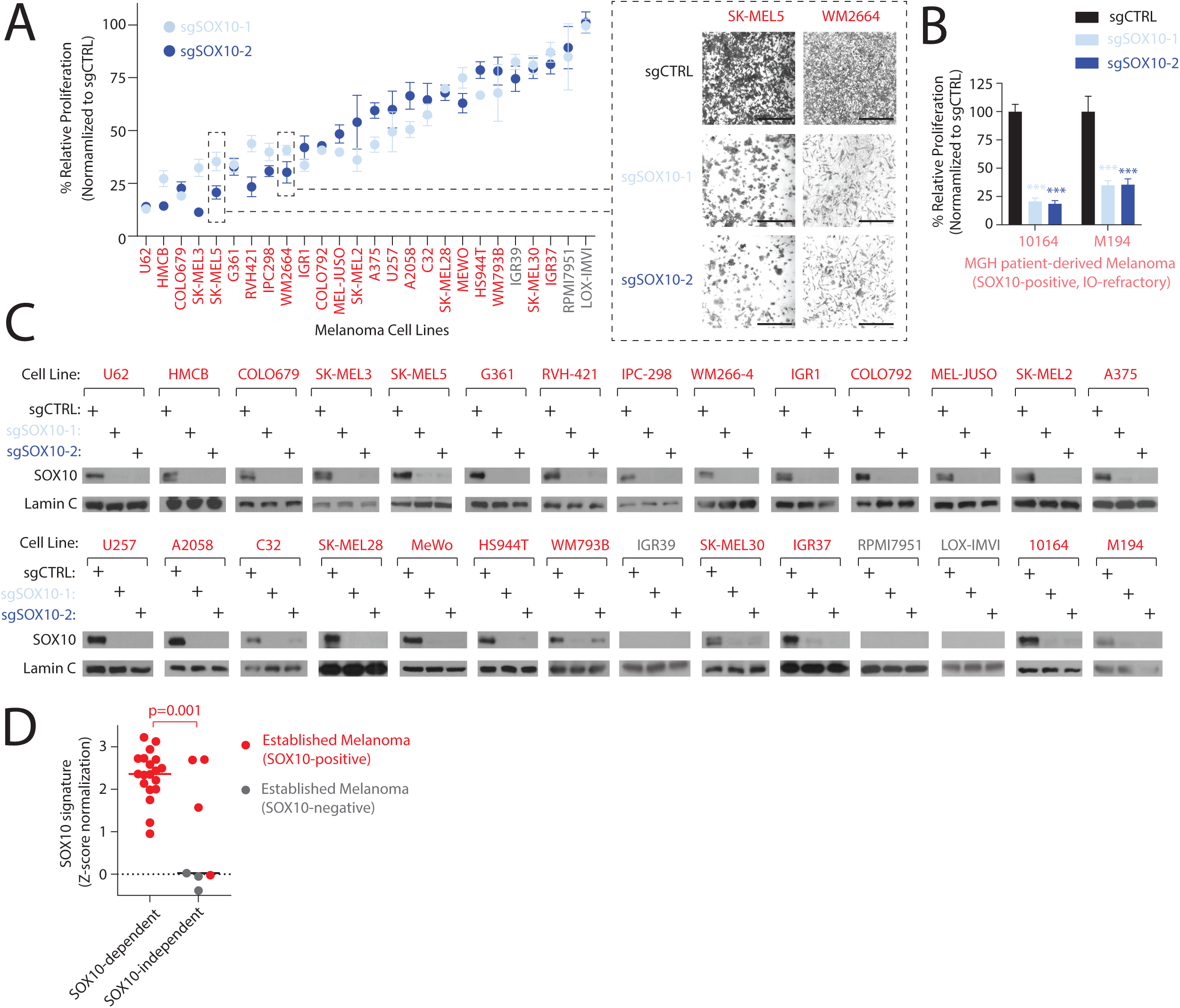
SOX10 is required for melanoma proliferation, related to Figure 4. (A-B) SOX10 is required for the proliferation of established and immunotherapy-resistant melanoma cell lines. Relative proliferation of established melanoma cell lines (A) and immunotherapy-resistant patient-derived lines (B) expressing the indicated sgRNAs was determined by measuring cellular ATP concentrations after six days (A). Cell lines with a decrease in proliferation of >30% following SOX10 depletion are characterized as dependent. Inset, micrographs of SKMEL5 and WM266-4 expressing the indicated sgRNAs. (C) Immunoblot analysis of SOX10 in melanoma cell lines expressing the indicated sgRNAs. (D) SOX10 dependency correlates with SOX10 transcriptional activity (see Methods). Scale bar=200 µm. Data are represented as mean ± SD. ***p< 0.0001. Student’s *t*-test (two-tailed, unpaired) was used to determine statistical significance.

**Figure S9:**
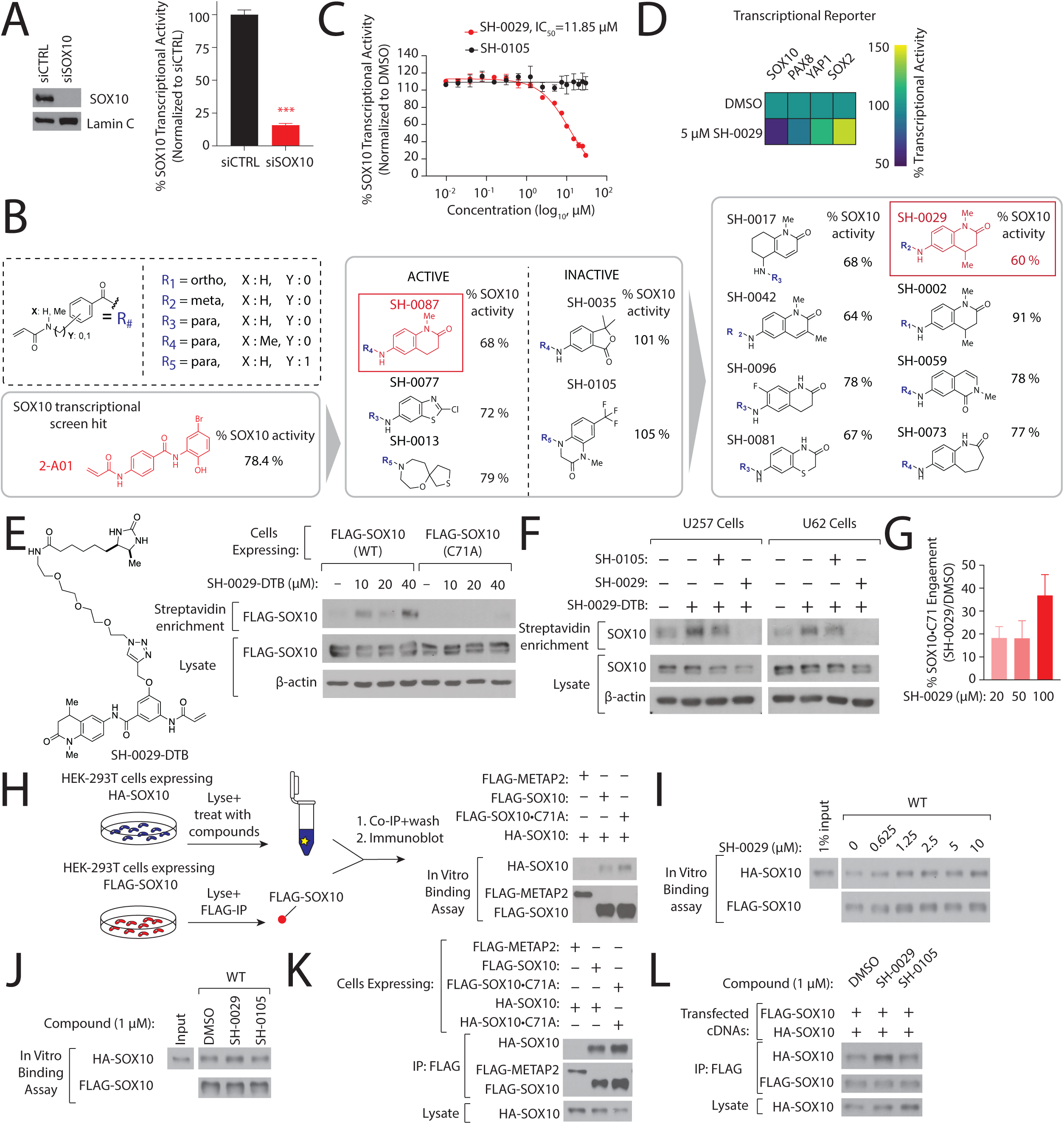
Development of a covalent probe for SOX10•C71, related to Figure 4. (A) Validation of SOX10 transcriptional reporter. Left, SOX10 protein levels following siRNA treatment in U257 cells were examined by immunoblot. Right, U257 cells harboring a luciferase-based SOX10 transcriptional reporter were transfected with the indicated siRNAs and relative transcriptional activity was determined 96 hrs later. (B) Generating an advanced SOX10 ligand. Structures and activities of hit compound 2-A01 and related derivatives, identifying SH-0029 (dark red) as the most potent analog. SOX10 transcriptional activity was determined following treatment with the indicated compounds (5 µM) for 48 hrs (see also Figure 4C). (C) SH-0029 disrupts SOX10 transcriptional activity in a dose-dependent manner. Cells were treated with SH-0029 or the control analog SH-0105 at the indicated concentrations, and SOX10 activity was measured as described in (B). (D) SH-0029 does not perturb the activity of other transcriptional reporters. Cells expressing transcriptional reporters for the indicated transcription factor reporters were treated with SH-0029, and relative transcriptional activity was determined as in (A). (E) SH-0029 ligands SOX10•C71. Left, structure of a biotinylated analog of SH-0029 (SH-0029-DTB). Right, HEK-293T cells expressing FLAG-SOX10 or FLAG-SOX10•C71A were treated with SH-0029-DTB, and SOX10 levels were assessed by immunoblot following streptavidin enrichment from corresponding cell lysates. (F) SH-0029 ligands SOX10 in melanoma cells. U257 or U62 cells were treated with vehicle, SH-0029 or SH-0105 (90 µM) for 3 hrs followed by a chase with SH-0029-DTB (30 µM) for 1 hr. SOX10 levels were determined by immunoblot as in (E). (G) SH-0029 engages SOX10•C71 in a dose-dependent manner. SKMEL5 cell lysate was treated with vehicle or the indicated concentrations of SH-0029, and SOX10•C71 engagement was determined by iso-TMT (see Methods). (H) Left, schematic depicting SOX10 *in vitro* binding assay. Right, immunoblot demonstrating HA-SOX10 or HA-SOX10•C71 binding to FLAG-tagged counterpart but no interaction with a control protein (METAP2). (I) SH-0029 increase SOX10 interaction in a dose-dependent manner. SH-0029-induced SOX10-SOX10 interactions was assessed with the SOX10 *in vitro* binding assay as in (H). (J) SH-0029 but not SH-0105 increases SOX10 binding *in vitro*. (K) Probing SOX10 binding in cells. HEK-293T cells transiently expressing the indicated FLAG- or HA-tagged proteins were lysed, and SOX10 binding was determined by immunoblot following immunoprecipitation with anti-FLAG M2 beads. (L) SH-0029 but not SH-0105 increases SOX10 binding in cells. HEK-293T cells co-expressing FLAG-SOX10 and HA-SOX10 were treated with vehicle (DMSO), SH-0029 or SH-0105 for 3 hrs, and SOX10 binding was determined as described in (K). Data are represented as mean ± SD. ***p< 0.0001. Student’s *t*-test (two-tailed, unpaired) were used to determine statistical significance.

**Figure S10:**
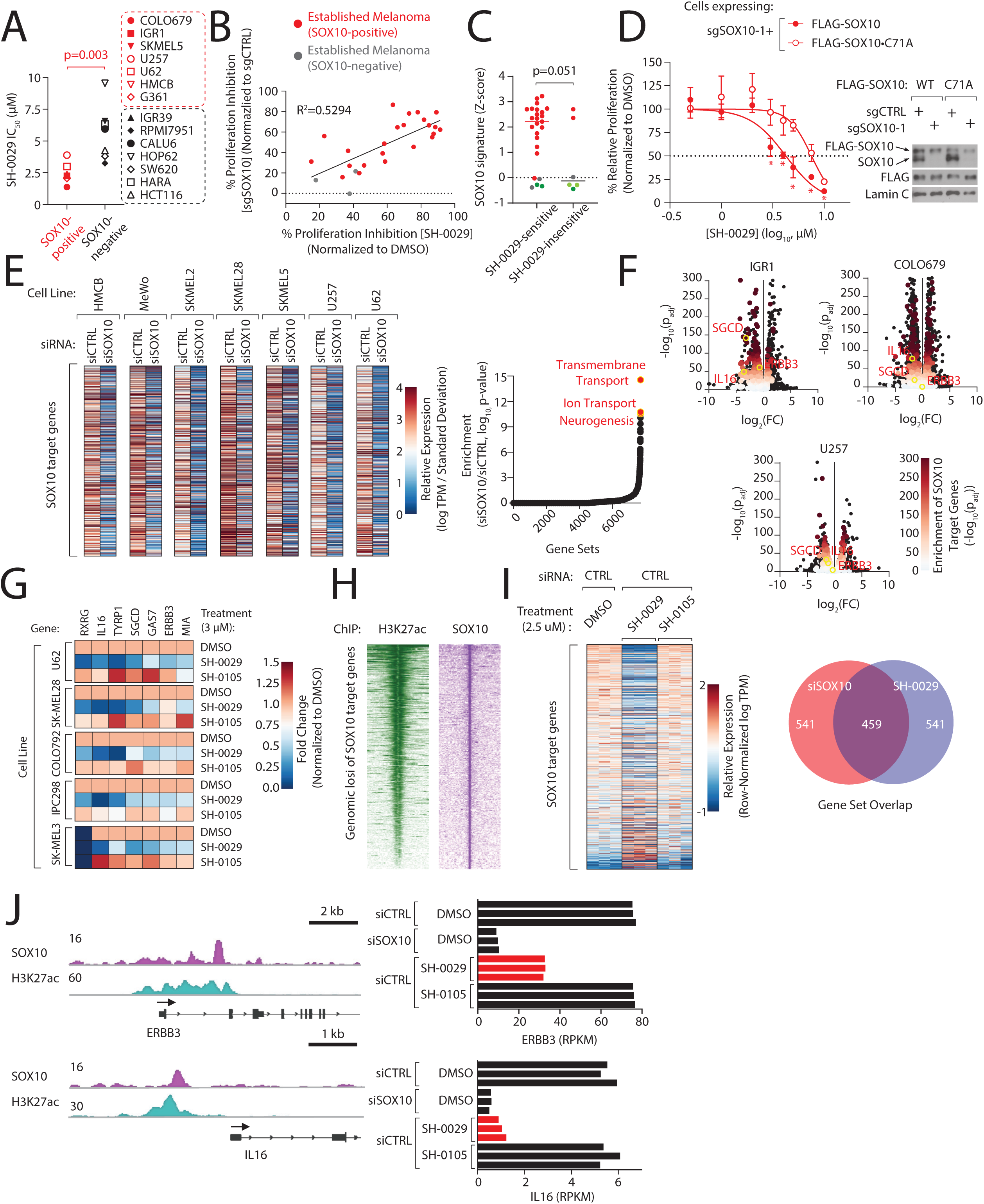
Characterization of SOX10 ligands in melanoma cells, related to Figure 4. (A) SOX10-positive cells are more sensitive to SH-0029 treatment. SH-0029 IC_50_ values were determined in SOX10-positive or -negative cell lines. Relative proliferation was determined using crystal violet staining four days post-treatment (see Methods). (B) SH-0029 sensitivity correlates with SOX10 dependency. Scatter plot comparing relative proliferation after SH-0029 treatment with relative proliferation after CRISPR-mediated SOX10 depletion. Relative proliferation was determined as in Figures 4H, **S8A**. (C) Comparison of SH-0029 sensitivity with SOX10 transcriptional signature. (D) Mutation of SOX10•C71A partially but significantly rescues SH-0029 sensitivity in U257 cells. Left, relative proliferation following SH-0029 or SH-0105 treatment in U257 cells expressing sgSOX10_1 and PAM-resistant FLAG-SOX10 or FLAG-SOX10•C71A. Right, immunoblot analysis of SOX10 levels in the indicated cell lines (see Methods). (E) Defining a SOX10 target gene signature in melanoma. Left, heatmap depicting commonly downregulated genes following treatment with siSOX10, compared to siCTRL, in a panel of melanoma cell lines. Right, GSEA enrichment plot for SOX10 target genes. Cells were transiently transfected with the indicated siRNAs, and changes in gene expression were determined by RNA-seq 72 hrs post-transfection (n=2-3 replicates/condition, see Methods). (F) SH-0029 alters melanoma transcriptional circuits. Volcano plots comparing fold change in transcript level to significance following treatment with SH-0029. SOX10 target genes are highlighted in red (see also Figure 4I). (G) SH-0029 regulates SOX1 target genes in diverse melanoma cell lines. Cells were treated with vehicle, SH-0029 or SH-0105 (3 µM, 48 hrs), and expression changes for the indicated genes were determined by qPCR. (H-I) Pharmacological and genetic regulation of the SOX10 transcriptional network. Heatmap depicting H3K27ac and SOX10 signal intensities (H) at genomic loci of genes downregulated following siSOX10 treatment in SKMEL5 cells. Heatmap of differentially expressed SOX10 target genes in SKMEL5 cells (I) following treatment with siCTRL and vehicle (DMSO), 2.5 µM SH-0029 or SH-0105 for 48 hrs (see Methods). Right, Overlap in regulated pathways following siSOX10 or SH-0029 treatment in SK-MEL5 cells (J) SH-0029 regulates SOX10-bound genes. SOX10 and H3K27ac ChIP-seq tracks for ERBB3 (top) and IL16 (bottom) and corresponding changes in transcript levels following the indicated treatments. Data are represented as mean ± SD. *p < 0.05. Student’s *t*-test (two-tailed, unpaired) was used to determine statistical significance.

## RESOURCE AVAILABILITY

### Lead contact

Further information and requests for reagents should be directed to the Lead Contact, Liron Bar-Peled.

### Materials availability

▯ All unique/stable reagents generated in this study are available from the Lead Contact with a completed Materials Transfer Agreement.

## KEY RESOURCES TABLE

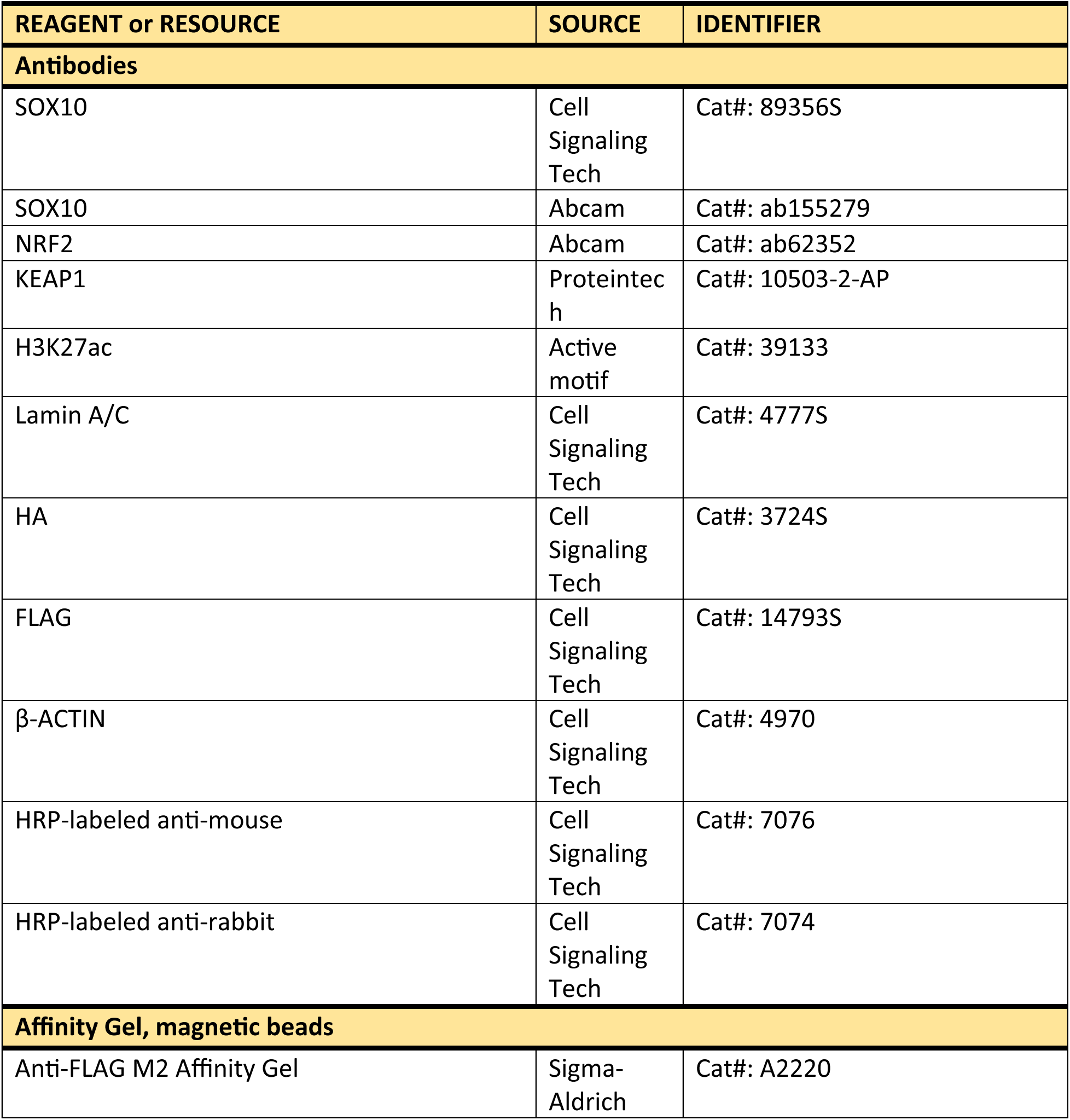
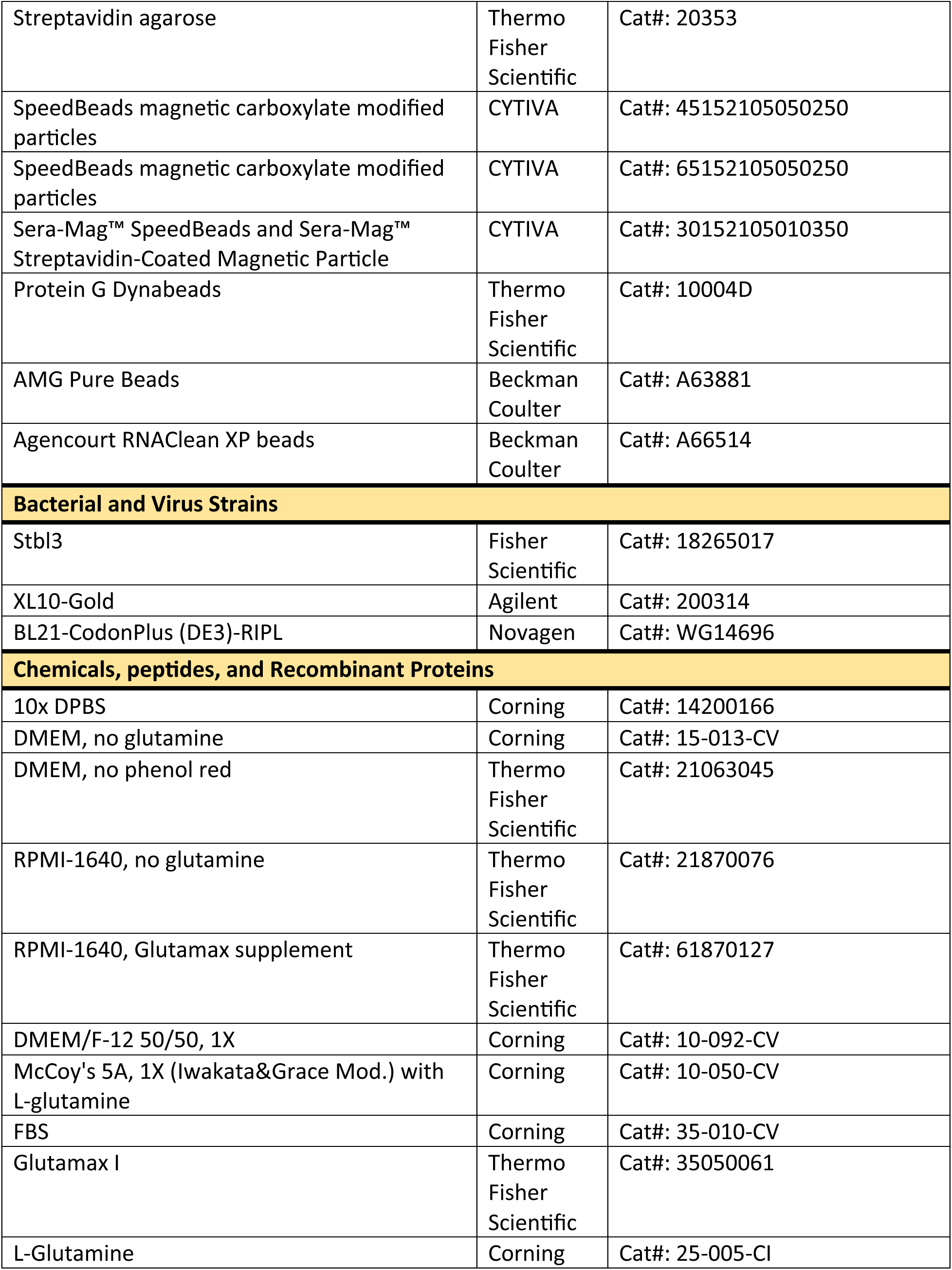
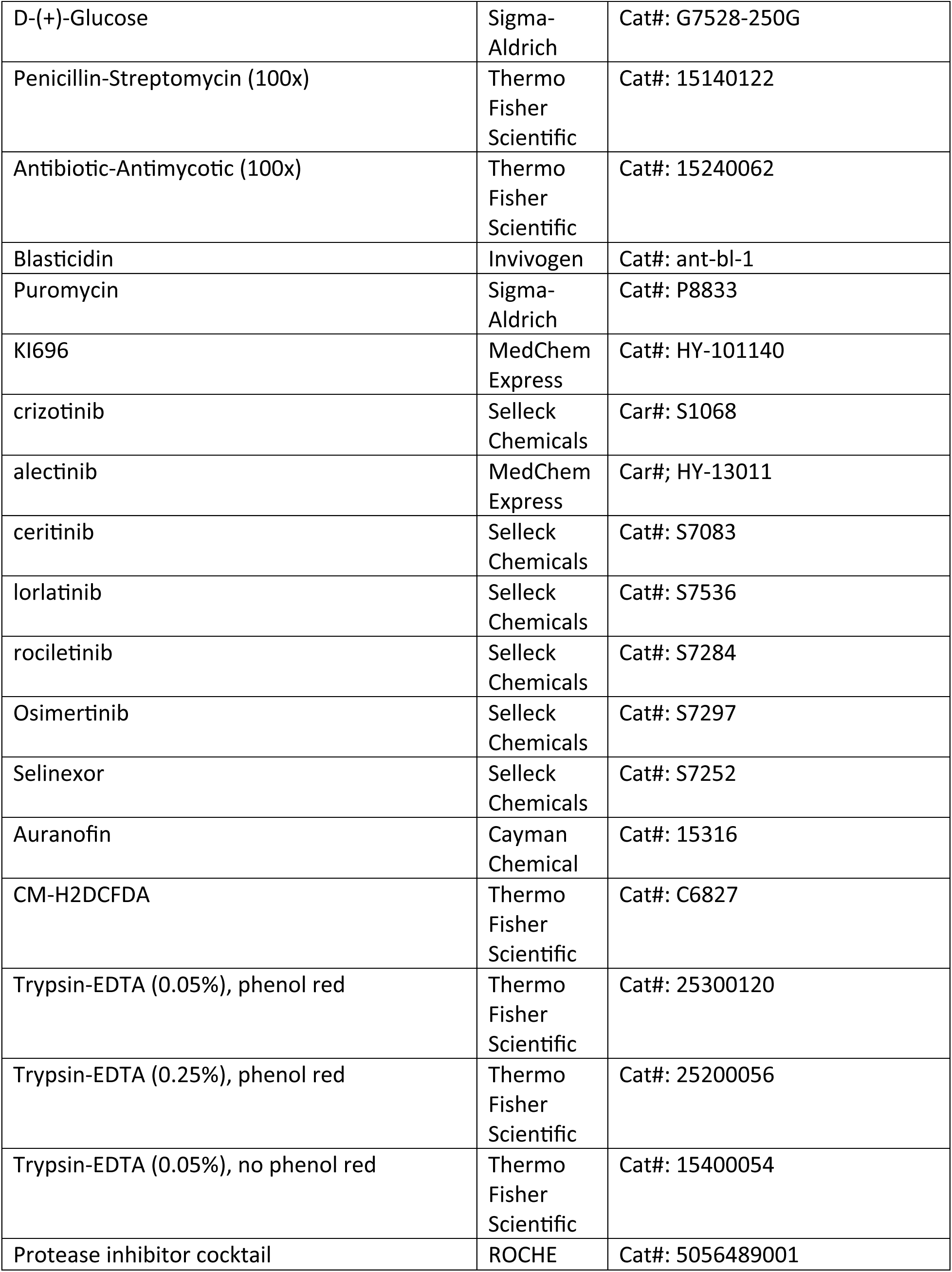
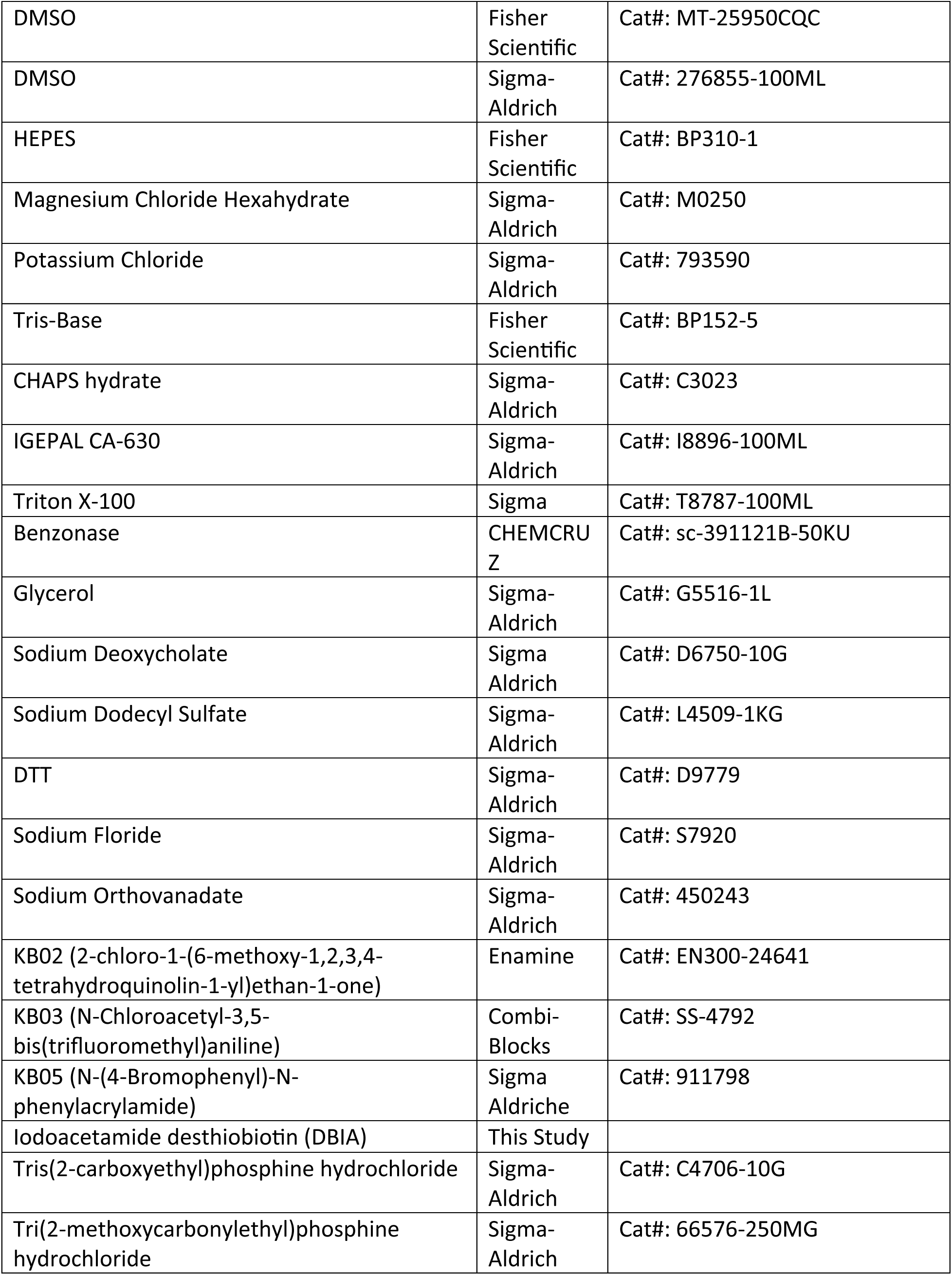
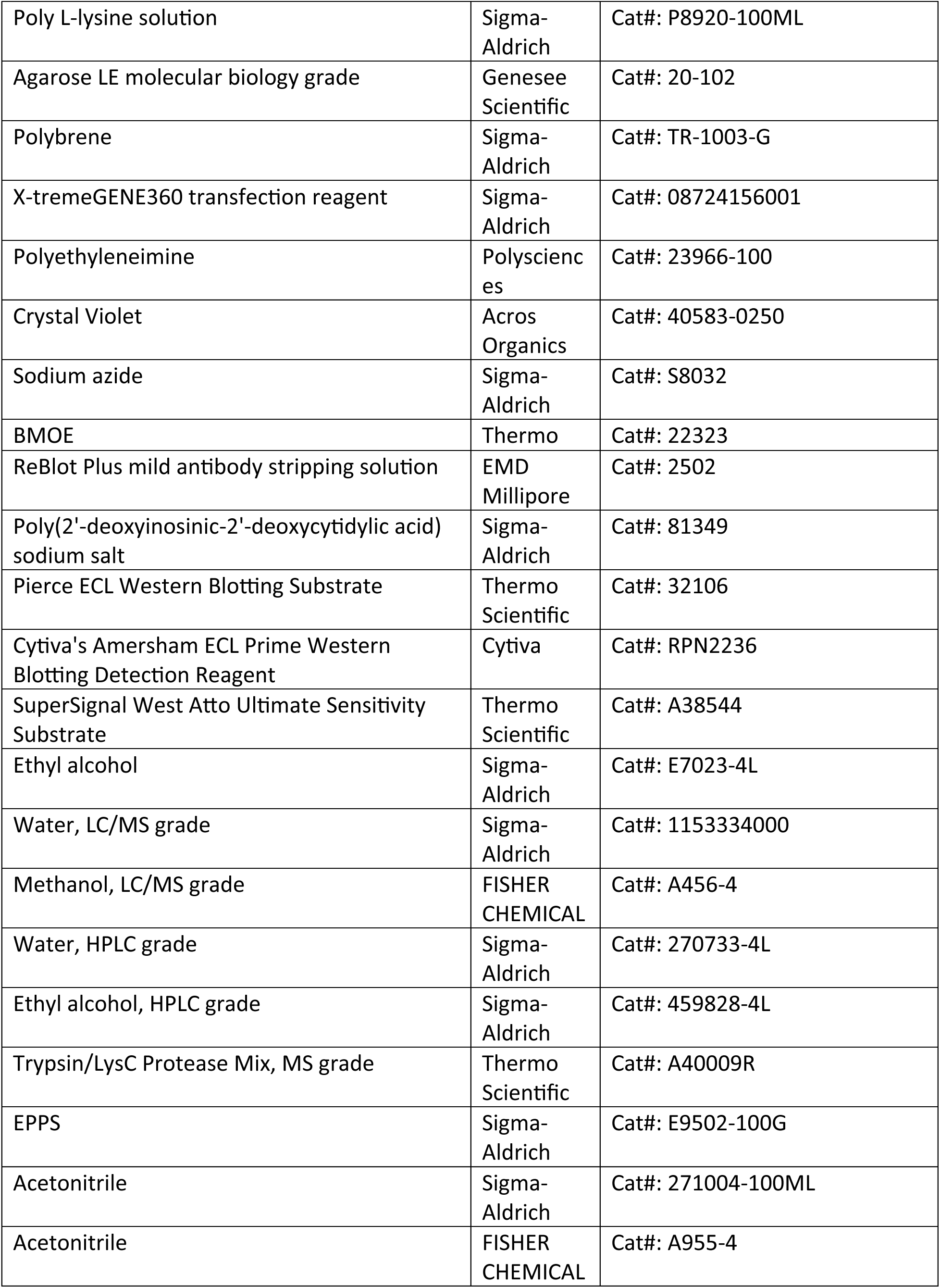
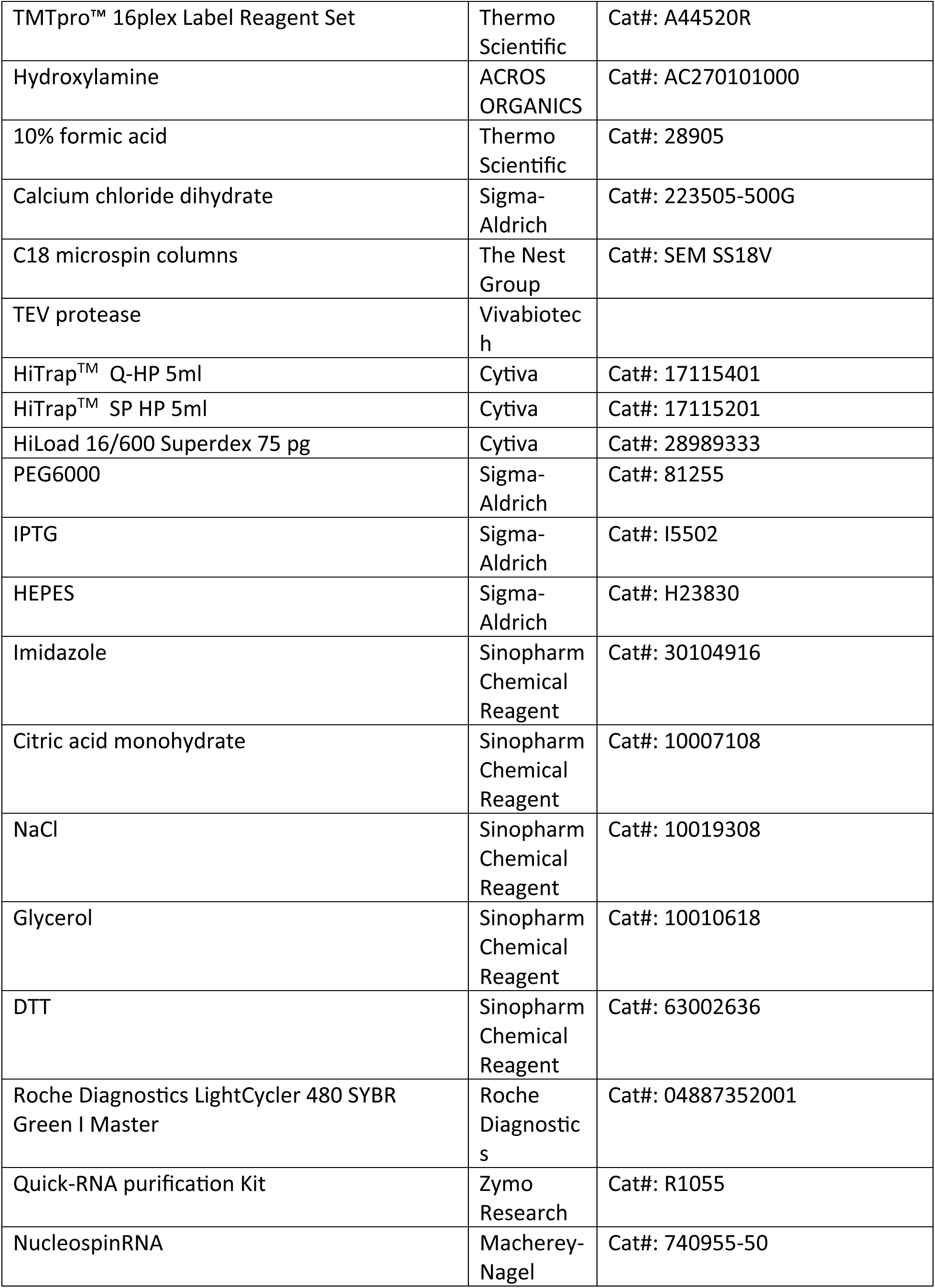
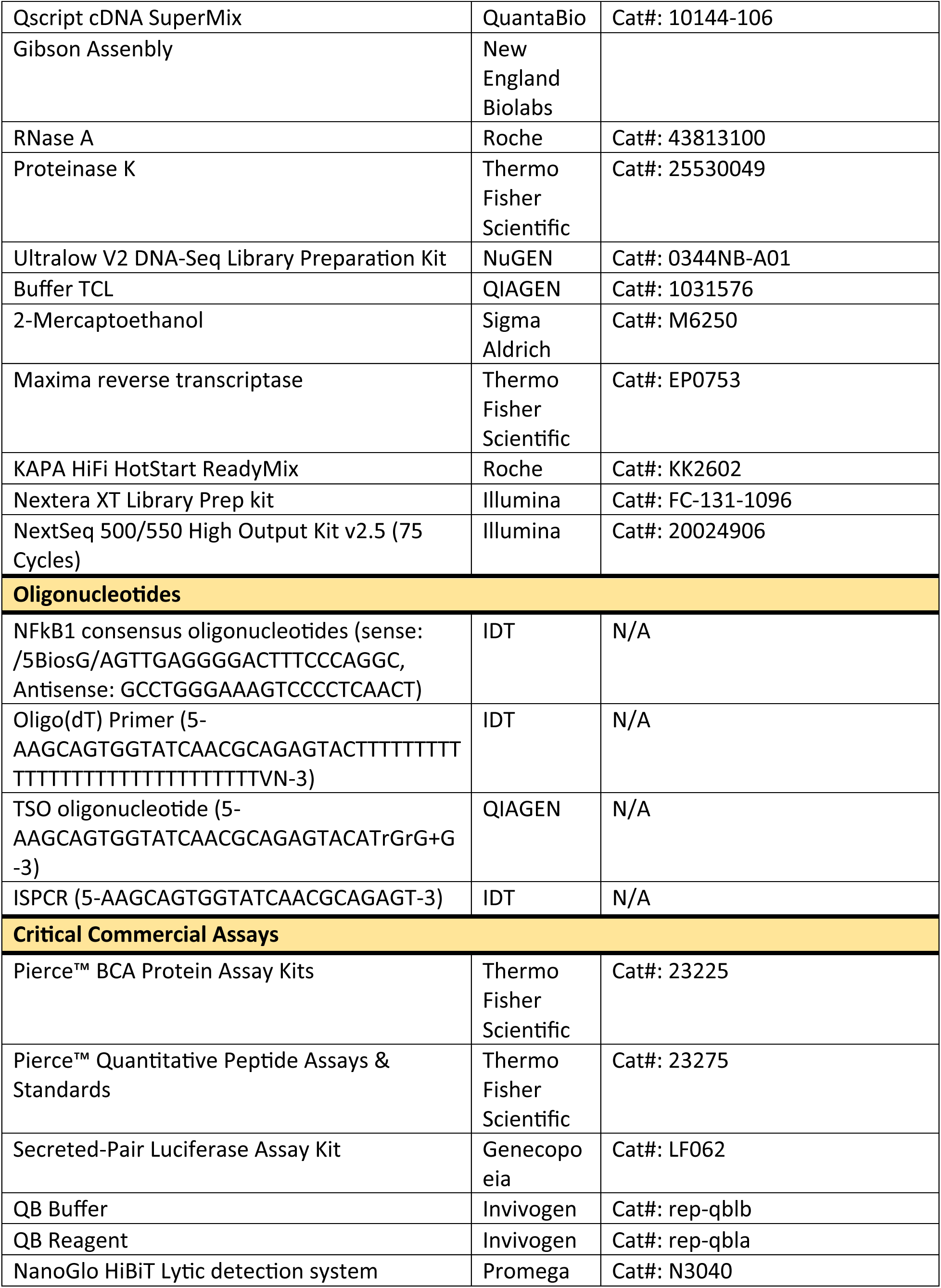
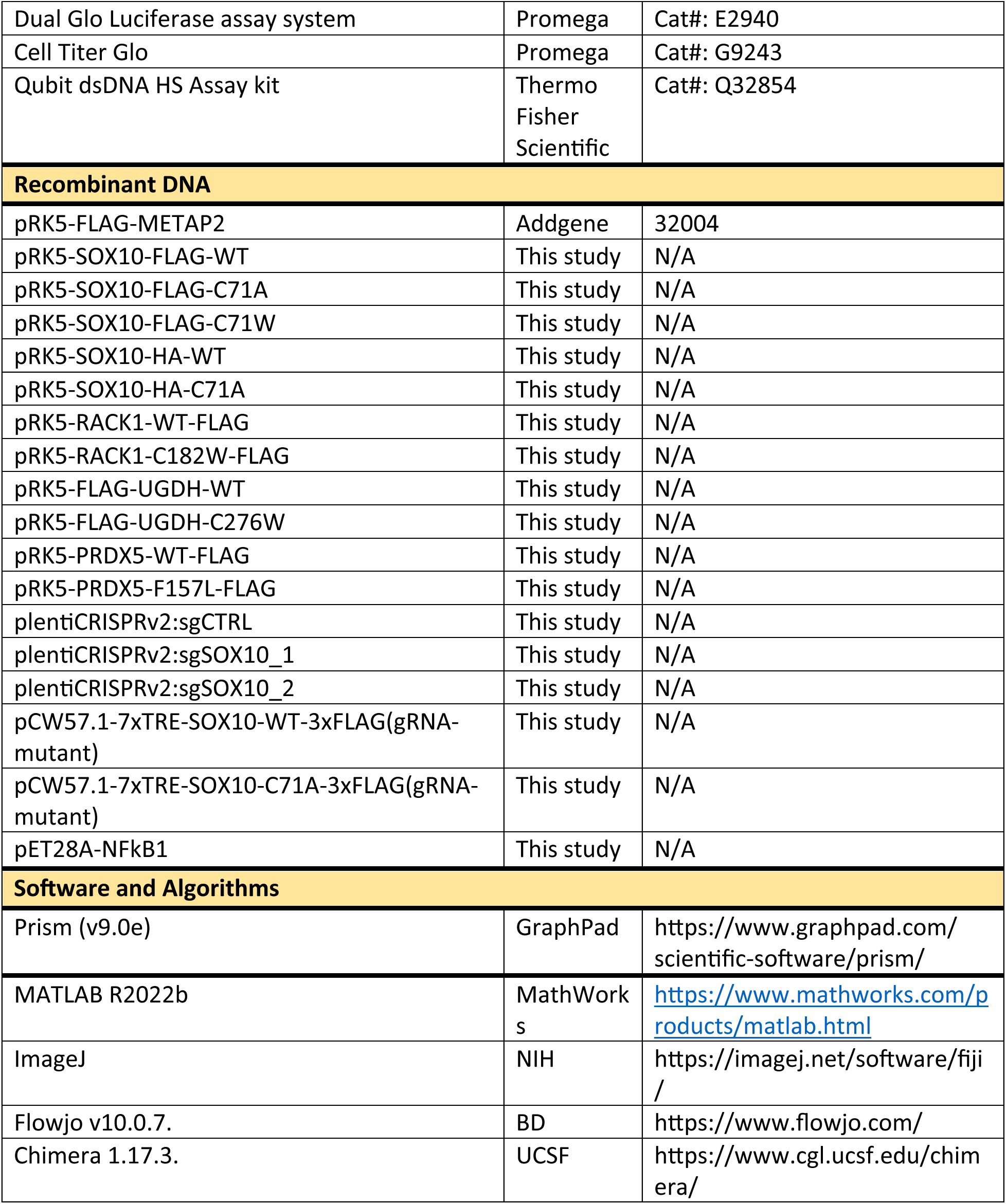

## EXPERIMENTAL MODEL AND SUBJECT DETAILS

### Cell culture

Culture methods for 416 cell lines used for DrugMap including glioma neurospheres^123^ ^124^and patient derived breast CTC cell lines^125^ are summarized in **Table S1**. A375, SKMEL2, RPMI7951, MeWo, IGR39, WM266-4, SKMEL5, HMCB, HS944T, SKMEL3, SKMEL30, HCT116, and HEK-293T were grown in DMEM (Corning) supplemented with 10% fetal bovine serum (FBS, Corning), Penicillin-Streptomycin (100 mg/ml, Thermo Fisher Scientific) and L-Glutamine (2 mM, Corning). K562, G361, U62, COLO679, IPC298, COLO792, RVH421, MELJUSO, WM793B, M234, M189, M194, M270, M258, M283, M187, M307, M321, M272, M148, M249, M171, M144, M308, M176, M175, 10164, CALU6, HARA, HOP62, OVISE, EFO27, OVTOKO, and EFO21 were grown in RPMI-1640 (Corning) supplemented with 10% fetal bovine serum (FBS, Corning), Penicillin-Streptomycin (100 mg/ml, Thermo Fisher Scieitific), and 1% GlutaMax (Thermo Fisher Scientific). SW620, SKMEL28, IGR1, A2058, C32, IGR37, LOX-IMVI, 10150, 10170, OVK18, TOV21G, and OAW28 were grown in DMEM/F12 (Corning) supplemented with 10% fetal bovine serum (FBS, Corning), Penicillin-Streptomycin (100 mg/ml, Thermo Fisher Scientific), and 1% GlutaMax (Thermo Fisher Scientifc). SKOV3 cells were grown in McCoy’s 5A (Corning) supplemented with 10% fetal bovine serum (FBS, Corning), Penicillin-Streptomycin (100 mg/ml, Thermo Fisher Scientific) and L-Glutamine (2 mM, Corning). PC9 cells were grown in RPMI (Corning) supplemented with 10% fetal bovine serum (FBS, Corning), 1% Antibiotic-Antimycotic (Thermo Fisher Scientific), and L-Glutamine (Thermo Fisher Scientific). All the cell lines were routinely tested for Mycoplasma and if not noted elsewhere were obtained from American Tissue Type Collection (ATCC).

## METHODS DETAILS

### Antibodies and Reagents

Antibodies and reagents used in this study are listed in Key Resources Table.

### Transient transfection

Transient transfections were conducted as previously described^32^. Briefly, HEK-293T cells were plated at a confluence of 3 x 10^6^ cells/10 cm dish. 24 hrs post seeding, cells were transfected with indicated plasmid in polyethyleneimine (PEI, Polysciences) with a ratio of 1:3 (DNA:PEI). Cells were harvested 48 hrs post transfection, washed once in ice-cold PBS buffer, and snap-frozen in liquid nitrogen. In experiments where cells were treated with compounds prior to harvest, tissue culture dishes were coated with poly-L-lysine (Sigma Aldrich, 1:100 diluted) for 20 min at 37°C prior to cell seeding. 48 hrs after transfection, cells were treated with SH-0029 or SH-0105 for 3h in serum-free media and then harvested as described above.

### Lentivirus generation

Lentiviral plasmids for either sgRNA expression or over-expression were cloned into the indicated lentiviral expression plasmids (see **Table S14**). Lentiviral expression plasmids were co-transfected with psPAX2 (Addgene #12260) and pMD2.G (Addgene #12259) envelope packaging plasmids into 1.8 x 10^6^ HEK293T cells using X-tremeGENE^TM^ 360 Transfection Reagent (Sigma-Aldrich). Virus-containing supernatants were collected 48 hrs after transfection.

### siRNA transfection

siRNAs were obtained from Dharmacon, and target sequences are listed in **Table S14**. siRNA oligonucleotide pools of 4 different siRNAs were transfected into melanoma cells using Lipofectamine RNAimax (Invitrogen) at a final concentration of 20 nM following the manufacturer’s instructions. Briefly, melanoma cells were trypsinized and seeded at a confluency of 3 x 10^5^ cells/well in 6-well plates with pre-plated transfection complexes incubated in Opti-MEM (Life Technologies). Culture media were replaced 24 hrs post transfection. 48 hrs post siRNA transfection, adherent cells were trypsinized and replated at a confluency of 1 x 10^4^ cells/well in 96-well plates for luciferase assays. Cells were harvested 72 hrs post-transfection to examine protein levels by immunoblotting.

### Quantitative reverse-transcriptase (qRT)-PCR

qRT-PCR was conducted as previously described with the following modificaitons^126^. Briefly, U62, SK-MEL28, COLO-792, IPC-298, SK-MEL3 cells were replated at a confluency of 1.5 x 10^5^ cells/well in 6-well plates. After 24 hrs, cells were treated with vehicle control (DMSO), 3 μM SH-0029, or SH-0105 for 48 hrs. MM1S, AMO1, RPMI8226, KMS11, and KMS27 were replated at a confluency of 5 x 10^5^ cells/well in 12-well plates and treated with vehicle (DMSO), SH-9791, or SH-1696 at the concentrations described in Figure 3 (10 μM) for 3 hrs. Total RNA was extracted with Quick-RNA extraction kits (Zymo Research) and cDNA synthesized using qScript cDNA synthesis kit (Quanta) according to manufacturer’s instructions. qRT-PCR was performed using LightCycler® 480 System (Roche) using the FastStart SYBR Green master mix (Roche). The relative expression of each gene was normalized to UBC(Ubiquitin C). Primer sequences are listed in **Table S14**.

### Cloning and mutagenesis

cDNAs were amplified using Q5 High-Fidelity 2X master mix (NEB), Phusion (NEB), or Pfu DNA polymerase (Promega) and subcloned into the pRK5 (Addgene #32004), pLJM1 (Addgene #100510) or pLX304 (Addgene #25890) by T4 ligation or Gibson cloning (NEB). Site directed mutants were constructed using primers containing the desired mutations. All constructs were verified by DNA sequencing. Lentiviral sgRNA constructs and SOX10, PAX8, YAP1, and SOX2 reporter constructs are listed in **Table S14**.

### SOX10 transcriptional reporter assays

To generate U257, SK-MEL5, HMCB, U62, and WM-2664 SOX10 reporter cell lines, cells were transduced with lentiviral particles harboring pEZX vector (Genecopoeia) with a secreted luciferase under the control of a SOX10 target gene promoter^127^. PAX8, YAP1, and SOX2 reporter cells were established in a similar manner in OVTOKO, MDA-MB-231, and H3122 cells, respectively^128–130^. To identify small molecule inhibitors of SOX10 transcriptional activity, U257 SOX10 reporter cells were replated at a confluency of 5,000 cells/well in 96-well plates. 24 hrs post seeding, cells were treated with vehicle (DMSO) or the indicated small molecule library (Enamine) for 48 hrs. To determine SOX10 transcriptional activity, culture supernatants were collected and secreted luciferase (GLUC) and secreted alkaline phosphatase (SEAP, expression control) levels were determined according to manufacturer’s instructions (Genecopoeia and Invivogen, respectively).

### Cell lysis and Immunoprecipitations

Unless noted otherwise, all lysis buffers were supplemented with protease inhibitors (Roche), 10 mM Sodium Fluoride (Sigma Aldrich), and 1 mM Sodium Orthovanadate (Sigma Aldrich). To determine melanoma protein expression, pellets were lysed with 1% Triton-X100 lysis buffer (40 mM HEPES pH 7.4, 10 mM KCl, 5 mM MgCl_2_, 1% Triton-X100) supplemented with benzonase (Santacruz Biotechnology) using a chilled bath sonicator (Q700, QSonica). Lysates were rotated for 30 min at 4°C and centrifuged at 21,000 x g for 10 min, and soluble fractions were collected. For co-transfection experiments, HEK-293T cells co-transfected with indicated plasmids were washed with PBS once and treated with SH-0029 or SH-0105 for 3 hrs in serum-free media. Plates were washed once with ice-cold PBS, snap-frozen for storage, then lysed using a chilled bath sonicator (Q700, QSonica) in CHAPS lysis buffer (40 mM HEPES pH 7.4, 10 mM KCl, 5 mM MgCl_2_, 0.3 % CHAPS). Lysates were clarified by centrifugation at 21,000 x g for 10 min. FLAG immunoprecipitations were conducted as previously described^32^. In brief, anti-FLAG M2 resin (Sigma Aldrich) was added to the HEK-293T lysates generated as described above, resin was incubated for 2 h at 4°C with end-over-end rotation. Following immunoprecipitation, beads were washed three times with CHAPS lysis buffer supplemented with 150 mM NaCl. Loading buffer was subsequently added and samples were denatured by boiling at 95°C for 5 min. Proteins were resolved by SDS-PAGE and analyzed by immunoblotting as previously described^32^ with antibodies described in **Key Resource Table**.

### Flow cytometric analysis of cellular ROS levels

Cellular ROS levels were determined using CM-H2DCFDA (Thermo Fisher Scientific) as described previously^32^. For CM-H2DCFDA staining in EFO27, SKOV3, EFO21, OVISE, OVTOKO, TOV21G, OVK18, and OAW28, cells were washed with prewarmed PBS, trypsin digested and harvested by centrifugation at 1200 x g at room temperature for 2 mins. The cell pellet was resuspended in PBS with 1 μM of CM-H2DCFDA and incubated for 30 min in a 37°C incubator with 5% CO_2_. For CM-H2DCFDA staining in K562 cells, cells were pretreated with 1 mM TMCEP (Sigma Aldrich) for 30 min or 1 μM KI696 (MedChemExpress) for 48 hrs and harvested by centrifugation at 1200 x g at room temperature for 2 mins. The cell pellet was resuspended in PBS with 1 μM of CM-H2DCFDA and 1 mM TMCEP or 1 μM KI696, then incubated for 30 min in a 37°C incubator with 5% CO_2_. Cells were subsequently washed with PBS and changes in CM-H2DCFDA fluorescence were determined via flow cytometry using Aurora (Cytek). Data was analyzed using Flowjo v10.6 for FITC intensity.

### SOX10 interaction with SH-0029-DTB

U257 or U62 melanoma cells were plated at a confluency of 4 x 10^6^ cells per 15 cm dishes. After 48 hrs, cells were treated with DMSO or SH-0029 (30, 60, or 90 μM) for 3h in serum-free RPMI, followed by SH-0029-DTB (30 μM) treatment for 1 hr. Cells were washed once with ice-cold PBS, snap-frozen for storage and then lysed in Buffer A (15 mM Tris-HCl pH 8.0, 15 mM NaCl, 60 mM KCl, 1 mM EDTA, 0.5 mM EGTA, 0.05% NP-40) for 5 min on ice. Lysates were clarified by centrifugation at 400 x g for 5 min. For streptavidin enrichment, streptavidin agarose beads(Thermo Fisher Scientific) were added to lysates and incubated for 3 hrs at 4°C with end-over-end rotation. Following enrichment, beads were washed three times with Buffer A supplemented with 150 mM NaCl. Loading buffer was added to the immunoprecipitated proteins which were subsequently denatured by boiling at 95°C for 5 min. Proteins were resolved by SDS-PAGE and analyzed by immunoblotting.

### SOX10 *in vitro* binding assays

*In vitro* binding assays were conducted as previously described^32^. In brief, HEK-293T cells expressing the indicated proteins were washed once with ice-cold PBS, snap-frozen for storage, then lysed using a chilled bath sonicator (Q700, QSonica) in CHAPS lysis buffer (40 mM HEPES pH 7.4, 10 mM KCl, 5 mM MgCl_2_, 0.3 % CHAPS). Lysates were clarified by centrifugation at 21,000 x g for 10 mins. For FLAG immunoprecipitations, anti-FLAG M2 resin (Sigma Aldrich) was added to lysate from cells expressing SOX10-FLAG, SOX10•C71A-FLAG or FLAG-METAP2 and incubated for 2 hrs at 4°C with end-over-end rotation. Lysates isolated from cells expressing SOX10-HA or SOX10•C71A-HA, were treated with DMSO or the indicated compounds for 3 hrs at 4°C and then incubated with SOX10-FLAG-bound resin for 1 hr at 4°C with end-over-end rotation. Following immunoprecipitation, beads were washed three times with CHAPS lysis buffer supplemented with 150 mM NaCl. Loading buffer was added to the immunoprecipitated proteins which were subsequently denatured by boiling at 95°C for 5 min. Proteins were resolved by SDS-PAGE and analyzed by immunoblotting.

### Thermal Shift Assays

HEK-293T pellets transiently expressing each protein were lysed with DPBS supplemented with protease inhibitors (Roche), Sodium Fluoride (Sigma Aldrich), and Sodium Orthovanadate (Sigma Aldrich), using a chilled bath sonicator (Q700, QSonica). Lysates were clarified by centrifugation at 300 × g for 3 min. Supernatants were diluted to 1.25 mg/mL using DPBS, aliquoted at 40-50 μL/well in a PCR plate, and allowed to warm to RT. Vehicle (DMSO) or the indicated compound were subsequently added, and the samples were allowed to incubate at room temperature for 1 hr in a PCR plate. Samples were heated to the indicated temperature for 3 min in a BioRad T100 Thermal Cycler (BioRad). Samples were allowed to cool at room temperature for 5 min, incubated for 3 min on ice, transferred into 1.5 mL tubes, and centrifuged at 21,000 × g for 1 hr. Soluble fractions were collected for immunoblot analysis as described above. Band intensity was assessed with Image J analysis and normalized to the value at the lowest temperature (RACK1: 36°C, UGDH: 48°C) used in each experiment. The melting curve for each protein was generated with Prism v9 (GraphPad).

### CRISPR-mediated knockdown in melanoma models

3 x 10^5^ melanoma cell lines were transduced with lentiviral particles containing non-targeting sgRNAs or sgRNAs targeting Exon 1 of SOX10 in the presence of 10 μg/mL polybrene (Millipore Sigma). 24 hrs post infection, cell culture media were replaced, and cells were cultured for an additional 24 hrs. Cells were subsequently transferred into 10 cm dishes in fresh media to culture for 3 days. Afterwards, cells were replated into 96-well plates in the presence of puromycin (0.25-1 μg/mL) for proliferation assays 6 days post-seeding (see below) or were replated in a 6 cm dish or a 10 cm dish for immunoblot analysis blot analysis following 6 days of puromycin selection.

### Cell Proliferation assays

To examine the impact of SOX10 depletion on melanoma cell lines, 1,000-7,000 melanoma cells expressing the indicated sgRNAs (see above) were seeded into a 96-well plate, and proliferation was determined 6 days post-seeding by measuring relative ATP levels as previously described^32^. In brief, 50 µL Cell Titer Glo^TM^ (Promega) was added to each sample well, and the luminescence was read on the SpectraMax M5 plate reader (Molecular Devices). To examine the impact of SH-0029 or SH-0105 on cell proliferation, 2,000-7,000 cells were seeded in 96-well plates/well in 100 µl media. The following day, cells were treated with the indicated compounds for 96 hrs. At which point culture media were removed, and cells were stained with Crystal Violet staining solution (0.5 % in 20 % methanol) for 30 min at room temperature. Viability was assessed with Image J analysis as previously described^64^. To calculate half maximal inhibitory concentrations (IC_50_), cells were treated with 0.0156-10 µM compounds for 96 hrs, cell proliferation was analyzed with crystal violet images, and IC_50_ values were calculated using log(inhibitor) vs % normalized response formula in Prism v9 (GraphPad). For proliferation In U257 stably co-expressing sgSOX10-1 and PAM-resistant FLAG-SOX10 or FLAG-SOX10•C71A, cells were replated in 96 well plates at 5,000 cells/well in 100 µl medium after six days of puromycin selection. The following day, cells were treated with 0.5-10 µM SH-0105 or SH-0029 for 96 hrs. Cell proliferation was determined by measuring relative ATP levels as described above.

### NFkB DNA HiBiT-Protein Complementation Assay

HEK-293T cells stably expressing HiBiT-tagged NFkB1 or NFkB1-C61S mutant were lysed with RIPA buffer (Pierce) supplemented with HALT Protease & Phosphatase inhibitor cocktail (Thermo Fisher Scientific) with homogenization through QIAshredder spin column (QIAGEN), and lysates were clarified by centrifugation at 15,000 rpm for 10 min. Supernatants were normalized to 0.25 μg/μl by BCA assay and incubated with 100 μM compounds for 3 hrs while shaking. Streptavidin-coated 384-well plates (Thermo Scientific) were washed with 1xTBS and preincubated with 15 nM NFkB1 consensus oligo (IDT, sense: /5BiosG/AGTTGAGGGGACTTTCCCAGGC, Antisense: GCCTGGGAAAGTCCCCTCAACT) for 1 hr to allow them to bind to plates. Plates were washed three times with 1x TBS, followed by an incubation with Protein Free Block for at least 1.5 hr. Lysates pretreated with compounds were subsequently incubated with NFkB1 oligo-coated plates for 1 hr at room temperature, and NFkB1 binding was measured by luminescence using NanoGlo HiBiT Lytic Detection System (Promega) according to manufacturer’s instructions.

### NFkB Dual Luciferase Reporter Assay

HEK-293 NFkB1 reporter (BPS Bioscience) was seeded at 8,000 cells/well in 384-well plates with phenol red-free DMEM (Thermo Fisher Scientific) culture media. After 24 hrs, cells were treated with compounds for 3 hrs at the maximum concentration of 100 μM with serial dilution. NFkB1 transcriptional activity was calculated by Firefly luminescence and Renilla luminescence using Dual-Glo Luciferase Assay System (Promega) according to manufacturer’s instruction.

### RNAseq analysis of melanomas treated with SH-0029 or SH-0105

U257, COLO679, and IGR-1 were replated at 1.5 x 10^5^ cells/well into 6-well plates. The following day cells were treated with 3 μM SH-0029 or SH-0105 for 48 hrs. For SK-MEL5 treatment, 3 x 10^5^ cells were first transfected with the indicated siRNAs in 6-well plates. 24 hrs later, cells were treated with vehicle (DMSO) or 2.5 μM SH-0029 or SH-0105 for 48 hrs. Total RNA was extracted with NucleoSpin RNA extraction kit (MACHEREY-NAGEL) according to manufacturer’s instructions, and 1500 ng/30 μL RNA was sent to Novogene for sequencing.

### RNA-seq analysis following SOX10 depletion in melanoma using modified Smart-seq2 protocol

Seven melanoma cells (HMCB, MeWO, U257, U62, SKMEL5, SKMEL28, and SKMEL2) transfected with 20 nM of each indicated siRNA (as described above) were seeded in 96-well plates (5,000 cells/well) 48 hrs post transfection. Following 24 hrs proliferation in 96-well plates, cells were lysed with 25 uL of Buffer TCL (QIAGEN) supplemented with 1% β-mercaptoethanol (Sigma Aldrich) and stored at −80°C until use. Libraries from each cell line were generated based on the Smart-seq2 protocol^131, 132^ with the following modifications. RNA from 10 µL lysates was first purified with Agencourt RNAClean XP beads (Beckman Coulter) before oligo-dT primed reverse transcription with Maxima reverse transcriptase (Thermo Scientific) and locked TSO oligonucleotide, which was followed by 10 cycle PCR amplification using KAPA HiFi HotStart ReadyMix (Roche) with subsequent Agencourt AMPure XP bead purification (Beckman Coulter). Libraries were prepared with tagmentation using the Nextera XT Library Prep kit (Illumina) with custom barcode adapters (sequences available upon request). Libraries from the seven melanoma cell lines transfected with different siRNAs with unique barcodes were combined and sequenced with paired-end, 38-base reads, using a NextSeq 500 sequencer (Illumina).

### RNAseq Data Processing

Melanoma RNA Samples were processed in Novogene. Reads were aligned using STAR v.2.4.0h. FeatureCounts was used to quantify the aligned fragments and FPKM expression values were calculated by removing duplicates and using Cufflinks v.2.2.1 and hg19 RefSeq gene definitions. DEseq2 v3.10^133, 134^ was used to perform the differential expression analysis for the datasets in this study. Genes with 2-fold changes and P value < 0.05 were defined as differentially expressed genes. GO analysis was performed using datasets downloaded from the GSEA website^42^ (https://www.gsea-msigdb.org/qsea/index.isp). After integrating the SOX10 target genes, a hypergeometric test was performed to calculate p-values for the given gene sets. FDR was calculated by p.adjust(method = “fdr”) using R.

### ChIP–seq

ChIP assays were performed using 5 million cells per sample as previously described^135^. In brief, chromatin from 1% formaldehyde-fixed cells was fragmented to a size range of 200–700 bases with a Branson 250 Sonifier. Solubilized chromatin was immunoprecipitated with 5 μg antibodies against SOX10 (Abcam) or H3K27ac (Active Motif) at 4 °C overnight. Antibody–chromatin complexes were pulled down with protein G Dynabeads (Life Technologies), washed, and then eluted. After cross-link reversal and RNAseq (Roche) and proteinase K (Thermo Fisher Scientific) treatment, immunoprecipitated DNA was extracted with AMP Pure beads (Beckman Coulter). DNA was quantified with Qubit dsDNA HS Assay kit (Invitrogen). ChIP DNA samples were used to prepare sequencing libraries with Ultralow V2 DNA-Seq Library Preparation Kit (NuGEN). A Nextseq 1000 Illumina genome analyzer was used for sequencing.

### ChIP–seq Data Processing

ChIP–seq sequencing reads were aligned to the hg19 genome using bwa v.0.7.12 with default settings. After the removal of duplicate reads using picard-tools v.1.95, aligned reads were extended to 200 bp to approximate fragment sizes. Density maps were normalized to 10 M reads. IGV was used to visualize ChIP–seq coverage at specified loci. ChIP–seq peaks were identified using MACS2 v.2.2.7.1 with a *q*-value of 10^−5^. The narrow peak setting was used for TFs while broad peaks were used for histone marks. Peaks within 2 kb of a TSS were considered promoter sites and the remaining sites were considered distal sites. Chromatin and TF signals associated with peaks were quantified using pyBigWig v0.3.18.

### Generation of recombinant NFkB1 proteins

NFκB1 (Uniprot P19838) constructs, HIS6-TEV-FLAG-NFκB1 (40-352), HIS6-TEV-FLAG-NFκB1 (40-352; C61S) and crystallization construct, and HIS6-TEV-NFκB1 (40-245), were cloned into a pET28A vector (Novagen) and overexpressed with IPTG (Sigma Aldrich) induction in BL21-DE3 codon plus cells (Novagen). Cells were lysed in 50 mM HEPES pH 8.0, 500 mM NaCl, and 5% Glycerol and captured on Nickel resin (Qiagen). The column was washed in buffer supplemented with 20 mM imidazole (Sinopharm Chemical Reagent) followed by protein elution with 250 mM imidazole. The eluate was digested with TEV protease (Vivabiotech) to remove the tag, and flowed over a second, subtractive nickel column. Protein was diluted into 20 mM HEPES pH 7.0, 1mM DTT. For ion exchange, NFkB1 used in biochemical experiments was applied to a HiTrap Q HP column (Cytiva) while the crystallization construct was applied to a HiTrap SP HP column (Cytiva). Proteins were eluted with a linear gradient of 20 mM HEPES pH 7.0, 1 M NaCl, and 1 mM DTT. Ion exchange fractions were pooled, concentrated, and applied to a Superdex 75,16/600GL column for size exclusion in 20 mM HEPES pH 7.0, 250 mM NaCl, and 1 mM DTT. Proteins were exchanged into 20 mM HEPES pH 7.0, 250 mM NaCl, and 0.1 mM DTT, concentrated, and flash frozen for storage.

### Crystallization, Structure Solution, and Refinement

Protein was diluted with its storage buffer and incubated with a 5-fold molar excess of compound for 3 hrs at room temperature. Compound-bound protein was concentrated to 10 mg/mL and mixed 1:1 with 0.1 M Citrate (Sigma Aldrich) pH 5.0, 20% w/v PEG 6000 (Sigma Aldrich) for crystallization. A 2.02 angstrom dataset was collected at SPring-8 (BL45XU) and processed with XDS^136^, POINTLESS, and AIMLESS^137^. PDB 1SVC was used for molecular replacement. Protein modeling and building were performed in Coot^82^ and refined with Refmac5^138^. The model was refined using TLS and required twin refinement due to a significant twin fraction.

### iso-TMT sample preparation

Iso-TMT samples were prepared as previously described^32, 35, 38^. In brief, adherent or suspension cells were cultured until reaching ∼80% confluency. Cells were washed once with ice-cold PBS, snap-frozen in liquid nitrogen, and stored at −80°C until use. To measure the impact of intracellular redox on cysteine ligandability, K562 cells were seeded at 2 x 10^5^ cells/ml and treated with 1 mM Tris(2-methoxycarbonylethyl)phosphine (TMCEP) (Sigma-Aldrich) for 1 hr or 1 μM KI696 (MedChem Express) for 48 hrs and subsequently harvested as described above. Frozen cell pellets were lysed in DPBS supplemented with Benzonase (Santacruz) and protease inhibitors (Roche) using a chilled bath sonicator (QSONICA) and centrifuged for 3 min at 300 x *g*. Proteins were quantified by BCA assay (Thermo Fisher Scientific) and a total of 50 μg of total protein extracts were used per compound treatment. Lysates were treated with vehicle (DMSO) or 500 μM of KB02, KB03, or KB05 (Sigma-Aldrich) for 1 hr, followed by 1 mM DBIA treatment for 1 hr. For NFkB1 hit finding, lysates were treated with 200 μM compound for 90 minutes and subsequently treated with 1 mM DBIA for 1 hr. For experiments that investigated UGDH complex formation, K562 cell lysates were treated with 20, 100, or 1000 μM UDP-xylose (UGA Complex Carbohydrate Resource Center) for 1 hr before KB05 treatment. For experiments that investigated EGFR or XPO1 liganding, PC9 or K562 cell lysates were treated with 1 μM Osimertinib (Selleck Chem) or Selinexor (Selleck Chem), respectively, for 1 hr before KB05 treatment.

Following DBIA incubation, lysates were reduced with 5 mM 5-tris(2-carboxyethyl)phosphine hydrochloride (TCEP) (Sigma-Aldrich) for 2 min at room temperature, followed by alkylation using 20 mM Iodoacetamide (Sigma-Aldrich) for 30 min in the dark at room temperature. Proteins were precipitated using SP3 magnetic beads. In brief, SP3 magnetic beads (Cytiva) were prewashed with LC-MS grade water (Sigma Aldrich), and 250 µg combined SP3 beads (1:1, hydrophobic:hydrophilic) and LC-MS grade ethanol (Sigma Aldrich)were added to each sample to reach a final concentration of 50% ethanol^139^. SP3 incubation was performed for 30 min at room temperature, and beads were subsequently washed 3 times with 80% HPLC grade-ethanol (Sigma Aldrich) and then resuspended with 175 uL of Trypsin/Lys-C (1 µg, Thermo Fisher Scientific) in 200 mM EPPS (Sigma Aldrich) pH 8.4, 5 mM CaCl_2_. Proteins were digested overnight (16 h) at 37°C and digested peptides were enriched with streptavidin magnetic beads (Cytiva) for 1 hr at room temperature. Beads were subsequently washed three times with DPBS, twice with HPLC grade-water (Sigma Aldrich). Peptides were eluted with 50 % acetonitrile (Sigma Aldrich), 0.1% formic acid (Thermo Fisher Scientific), and dried using a Speedvac (Thermo Fisher Scientific).

Cysteine-enriched peptides were reconstituted with 30 % acetonitrile, 70 % 200 mM EPPS pH 8.4 and labeled with 25 µg of TMT reagent (Thermo Fisher Scientific) per channel for 75 min at room temperature with rotation. Labeling was terminated by the addition of 5% hydroxylamine (Acros Organics) for 15 min followed by addition of 10% formic acid. Samples were pooled and dried using a Speedvac (Thermo Fisher Scientific). Peptides were desalted with stage tips using the following procedure: peptides were reconstituted with 5% acetonitrile/0.1 % formic acid and loaded onto C18 Micro Spin columns (Nest Group) pre-equilibrated with LC-MS grade methanol (Fisher Chemical) and LC/MS-grade water containing 0.1% formic acid. C18 spin columns were washed 10 times with LC/MS grade water containing 0.1 % formic acid and subsequently eluted with 80% acetonitrile, 0.1% formic acid and dried using a Speedvac (Thermo Fisher Scientific).

### Mass spectrometry Data Acquisition

All mass spectrometry data were acquired using an Orbitrap Eclipse™ Tribrid™ Mass Spectrometer in-line with an Easy NanoLC-1200 system (Thermo Fisher Scientific)^37^. Peptides were separated using 75μm capillary column packed with 50 cm of C18 resin (2 μm, 100 Å; Thermo Fisher Scientific) using 180, 205, or 210 min gradients of 10–35% acetonitrile in 0.1% FA per run, unless otherwise noted. Eluted peptides were acquired by data-dependent acquisition and quantified using the synchronous precursor selection (DDA-SPS–MS3) method for TMT quantification. Briefly, MS1 spectra were acquired at 120-K resolving power with a maximum of 50-ms ion injection in the Orbitrap with high-field asymmetric-waveform ion-mobility spectrometry (FAIMS) values at −40, −50, and −70 compensate voltage (CV). MS2 spectra were acquired by selection of the top twenty most abundant features via collisional induced dissociation in the ion trap using an automatic gain control (AGC) setting of 10 K, quadrupole isolation width of 0.7 *m*/*z,* and a maximum ion accumulation time of 50 ms. These spectra were passed in real time to the external computer for online database searching. Intelligent data acquisition using Comet real-time searching (RTS) was performed with a database including cell line mutations (DepMap)^113^ and human protein databases (release_20210506)^38, 140^. The same forward- and reversed-sequence human protein databases were used for both the RTS search and the final search (Uniprot)^141^. Next, peptides were filtered using simple, initial parameters that included the following: not a match to a reversed-sequence, containing TMTPro16 isobaric tags, maximum PPM error <50, minimum PPM error >5, minimum ΔCorr 0.10. If peptide spectra matched those above, an SPS–MS3 scan was performed using up to 20 *b*- and *y*-type fragment ions as precursors with an AGC of 250 K for a maximum of 250 ms, with a normalized HCD collision energy setting of 55 (TMTPro16)^38^.

### Mass spectrometry Data Analysis

The mass spectrometry data were searched by MSFragger^142^ (v3.4), X! Tandem^143^ (v2013.06.15.1), Comet^144^ (v2021.01.rev0), MSGF+^145^ (v20220107) or Proteome Discover (2.5, Thermo Fisher Scientific) using the following parameters: 6 amino acids for minimum peptide length, 20 ppm precursor tolerance and 700 ppm fragment ion tolerance, allow tryptic peptides only, up to two missed cleavages, oxidation of methionine (+15.9949 Da) and DBIA on cysteine residues (+239.1634) as variable modifications while TMTPro16 (+304.2071 Da) on lysine and peptide N-termini, and cysteine carbamidomethylation (+57.0214 Da) were static modifications. For each mass spectrometry run, the searches were performed against a custom cell-line specific FASTA database that included the canonical UniProt human (v2021) protein sequences, common contaminants, cell-line-specific mutated peptide sequences, and reversed versions of UniProt sequences. The reversed sequences were added as decoys for target-decoy analysis. The initial search result from each search engine was then analyzed by PeptideProphet^146^. Then the PeptideProphet results from the multiple search engines were merged by iProphet^147^ analysis. Peptide ion FDRs were estimated by target-decoy approach at cell-line level based on iProphet probability. For the collective DrugMap dataset, peptides were filtered to obtain a 1% global FDR. PeptideProphet and iProphet analyses were performed via Trans-Proteomic Pipeline^148^ (TPP) (v6.0.0). TMT reporter-ion quantification from the MS3 scans of all identified PSMs was extracted by an in-house program. The raw reporter ion intensities were adjusted for impurity correction according to the manufacturer’s specifications. For quantification of each MS3 spectrum, a total sum signal-to-noise of all reporter ions of 100 (TMT16-plex) was used.

### Intact Mass Spectrometry data acquisition and processing

Recombinant NFkB1 proteins were diluted to 2.8 µM in 50 mM HEPES, 100 mM NaCl, and 0.01% Pluronic F127 and added to compound-containing wells for treatment at 10 µM. Reactions were incubated at room temperature for 3 hrs on a plate shaker and then quenched with a 1:3 dilution of 6M guanidine-HCl, 0.6% formic acid.

Following compound treatment, proteins were separated by liquid-chromatography (LC) using a Vanquish Horizon UHPLC system (Thermo Fisher Scientific) configured with a PLRP-S (Agilent) reverse-phase column. Separated proteins were introduced and measured on an Orbitrap Exploris 480 mass spectrometer (Thermo Fisher Scientific). ESI source settings for protein ionization were as follows: source voltage of 3.4 kV, heated capillary temperature of 320°C, and RF-Lens set to 50 percent. Data acquisition was performed over a mass-to-charge (m/z) range of 600-2300, at a 15,000 resolving power (at 200 m/z), an AGC target value of 1 x 10^6^, 3 mscans/spectrum, and maximum injection time of 50 ms. Intact protein MS data was deconvoluted using the open source UniDec software^149^. Relative covalent adduct formation was calculated using the intensity ratio of the adducted protein (parent + adduct) to the unmodified recombinant protein (parent).

### Calculation of Cysteine Engagement

To study cysteine ligandability, we calculated “engagement” (ε) by cysteine-reactive compounds (e.g. KB05, KB03, KB02) (Equation 1). Cysteine engagement is measured in units of percent.

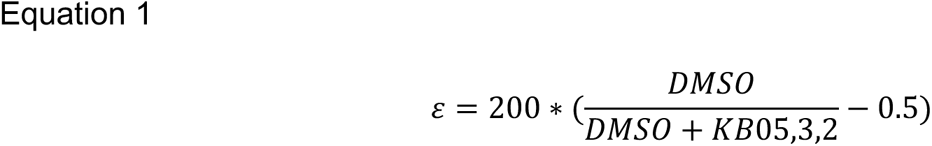

Engagement is formally equivalent to a probability of liganding by a cysteine-reactive compound, whereas the commonly calculated ratio (Equation 2) is like a mathematical odds.

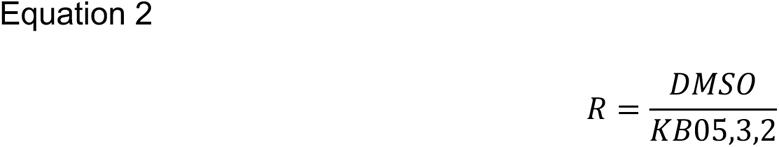

These similar constructions have different emphases. For example, ratios exhibit more prominent fluctuations at positive extremes, whereas engagement equalizes the width of the number line that is allotted to different levels of cysteine ligandability. To allow direct comparison, we provide a conversion from ratios to engagement (Equation 3).

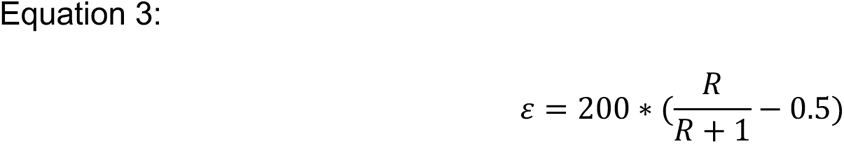

To compare these two metrics, consider a cysteine whose ratio is initially 10 and doubles to 20 in a second measurement. While this equals a 100% absolute increase in reactivity as understood with ratios, this difference only spans roughly 10 percentage points in the engagement space. Thus, engagement gives equal attention to cysteines which fluctuate from “reactive” to “highly reactive” and those which fluctuate from “not reactive” to “reactive.” To complete our data processing workflow, we partitioned the data by scout (KB05, KB03, KB02) and quantile-normalized the engagement values for all cell lines profiled, effectively creating three datasets. Per scout, this operation equalized both the centers and the widths of the distributions of cysteine engagement values across all cell lines (**Figure S1G**).

### CSEA

In order to systematically enumerate features that are unusually prevalent among a set of cysteines, such as biological pathways or protein structural elements, we developed Cysteine Set Enrichment Analysis (CSEA). CSEA requires two user-defined lists of cysteines: one should contain cysteines of interest, and the other background set should contain cysteines not expected to share meaningful sources of variation with cysteines in the set of interest. CSEA uses the background set as a reference against which to detect unexpectedly high enrichment of a feature among the cysteines of interest. We compiled >6000 sets of cysteines, conceptually binned into four kinds of set, based on: molecular functions, biological pathways, experimental literature, and protein structures (**Table S16**). This repository constitutes a library against which users can check for features enriched in their cysteines of interest. CSEA can be run against the entire library, or any subset of it—even a single cysteine set.

CSEA uses the common permutation test^42^ to establish the significant enrichment of a particular feature in a user-defined set of interest. Provided an appropriately large background set of cysteines, CSEA will tabulate the overlap between random draws of cysteines from the background set and a set of cysteines from the library, e.g. the set of all cysteines which belong to myosin motor domains. This process is formally equivalent to evaluating the intersection in a two-way Venn diagram. Each overlap calculation represents one permutation. Users can choose how many permutations CSEA performs, but the results typically stabilize after ∼200 permutations. Collectively, these permutations form an empirical null distribution against which the “true” overlap between the cysteines of interest and the feature in question is compared. This comparison is made mathematically formal by fitting a Gaussian to the null overlap distribution (Equation 4),

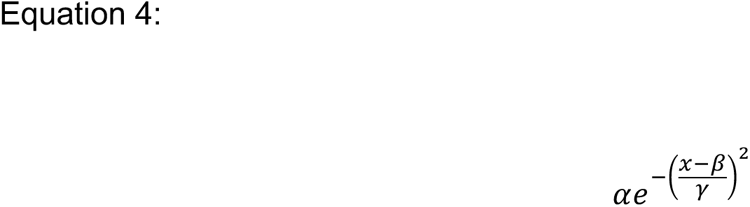

where α, β, and γ are fitting constants, and *x* is the probability density of the null distribution. After fitting, a one-sided tail integration is computed from the “true overlap” to infinity. This tail area equals the p value for the feature of interest. In a case where the null set is completely devoid of any cysteines having the feature studied, CSEA interpolates a Gaussian with minimal density near 0 in order to facilitate p-value calculation. P-values generated in this case are denoted in the output; however, this use illustrates one setting where the user-defined background may be suboptimal and warrant further consideration by the user. This procedure is repeated for each feature or library set the user wishes to evaluate. The Benjamini-Hochberg method is then applied to control the false discovery rate in a set of p-values generated over a single run. Users are free to substitute more stringent, family-wise error rate correction procedures, such as the Bonferroni correction.

### Circos Plot

The Circos plot summarizes variations in cysteine liganding across the panel of cell lines profiled in this study. The plot was calculated with respect to scout KB03. As the plot is read from the center to the periphery, information is summarized with increasing density. The innermost ring displays differences in cysteine ligandability at single sites among individual cell lines, whereas the outermost ring represents the ligandability of all cysteines detected in a particular signaling pathway, summarized across all cell lines of a particular tissue. Lineage-specific information in the Circos plot is organized angularly.

The innermost ring of the Circos plot highlights a “batch-aware” analysis, wherein the quantile-normalized engagement values of different cell lines are grouped by batch and the batch median is subtracted, yielding a matrix of differences instead of absolute engagement values. This step is common to routine batch correction methods^150^ and serves to partially mitigate the complex batch effects innate to biological data, facilitating analyses of cysteine ligandability across groups of cell lines. For example, the innermost ring of the Circos plot highlights local variation in cysteine ligandability that might be specific to a single cancer cell line. This “batch-aware” analysis underpins our analysis of heterogeneous liganding (see “Cysteine Heterogeneity” below).

In the innermost ring, all cell lines belonging to the labelled lineages are plotted, causing some slices to appear more granular than others. The second ring of the Circos plot highlights the ligandability of cysteine-containing protein domains, stratified by lineage. We represented the 49 most commonlydetected protein domains across our panel of cell lines. For each of these domains, and specifically for each lineage, we show how often the cysteines in a particular domain were liganded. The third ring of the Circos plot displays the ligandability of 12 broad protein classes and their subclasses as defined by the PANTHER database^151^. Similar to the second ring, the amount of color represents the fraction of cysteines in proteins of a particular class that were liganded in that lineage. The outermost ring of the Circos plot shows the maximum ligandability of all detected cysteines belonging to each of 14 PANTHER signaling pathways^151^, subsetted by lineage.

### Structural Analysis of Cysteine Ligandability

In order to analyze the structural underpinnings of cysteine ligandability, we built a database of protein biophysical parameters (**Table S13**), each characterizing a facet of the geometric or chemical/electrostatic locale around a cysteine. We first downloaded all PDBs of human proteins that been deposited at wwPDB.org as of November 2022 and divided them into their biological assemblies. We then implemented custom scripts to filter out PDBs lacking cysteines, and mapped the remaining PDB chains to the Uniprot database. We chose a PDB to represent each cysteine, prioritizing by best coverage of the protein’s amino acid sequence, and using highest resolution as a tiebreaker. We then analyzed our entire directory of mapped PDBs to collect structural parameters of all cysteines, using custom Python scripts to calculate the Euclidean distances between neighboring amino acids, and the Bio.PDB module of the Biopython package, the DSSP package^48, 51^, the p2rank^60^ and DeepPocket^58^ pocket prediction algorithms, and the Voss Volume Voxelator to calculate the other parameters discussed in this study.

### Neural Network Prediction of Cysteine Ligandability

The neural network in this study features two components: a convolutional neural network (CNN) and a feed-forward network, i.e. a multi-layered perceptron (MLP). The MLP was fed tabularized biophysical calculations on cysteines (the features computed above) constituting a complete structural parametrization, plus corresponding cysteine liganding data from the mass spectrometry analysis. A set of 30-row x 20-column binary distance matrices, with each column representing an amino acid and each row a half-Angstrom-thick shell centered on a cysteine of interest, were treated as images and fed into the CNN. We designed the CNN to convolve along the distances in 1D in order to abstract the spatial arrangement of amino acids surrounding a particular cysteine. The output from these networks was then concatenated and sent through further feed-forward layers (with dropout) before returning a classification (ligandability prediction) for a single cysteine. The network was trained with the python TensorFlow package.

### Cysteine Heterogeneity

In order to identify cysteines with unusually high variance in ligandability, we integrated both “global” and “local” analyses of heterogeneity. With respect to scout KB03 alone, we first calculated the coefficient of variation (CV) of cysteine ligandability across all cell lines, effectively summarizing macroscopic or global fluctuations of cysteine ligandability across our entire panel of 416 cell lines into a single vector. We then calculated “local” variance by repeating the previous calculation but instead grouping cell lines by batch, producing a CV matrix where columns and rows represented batches and cysteines, respectively. In order to harmonize these global and local views of heterogeneity, we first collapsed the local CV matrix into a column vector by counting the number of times a particular cysteine had CV>10% in quantile-normalized engagement within a batch. The choice to strictly threshold CV here allowed us to prioritize strong intra-batch ligandability differences. At this point, the data was reduced to two vectors, which were then individually rank-ordered. Cysteines were finally nominated as heterogeneous if they scored within the top 5% of each vector. Such a cysteine is thus both heterogeneous at the global level (“on average”) and also frequently heterogeneous at the local level (within batches). Lastly, in order to isolate cysteines whose reactivities are potentially controlled by processes that vary across diverse cellular contexts and are not merely singleton events, such as private amino acid mutations, we required that any candidate heterogeneous cysteine be detected in at least 50% of cell lines profiled in this study.

### Network analysis of heterogeneous cysteines

The MATLAB corr() function was used to calculate pairwise correlation coefficients between all pairs of cysteines. These correlation coefficients were taken to characterize edges between two cysteines. The MATLAB clustergram() function was used to cluster the cysteine-cysteine correlations, forming networks that could be analyzed at the global level.

### Integration of cysteine ligandability data with cell-intrinsic molecular feature data

All gene expression, mutation, and CRISPR-essentiality data were downloaded from the Cancer Cell Line Encyclopedia’s (CCLE) data portal^152^. Cell lines that underwent RNA-Seq, WGS, and genome-wide CRISPR screening, as well as cysteine ligandability profiling were included for analysis. We identified RNA expression clusters by first calculating each gene’s standard deviation of log TPM values across all cell lines and then using the 2000 highest-scoring genes to partition the cell lines under study into three transcriptionally defined clusters. Further sub-clustering was then performed at the level of each molecular feature. For example, at the genetic dependency level, CRISPR scores were first row-normalized by z-score, and then the 2,000 genes with the highest standard deviation in each major cluster, totaling 6,000 genes, were combined and row-clustered. Columns were sorted by embryonic tissue of origin. The same operation was repeated at the RNA level, except that log TPM values were first row-clustered, and we then visualized row-normalized z-scores of the transcript levels. For each cell line, the max cysteine ligandability was taken for each protein, allowing us to match the rows in the cysteine liganding heatmap to those shown at the genetic dependency level. At the genome level, >1000 commonly mutated genes organized in KEGG^153^– and PANTHER^151^–based oncogenic and signaling pathways, respectively, are shown. Proteins containing at least one missense mutation were considered mutated. A binary matrix indicating protein mutational status was clustered, and the row sorting was used to visualize somatic mutations across the panel of cancer cell lines.

### Identification of mutations that associate with changes in cysteine ligandability

In order to find genetic variants that influence cysteine ligandability, we enumerated *cis*-associations between protein mutation status and local differences in cysteine engagement. We first intersected the cell lines profiled in this study with those profiled by WGS in the CCLE. We then sorted all proteins by the genomic coordinates of their missense mutations and plotted the “batch-aware” cysteine engagement values of any cysteine quantified within that mutant protein. Our analysis assumes that, absent a protein mutation, a cysteine expressed in one cell line should display roughly the same ligandability as it does in another cell line. On this premise, the batch-aware cysteine engagement value represents the effect size of a corresponding, putatively causal missense variant within the same protein. Thus, it allows identification of singleton events (i.e. mutations observed in only one cell line) that might influence cysteine ligandability. What we loosely term an “association” is predicated on the intuitive notion that differences in cysteine ligandability among a handful of cell lines profiled in the same batch—a local form of variation— can be associated with a corresponding amino acid mutation, either proximal or distal to a cysteine of interest.

### Logistic Regression Analysis

In order to transition from singleton changes in cysteine ligandability to bona-fide associations between amino acid mutation and alterations in cysteine ligandability, we performed a cysteinome-wide logistic regression analysis. Specifically, we asked whether local differences in cysteine engagement, rather than cysteine engagement itself, can be used to infer the mutational status of a protein. Toward this end, we first restricted our screen to cysteines that were detected in at least 25 cell lines and missense-mutated in at least 10 of them. We then partitioned the batch-aware, quantile-normalized cysteine engagement values of the relevant cell lines into a “wild-type” category and a “mutant” category. For our initial analysis, we did not sub-stratify by the mutant amino acid. We then performed a logistic regression via the MATLAB mnrfit() function according to Equation 5:

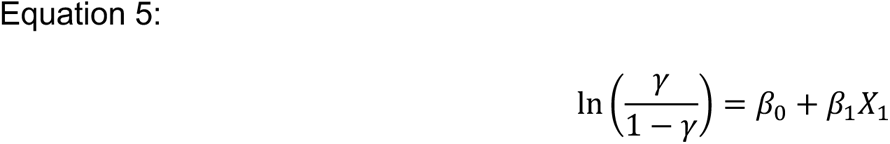

where *γ* represents the probability that a measurement *X*_1_ i.e. a single batch-aware, quantile-normalized engagement value, belongs to the “wild-type” category not the “mutant” category, and the *β_i_* are coefficients learned by the model. Recalling that the batch-aware analysis returns a matrix of local differences, the y-intercept of this model then intuitively represents the log-odds that, given the observed distribution of batch-aware engagement data, a difference of zero corresponds to a wild-type protein. Thus, we z-scored these intercepts and multiplied them by −1 to yield a log-odds of membership in the “mutant” category. In order to ascertain the direction of change between mutant and wild-type, we subtracted the median signed difference of cell lines harboring a missense mutation in the gene encoding that cysteine from that of the corresponding wild-type cell lines.

### Identification of Lineage-restricted Dependencies

In order to delineate targets for chemical probe development, we identified genes that exhibit lineage-restricted genetic dependency and also contain ligandable cysteines. We first intersected the cell lines profiled in this study with those that underwent corresponding CRISPR-based knockout and RNA-Seq profiling in DepMap. We then row-normalized the data reported for both ‘omic features (i.e. CRISPR scores and RNA-Seq log TPM) and summarized these two tables into their constituent lineage subtypes as reported in CCLE. Specifically, we collapsed columns by taking medians across all available cell lines belonging to a single lineage subtype (e.g. melanoma). At the level of lineage subtypes and not individual cell lines, we then estimated the relative importance of a particular gene within a lineage subtype by subtracting the row-normalized values of all other lineage subtypes from the one in question, forming a distribution of differences. For example, in this study we profiled cysteine ligandability in >20 tissue types spanning ∼70 lineage subtypes, meaning that ∼70 subtractions were performed for each gene, where one lineage subtype at a time was held constant. We summarized each distribution by taking its median. In principle, any gene showing persistent, positive differences, as encapsulated in the median difference, across tens of lineage subtypes, should be uniquely important for cell lines of that lineage subtype, but not other lineages. We finally calculated the inner product of the different matrices for each ‘omic feature, obtaining a final “score” for all genes jointly assayed in both datasets, across all lineages. Higher scores indicate mutual elevations in both transcriptional exclusivity and CRISPR-based essentiality for a particular gene within a given lineage subtype. The angular coordinate of the plot is used to summarize scores for each subtype within a particular lineage, and the radial coordinate indicates the magnitude of the score. Negative scores were filtered out. Given that transcription factors constituted the class of proteins most enriched at the periphery of the plot (as evidenced by simple counting), we subsetted to transcription factors. The maximum ligandability, per lineage, of cysteines belonging to multiple transcription factors detected in this study are shown.

### NRF2 and SOX10 Signature Calculation

To identify cell lines characterized by high levels of NRF2 activity with corresponding KEAP1 mutations, we intersected the cell lines profiled in this study with those that underwent WGS and RNA-Seq profiling in the CCLE. For this subset of jointly profiled cell lines, we first calculated row-normalized z-scores on the log TPM data. This step accentuated the relative differences in expression level across cell lines, preventing housekeeping genes with high basal transcription from dominating our transcriptional signature. We then calculated the median of these row-normalized transcript levels across canonical NRF2 target genes^64^, forming a distribution of NRF2 transcriptional activity. Cell lines with missense mutations in KEAP1 and high NRF2 transcriptional activity were operationally defined as possessing functionally-inactivated KEAP1 and were included for cysteine reactivity analyses. To estimate SOX10 activity in cell lines, we first calculated row-normalized z-scores on the log TPM data in the CCLE. Specifically, for each transcript, z-scores were calculated across all available cell lines, irrespective of tissue of origin. We then calculated the median of these row-normalized transcript levels across canonical SOX10 target genes^87, 154, 155^. This step defined a single number, i.e. a SOX10 signature level, for each cell line.

### Visualization of Protein Structures

Scenes containing representations of protein structures were generated in Chimera 1.17.3^156^.

### Quantification and statistical analysis

Statistical analyses were performed with Excel (Microsoft), Prism (GraphPad v9.5.1), and MATLAB (R2022b). Error bars represent mean ± s.d. Statistical significance was assessed using unpaired, two-tailed Student’s t-tests. P values are indicated in figure legends and source data. P < 0.05 is indicated with single asterisks, P < 0.001 with double asterisks, and P < 0.0001 with triple asterisks.

### Compound synthesis

Reagents and solvents were obtained from commercial suppliers and used without further purification, unless otherwise stated. Reactions were carried out under a positive atmosphere of nitrogen and monitored by thin layer chromatography (TLC) using shortwave UV light or by SHIMADZU LC/MS. Normal phase flash column chromatography was performed using silica gel 60 N (spherical, 75-150 μm). ^1^H NMR spectra were recorded on Bruker Avance 400 MHz spectrometer and were calibrated using residual non-deuterated solvent as the internal references (CDCl_3_: 7.26 ppm;; MeOH-D_4_: 3.31 ppm, acetone-D_6:_ 2.05 ppm; DMSO-D_6_: 2.50 ppm). The following abbreviations were used to explain NMR peak multiplicities: s = singlet, d = doublet, t = triplet, q = quartet, p = pentet, m = multiplet, br = broad.

### Analytic procedures

#### LCMS

##### Instrument specifications

Agilent 1100 Series LC/MSD system with DAD\ELSD Alltech 2000ES and Agilent LC\MSD VL (G1956B), SL (G1956B) mass-spectrometer.

Agilent 1200 Series LC/MSD system with DAD\ELSD Alltech 3300 and Agilent LC\MSD G6130A, G6120B mass- spectrometer.

Agilent Technologies 1260 Infinity LC/MSD system with DAD\ELSD Alltech 3300 and Agilent LC\MSD G6120B mass- spectrometer.

Agilent Technologies 1260 Infinity II LC/MSD system with DAD\ELSD G7102A 1290 Infinity II and Agilent LC\MSD G6120B mass-spectrometer.

##### General Parameters

Detection: DAD – DAD1A 215 nm, DAD1B 254 nm MSD – single quadrupole, AP-ESI (positive/negative mode switching)

Temperature: 25 °C

##### LCMS procedure

Column: InfinityLab Poroshell 120 SB-C18 4.6×30mm 2.7 Micron with Guard: UHPLC Guard 3PK InfinityLab Poroshell 120 SB-C18 4.6×5mm 2.7 Micron

Mobile phases: A - Deionized water: Formic acid (99.9:0.1%). B - HPLC-grade MeCN: (Deionized water: Formic acid (99.9:0.1%)) (95:5%)

Gradient: from A - 99%, B - 1% to A - 1%, B - 99%

#### HPLC procedures

Instrument specifications:

Agilent 1260 Infinity systems equipped with DAD and mass-detector *General Parameters:* Temperature: 25 °C

##### HPLC procedure 1

Column: Chromatorex 18 SMB 100-5T 100A, 5 μm, 19 mm x 100mm with SiliaSphere C18 100A 5μm 100 A, 19mm x 10 mm

Detection: DAD – DAD1A 215 nm, DAD1B 254 nm. MSD – single quadrupole, AP-ESI Mobile phases: A - Deionized water (100%). B - HPLC-grade CH3CN (100%)

##### HPLC procedure 2

Column: Chromatorex 18 SMB 100-5T 100A, 5 μm, 19 mm x 100mm with SiliaSphere C18 100A 5μm 100 A, 19mm x 10 mm

Detection: DAD – DAD1A 215 nm, DAD1B 254 nm. MSD – single quadrupole, AP-ESI Mobile phases: A - Deionized water (100%). B - HPLC-grade MeOH (100%)

##### HPLC procedure 3

Column: Chromatorex 18 SMB 100-5T 100A, 5 μm, 19 mm x 100mm with SiliaSphere C18 100A 5μm 100 A, 19mm x 10 mm

Detection: DAD – DAD1A 200 nm, DAD1B 215 nm. MSD – single quadrupole, AP-ESI Mobile phases: A - Deionized water (100%). B - HPLC-grade CH3CN (100%)

##### HPLC procedure 4

Column: Chromatorex 18 SMB 100-5T 100A, 5 μm, 19 mm x 100mm with SiliaSphere C18 100A 5μm 100 A, 19mm x 10 mm

Detection: DAD – DAD1A 200 nm, DAD1B 254 nm. MSD – single quadrupole, AP-ESI Mobile phases: A - Deionized water (100%). B - HPLC-grade MeOH (100%)

##### HPLC procedure 5

Column: XBridge Shield RP18 OBD Column, 30*150 mm, 5μm Detection: UV - wavelength: 254 and 220 nm

Mobile Phases: A: Water (10 mmol/L NH_4_HCO_3_). B: ACN; Flow rate: 60 mL/min.

##### HPLC procedure 6

Column: conditions (Column: YMC-Actus Triart C18, 30*150 mm, 5μm Detection: UV - wavelength: 254 and 220 nm

Mobile Phases: A: Water (10 mmol/L NH_4_HCO_3_). B: ACN; Flow rate: 60 mL/min.

### General synthetic methods

Reagents and solvents were obtained from commercial suppliers and used without further purification, unless otherwise stated. Reactions were carried out under a positive atmosphere of nitrogen and monitored by thin layer chromatography (TLC) using shortwave UV light or by SHIMADZU LC/MS. Normal phase flash column chromatography was performed using silica gel 60 N (spherical, 75-150 μm). ^1^H NMR spectra were recorded on Bruker Avance 400 MHz spectrometer and were calibrated using residual non-deuterated solvent as the internal references (CDCl_3_: 7.26 ppm;; MeOH-D_4_: 3.31 ppm, acetone-D_6:_ 2.05 ppm; DMSO-D_6_: 2.50 ppm)^157^. The following abbreviations were used to explain NMR peak multiplicities: s = singlet, d = doublet, t = triplet, q = quartet, p = pentet, m = multiplet, br = broad.

### General amide coupling procedure

Amide coupling procedure was performed as previously described^158^. The -NH_2_ substrate, -COOH substrate, EDC, HOAt, NEt_3_, and DMF are added to a flame dried round bottom flask under nitrogen atmosphere and stirred for 12 h. The reaction mixture is then diluted with EtOAc, transferred to a separatory funnel, washed with H_2_O, dried over magnesium sulfate, and concentrated.

### Compound preparation

#### SH-9791

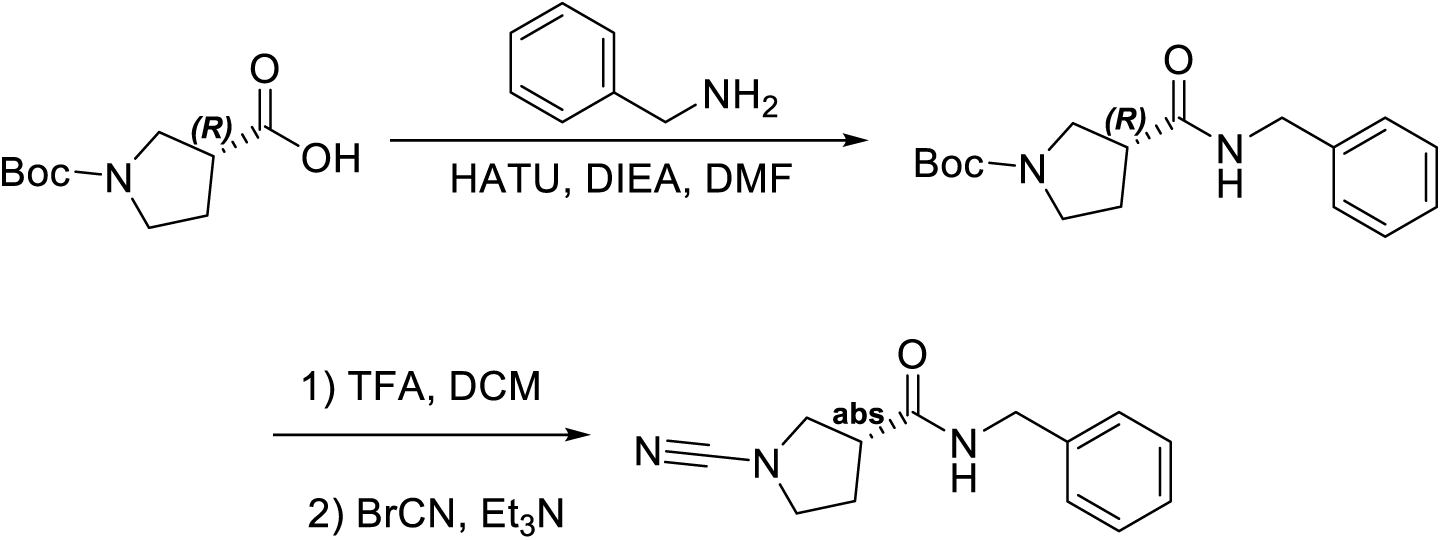

A solution of (3R)-1-(tert-butoxycarbonyl)pyrrolidine-3-carboxylic acid (400 mg, 1.86 mmol, 1 equiv) in DMF (5 mL) was treated with HATU (1060 mg, 2.79 mmol, 1.5 equiv) and DIEA (721 mg, 5.57 mmol, 3 equiv) at 0°C under nitrogen atmosphere followed by the addition of benzylamine (219 mg, 2.04 mmol, 1.1 equiv) in portions at room temperature^159^. The resulting mixture was stirred for 3 hrs at room temperature under nitrogen atmosphere. The resulting mixture was diluted with water (20 mL). The aqueous layer was extracted with CH_2_Cl_2_ (3×10 mL). The organic layers were concentrated under vacuum. The residue was purified by Prep-TLC (PE / EA 1:3) to afford tert-butyl (3R)-3-(benzylcarbamoyl)pyrrolidine-1-carboxylate (520 mg, 92%) as a yellow solid. LC-MS: (M-H)+ found 303.15.

A solution of tert-butyl (3R)-3-(benzylcarbamoyl)pyrrolidine-1-carboxylate (200 mg, 0.66 mmol, 1 equiv) in DCM (3 mL) was stirred at room temperature under nitrogen atmosphere, then the addition of TFA (1 mL) was administered dropwise^160^. The resulting mixture was stirred for 2 hrs under nitrogen atmosphere. The resulting mixture was concentrated under vacuum to afford (3R)- N-benzylpyrrolidine-3-carboxamide (200 mg, crude) and was used in the next step directly without further purification. LC-MS: (M+H)+ found 205.10.

A solution of (3R)-N-benzylpyrrolidine-3-carboxamide (200 mg, 0.98 mmol, 1 equiv) in DCM (3 mL) was treated with Et_3_N (495 mg, 4.89 mmol, 5 equiv) at 0°C under nitrogen atmosphere followed by the addition of BrCN (114.08 mg, 1.077 mmol, 1.1 equiv) in portions at 0°C^161^. The resulting mixture was stirred for 2 hrs at room temperature under nitrogen atmosphere. The mixture was diluted with CH_2_Cl_2_ saturated NaHCO_3_ and the aqueous layer was extracted with CH_2_Cl_2_ (3×10 mL). The organic layers were concentrated under vacuum. The crude product (150 mg) was purified by HPLC procedure 5 (Gradient: 10% B to 34% B) in 9 min to afford (3R)-N-benzyl-1-cyanopyrrolidine-3-carboxamide (40 mg, 18%) as a white solid. ^1^H NMR (400 MHz, Chloroform-*d*) *δ* 7.42 –7.31(m, 3H), 7.31 – 7.29 (m, 1H), 7.28 - 7.25 (m, 1H), 5.91 (s, 1H), 4.47 (d, J = 5.6 Hz, 2H), 3.71 – 3.57 (m, 3H), 3.45 – 3.40 (m, 1H), 2.92 – 2.85 (m, 1H), 2.25 – 2.10 (m, 2H). LC-MS: 228.00 (MH^+^)

**Figure 1.**
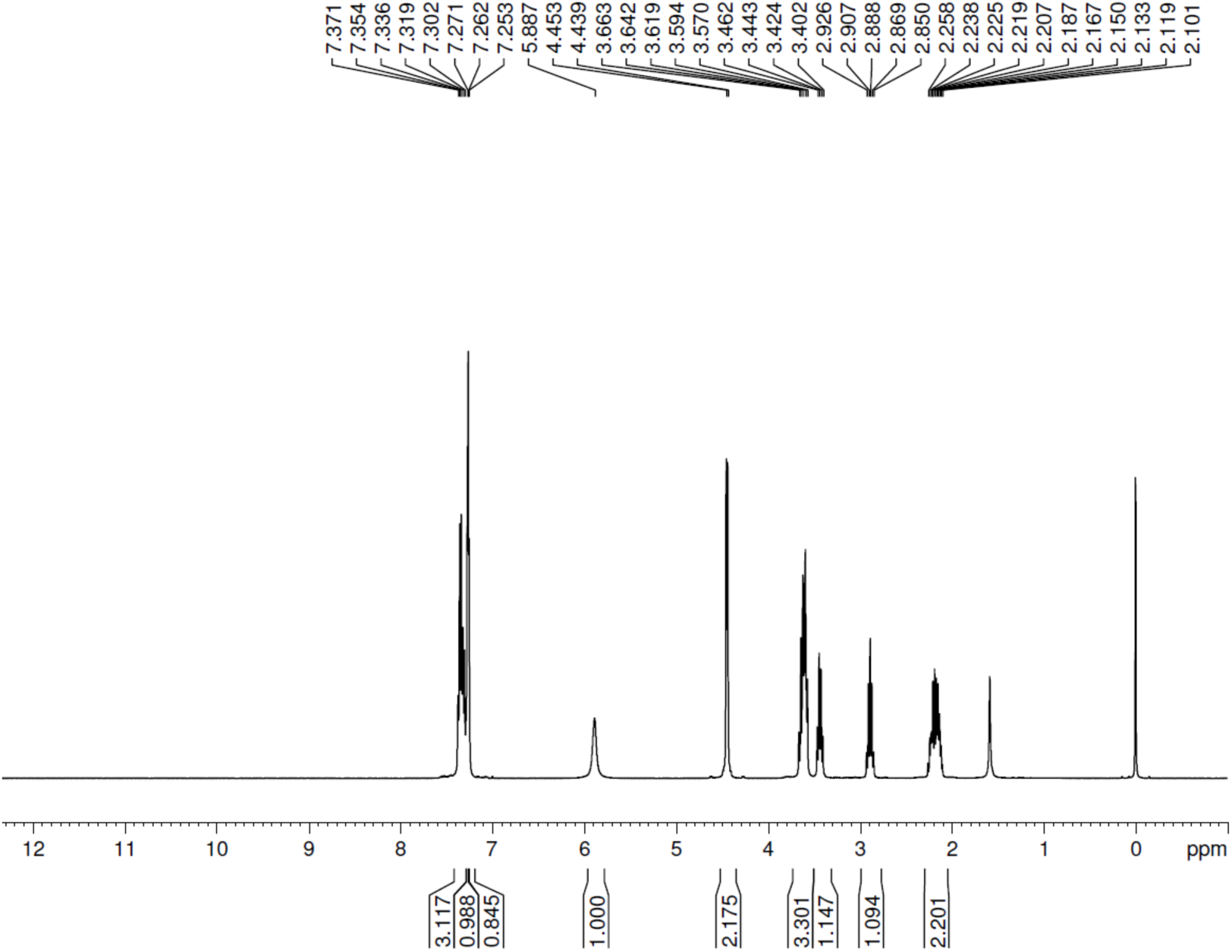
^1^H NMR spectrum of **SH-9791.**

#### SH-9857

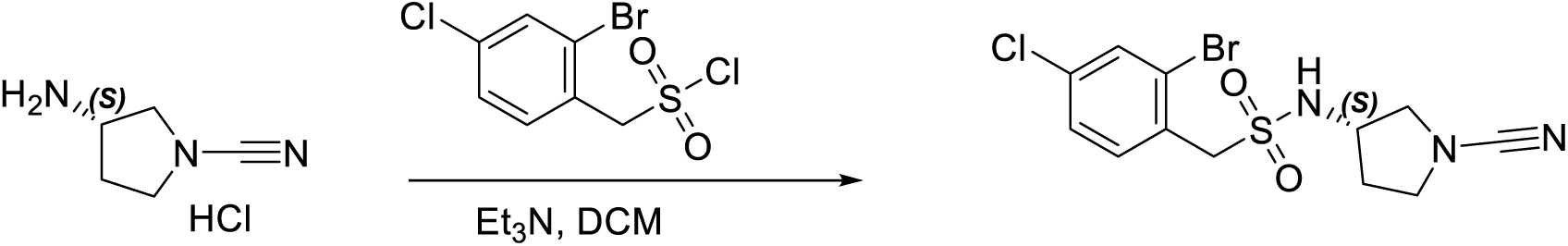

A solution of (3S)-3-aminopyrrolidine-1-carbonitrile (100 mg, 0.900 mmol, 1 equiv) in DCM (0.7 mL) was treated with Et_3_N (455 mg, 4.50 mmol, 5 equiv) for 5 min at room temperature under nitrogen atmosphere followed by the addition of (2-bromo-4-chlorophenyl) methanesulfonyl chloride (273.49 mg, 0.900 mmol, 1 equiv) in DCM (0.3 mL) dropwise at 0°C^160^. The resulting mixture was stirred for 2 hrs at room temperature under nitrogen atmosphere. The reaction was quenched with water at 0°C. The resulting mixture was extracted with CH_2_Cl_2_ (2 x 15 mL). The combined organic layers were washed with brine (1×10 mL) and dried over anhydrous Na_2_SO_4_. After filtration, the filtrate was concentrated under reduced pressure. The residue (100 mg) was purified by HPLC procedure 6, gradient: 33% B to 55% B in 9 min to afford 1-(2-bromo-4-chlorophenyl)-N-[(3S)-1-cyanopyrrolidin-3-yl] methanesulfonamide (39 mg, 11%) as a yellow solid. LCMS: (M+H)^+^ found 377.75. ^1^H NMR (300 MHz, Chloroform-*d*) δ 7.67 (d, J = 2.0 Hz, 1H), 7.53 (d, J = 8.3 Hz, 1H), 7.38 (d, J = 8.3, 1H), 4.52 (s, 3H), 3.84 (t, J = 5.7 Hz, 1H), 3.58 – 3.37 (m, 3H), 3.22 (d, J = 10.2, 1H), 2.15 – 2.01 (m, 1H), 1.87 – 1.75 (m, 1H).

**Figure 2.**
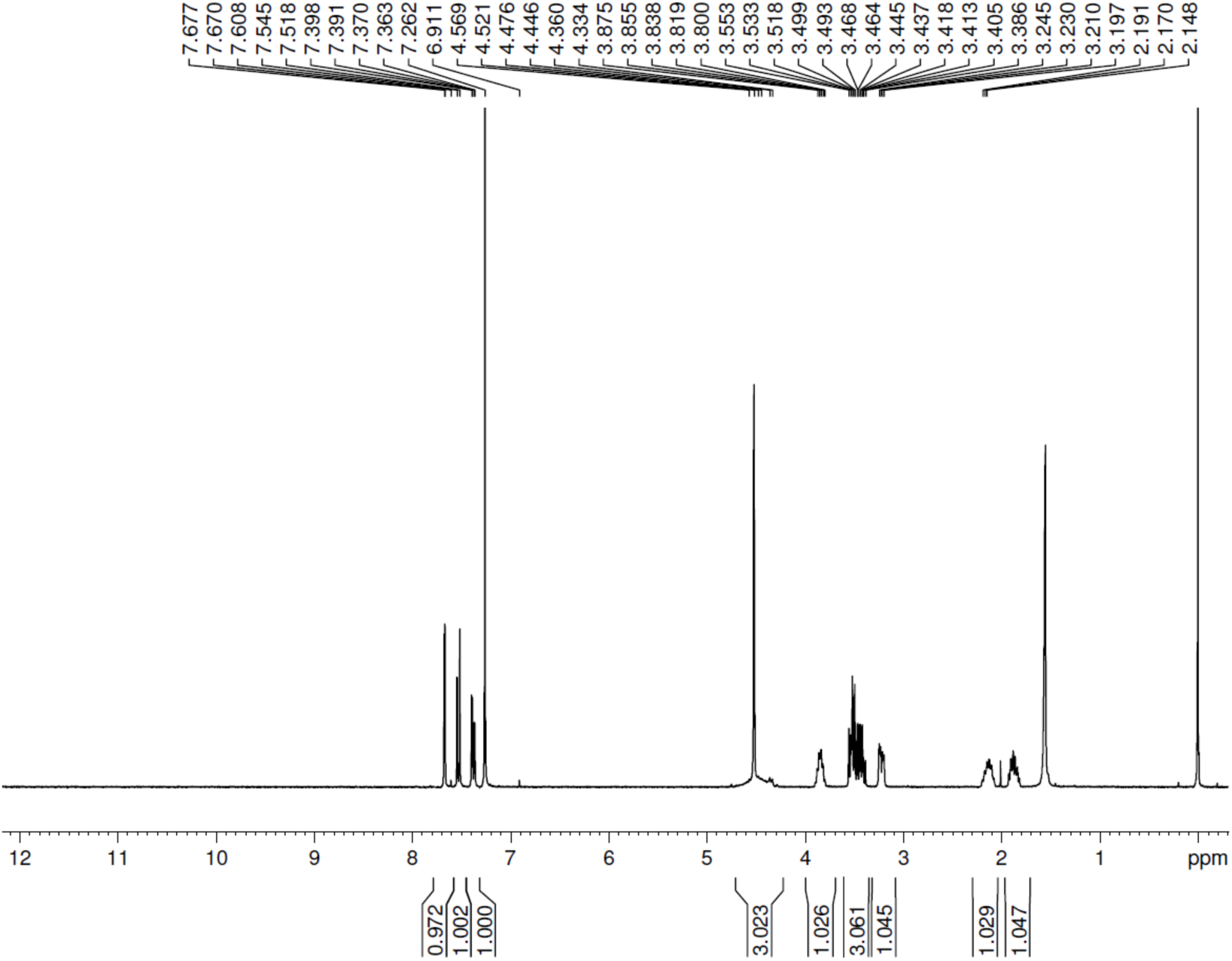
^1^H NMR spectrum of **SH-9857**.

#### SH-7346

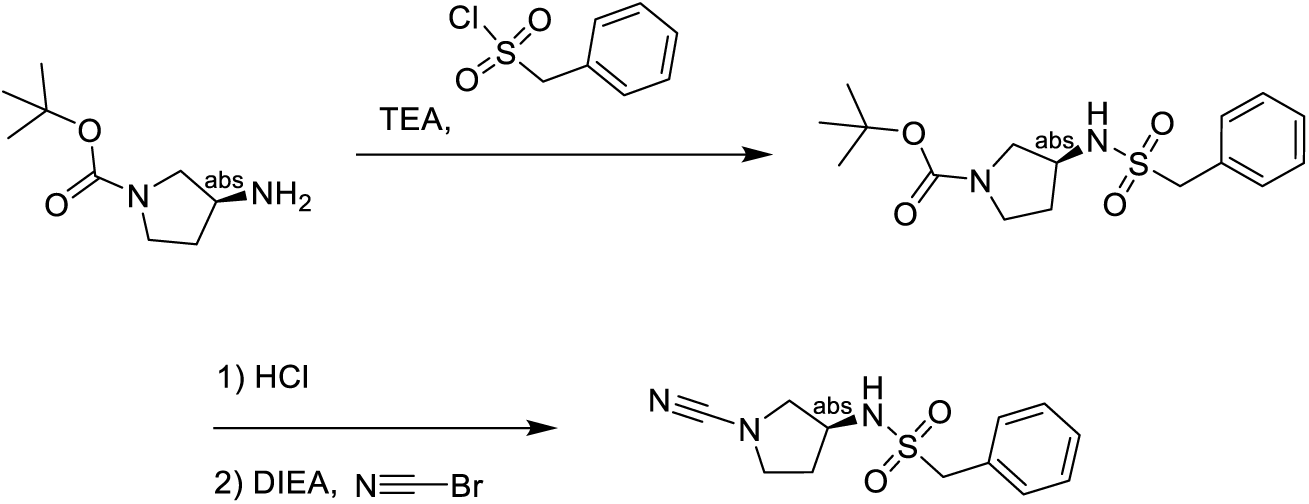

tert-butyl (S)-3-aminopyrrolidine-1-carboxylate (130 mg, 0.70 mmol, 1.00 equiv.) and phenylmethanesulfonyl chloride (130 mg, 1.05 mmol, 1.50 equiv.) were dissolved in ACN (5.00 mL), and then, TEA (0.31 mL, 2.10 mmol, 3.00 equiv.) was added to the solution^160^. The mixture was heated at 30 °C for 2 hrs. The mixture was concentrated in vacuo and the crude product was used into next step without further purification.

To a solution of tert-butyl (S)-3-((phenylmethyl)sulfonamido)pyrrolidine-1-carboxylate (crude) in DCM (0.50 mL), HCl-Dioxane (3.50 mL, 4.00 M) was added dropwise, then mixture was heated at 30 °C for 2 hrs^162^. The mixture was concentrated in vacuo and the crude product was used without further purification.

(S)-1-phenyl-N-(pyrrolidin-3-yl)methanesulfonamide (crude) was dissolved in DMSO (3.00 mL), and then DIEA (0.37 mL, 2.10 mmol, 3.00 equiv.) and cyanic bromide (147 mg, 1.40 mmol, 2.00 equiv.) were added to the solution^161^. The mixture was heated at 30 °C for 2 hrs. The mixture was purified by prep HPLC to give (S)-N-(1-cyanopyrrolidin-3-yl)-1-phenylmethanesulfonamide (40 mg, 0.15 mmol, 99%). ^1^H NMR (400 MHz, DMSO-*d*_6_) δ ppm 1.70 - 1.82 (m, 1 H) 1.98 - 2.10 (m, 1 H) 3.13 (dd, J=9.69, 4.82 Hz, 1 H) 3.35 - 3.39 (m, 1 H) 3.40 - 3.54 (m, 2 H) 3.88 (sxt, J=5.88 Hz, 1 H) 4.39 (s, 2 H) 7.31 - 7.43 (m, 5 H) 7.55 (br d, J=6.63 Hz, 1 H). MS-ESI, 266.1 (M+H^+^). HRMS-TOF, (M+H^+^), 266.0945.

**Figure 3.**
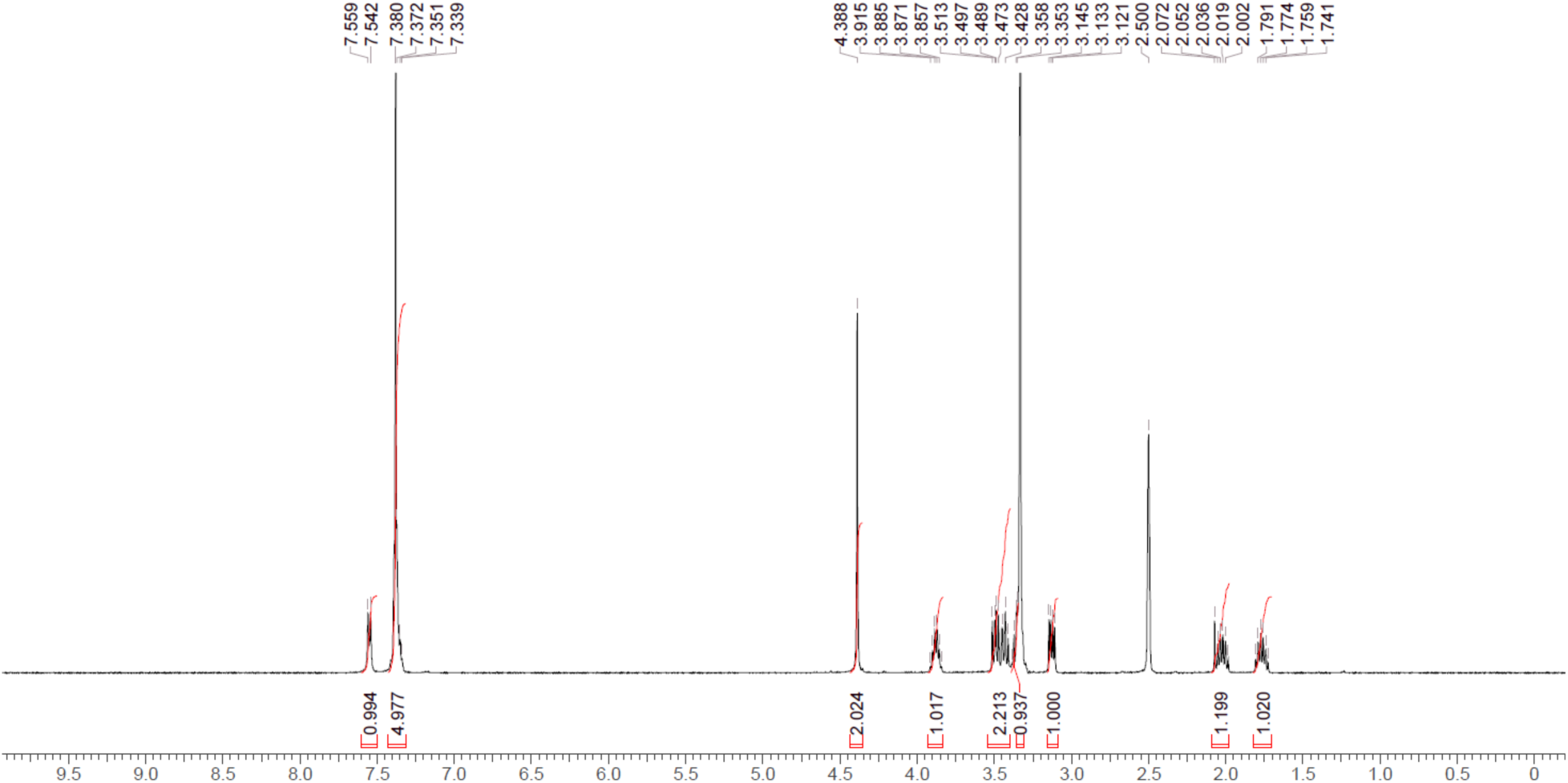
^1^H NMR spectrum of **SH-7346**.

#### SH-1696

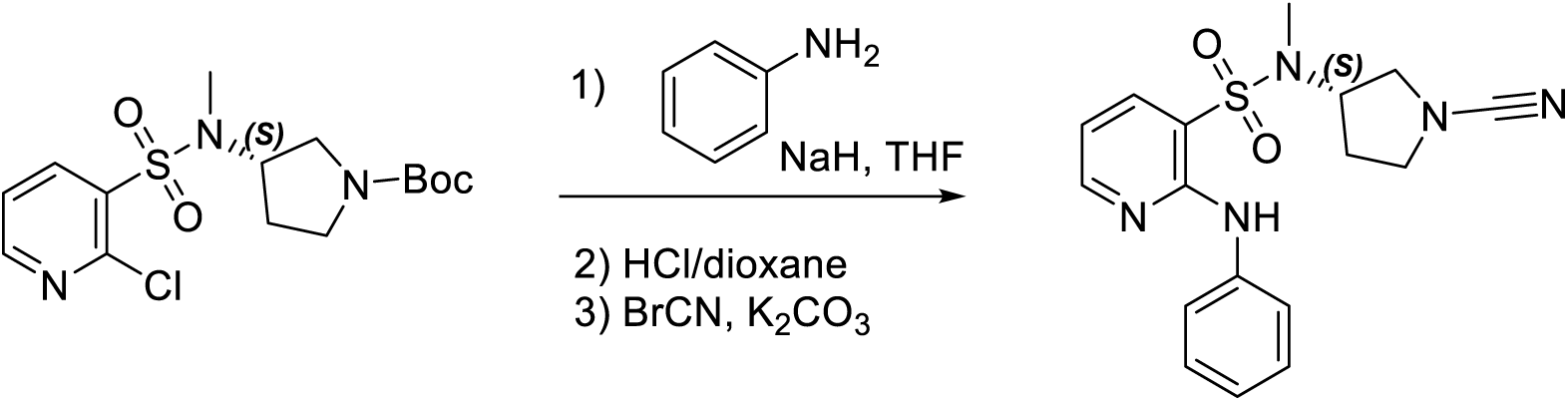

To a stirred solution of NaH (38 mg, 1.60 mmol, 2 equiv) and aniline (82 mg, 0.878 mmol, 1.1 equiv) in THF (5 mL) at 0°C, tert-butyl (3S)-3-(N-methyl2-chloropyridine-3-sulfonamido)pyrrolidine-1-carboxylate (300 mg, 0.798 mmol, 1 equiv) was added^163^. The resulting mixture was stirred for 12 hrs at 50°C then concentrated under reduced pressure. The residue was purified by reverse-phase flash chromatography with the following conditions: column, C18 silica gel; mobile phase, ACN in water, 0% to 100% gradient in 20 min; detector, UV 254 nm to afford tert-butyl (3S)-3-[N-methyl2-(phenylamino)pyridine-3-sulfonamido]pyrrolidine-1-carboxylate (80 mg, 23%) as a yellow oil. LC-MS: (M+H)^+^ found 433.

To a stirred solution of tert-butyl (3S)-3-[N-methyl2-(phenylamino)pyridine-3-sulfonamido]pyrrolidine-1-carboxylate (80 mg, 0.185 mmol, 1 equiv) in DCM (2 mL), HCl(gas) in 1,4-dioxane (0.5 mL, 16.456 mmol, 88.98 equiv) was added dropwise at room temperature^162^. The resulting mixture was stirred for 1hr and then concentrated under reduced pressure to afford crude N-methyl-2-(phenylamino)-N-[(3S)-pyrrolidin-3-yl]pyridine-3-sulfonamide (90 mg, crude) as a yellow solid. LC-MS: (M+H)^+^ found 333.

To a stirred solution of N-methyl-2-(phenylamino)-N-[(3S)-pyrrolidin-3-yl]pyridine-3-sulfonamide (90 mg, 0.27 mmol, 1 equiv) and BrCN (32 mg, 0.30 mmol, 1.1 equiv) in DMF (3 mL), K2CO3 (75 mg, 0.54 mmol, 2 equiv) was added at room temperature^161^. The resulting mixture was stirred for 1 hr then concentrated under reduced pressure and purified by HPLC procedure 5, gradient: 35% B to 57% B in 10 min to afford N-[(3S)-1-cyanopyrrolidin-3-yl]-N-methyl-2-(phenylamino)pyridine-3-sulfonamide (12 mg, 12%) as a yellow oil. ^1^H NMR (400 MHz, DMSO-d6) δ 8.54 (s, 1H), 8.41 (dd, 1H), 8.09 (dd, 1H), 7.63 – 7.57 (m, 2H), 7.39 – 7.30 (m, 2H), 7.12 – 7.04 (m, 1H), 7.03 – 6.96 (m, 1H), 4.79 – 4.67 (m, 1H), 3.46 – 3.40 (m, 2H), 3.30 – 3.27 (m, 1H), 3.27 – 3.23 (m, 1H), 2.77 (s, 3H), 1.95 – 1.77 (m, 2H). LC-MS: 358.0 (M+H^+^).

**Figure 4.**
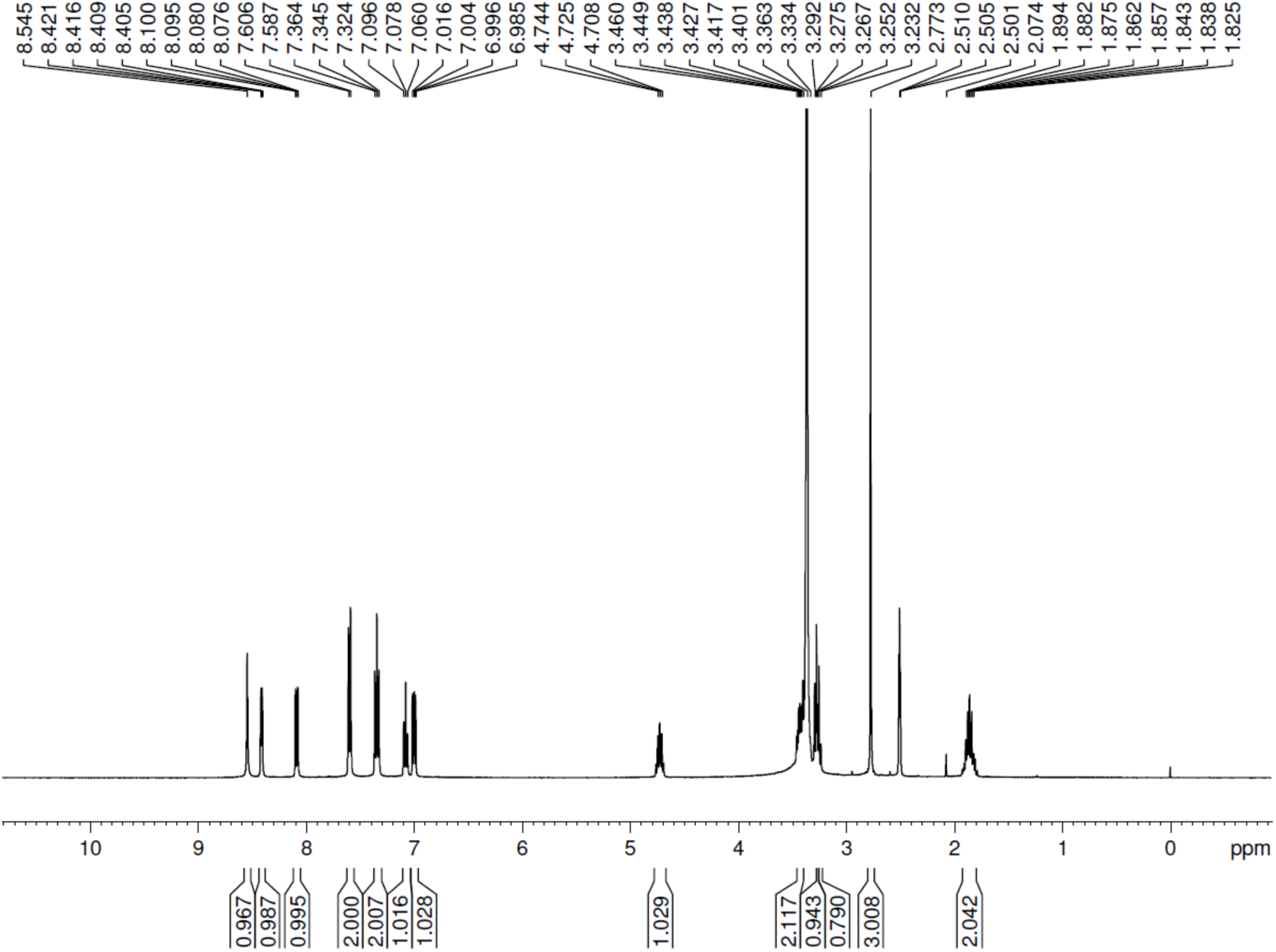
^1^H NMR spectrum of **SH-1696**.

#### SH-9840

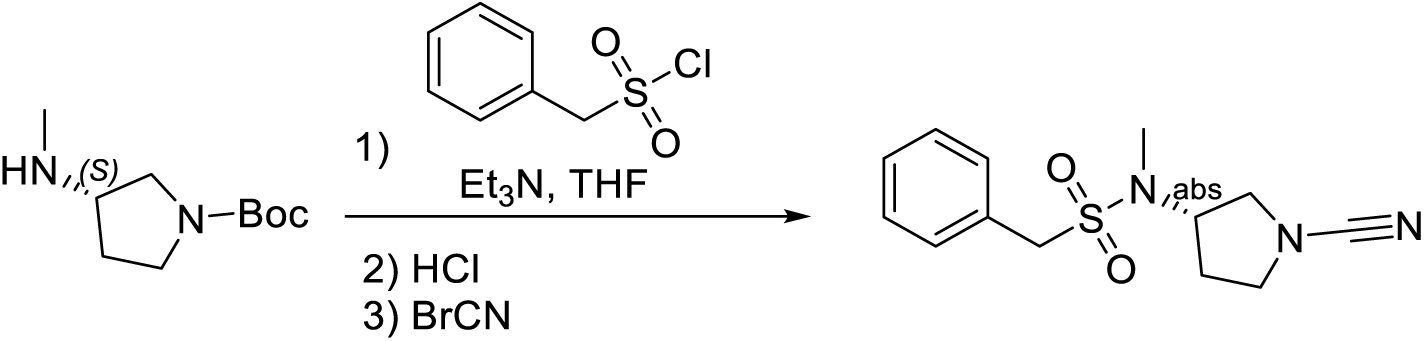

A round bottomed flask was charged with tert-butyl (3S)-3-(methylamino) pyrrolidine-1-carboxylate (500 mg, 2.50 mmol, 1 equiv), TEA (758 mg, 7.49 mmol, 3 equiv), and a stir bar. THF (10 mL) was added, the mixture was cooled to 0°C, then phenylmethanesulfonyl chloride (523 mg, 2.75 mmol, 1.1 equiv) was added^160^. The solution was stirred for 1 hr at room temperature before quenching the reaction by the addition of MeOH. The resulting mixture was concentrated in vacuo, and the residue was purified by silica gel column chromatography using PE/EA (2:1) to afford tert-butyl (3S)-3-(N-methylphenylmethanesulfonamido) pyrrolidine-1-carboxylate (480 mg, 54%) as off-white oil. LC-MS: (M+H)^+^ found: 355.20.

A round bottomed flask was charged with tert-butyl (3S)-3-(N-methylphenylmethanesulfonamido) pyrrolidine-1-carboxylate (480 mg, 1.35 mmol, 1 equiv) and a stir bar^162^. A solution of HCl in 1,4-dioxane (10 mL) was added, and the solution was stirred for 1 h at room temperature. The solution was concentrated in vacuo to afford crude N-methyl-1-phenyl-N-[(3S)-pyrrolidin-3-yl] methanesulfonamide (300 mg, 87%) as an off-white amorphous solid that was used without further purification. LC-MS: (M+H)^+^ found: 255.05.

A round bottomed flask was charged with N-methyl-1-phenyl-N-[(3S)-pyrrolidin-3-yl] methanesulfonamide (100 mg, 0.393 mmol, 1 equiv), TEA (119 mg, 1.18 mmol, 3 equiv), DCM (3 mL), and a stir bar^161^. The mixture was cooled to 0°C, cyanogen bromide (46 mg, 0.43 mmol, 1.1 equiv) was added, and the mixture stirred for 1 hr at room temperature. The mixture was diluted with water (30 mL), and the aqueous phase was extracted with DCM (30 mL) three times. The combined organic layers were washed with brine, dried over sodium sulfate, filtered, and concentrated in vacuo. The resulting crude material was purified by Prep-HPLC to give N-[(3S)-1-cyanopyrrolidin-3-yl]-N-methyl-1-phenylmethanesulfonamide N-[(3S)-1-cyanopyrrolidin-3-yl]-N-methyl-1-phenylmethanesulfonamide (54 mg, 49%) as an off-white amorphous solid. ^1^H NMR (400 MHz, DMSO-d6) δ 7.45 – 7.33 (m, 5H), 4.46 (s, 2H), 4.34 (p, J = 7.8 Hz, 1H), 3.46 (ddd, J = 9.2, 6.8, 5.5 Hz, 1H), 3.34 – 3.25 (m, 2H), 3.22 (dd, J = 9.7, 7.3 Hz, 1H), 2.68 (s, 3H), 1.98 – 1.84 (m, 2H). LC-MS: 280.05 (M+H)^+^

**Figure 5.**
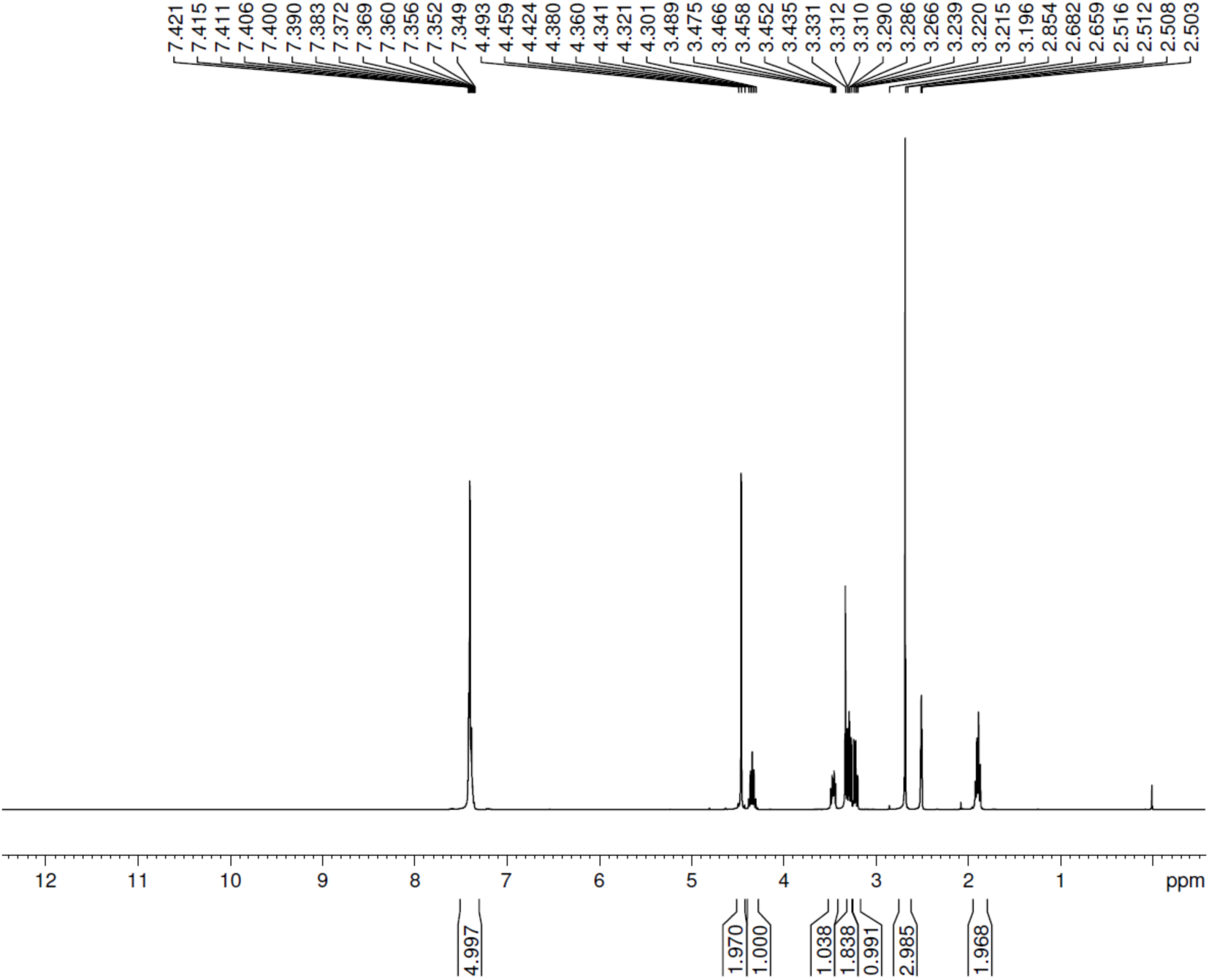
^1^H NMR spectrum of **SH-9840**.

#### SH-1688

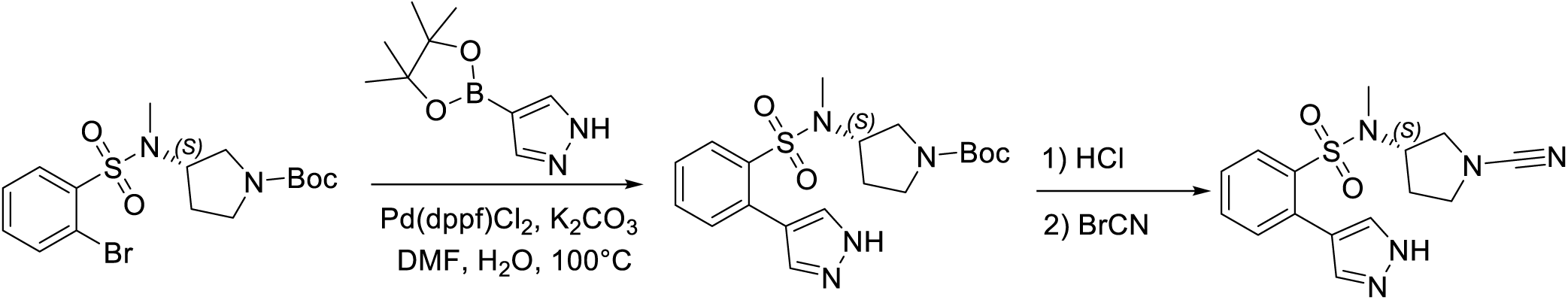

To a stirred solution of tert-butyl (3S)-3-(N-methyl2-bromobenzenesulfonamido)pyrrolidine-1-carboxylate (200 mg, 0.477 mmol, 1 equiv) and K_2_CO_3_ (198 mg, 1.43 mmol, 3 equiv) in DMF (2 mL) and H_2_O (0.4 mL) were added Pd(dppf)Cl_2_^164^. CH_2_Cl_2_ (38.9 mg, 0.048 mmol, 0.1 equiv) and 4-(4,4,5,5-tetramethyl-1,3,2-dioxaborolan-2-yl)-1H-pyrazole (111 mg, 0.572 mmol, 1.2 equiv) in portions at room temperature under nitrogen atmosphere. The resulting mixture was stirred for 2hrs at 80°C. The resulting mixture was extracted with EtOAc (3×3 mL) and concentrated under reduced pressure. The residue was purified by Prep-TLC (CH_2_Cl_2_ / MeOH 10:1) to afford tert-butyl (3S)-3-[N-methyl2-(1H-pyrazol-4-yl)benzenesulfonamido]pyrrolidine-1-carboxylate (110 mg, 56%) as a yellow solid.

To a stirred solution of tert-butyl (3S)-3-[N-methyl2-(1H-pyrazol-4- yl)benzenesulfonamido]pyrrolidine-1-carboxylate (110 mg, 0.271 mmol, 1 equiv) in 4 N HCl (gas) in 1,4-dioxane (2 mL) at room temperature^162^. The resulting mixture was stirred for 1 hr at room temperature and concentrated under reduced pressure to afford crude N-methyl-2-(1H-pyrazol-4-yl)-N-[(3S)-pyrrolidin-3-yl]benzenesulfonamide (100 mg) as a yellow solid.

To a stirred solution of N-methyl-2-(1H-pyrazol-4-yl)-N-[(3S)-pyrrolidin-3-yl]benzenesulfonamide (70 mg, 0.228 mmol, 1 equiv) and K_2_CO_3_ (94.73 mg, 0.684 mmol, 3 equiv) in DMF (0.7 mL) was added BrCN (26.6 mg, 0.251 mmol, 1.1 equiv) in portions at 0°C under nitrogen atmosphere^161^. The resulting mixture was stirred for 1hr at room temperature then concentrated under reduced pressure. The crude material was purified by Prep-HPLC (Column: XBridge Prep OBD C18 Column, 30*150 mm, 5μm; Mobile Phase A: Water (10 mmol/L NH_4_HCO_3_), Mobile Phase B: ACN; Flow rate: 60 mL/min; Gradient: 26% B to 45% B in 9 min) to afford N-[(3S)-1-cyanopyrrolidin-3-yl]-N-methyl-2-(1H-pyrazol-4-yl)benzenesulfonamide (13.9 mg, 18.30%) as a white solid. ^1^H NMR (400 MHz, Chloroform-*d*) *δ* 8.16 – 8.10(m, 1H), 7.85 (s, 2H), 7.61 (td, *J* = 7.6, 1.4 Hz, 1H), 7.48 (td, *J* = 7.7, 1.4 Hz, 1H), 7.42 – 7.36(m, 1H), 4.16 (p, *J* = 7.6 Hz, 1H), 3.43 – 3.37 (m, 1H), 3.27 – 3.22 (m, 1H), 3.12 – 3.08 (m, 1H), 2.99 – 2.95 (m, 1H), 2.40 (s, 3H), 1.80 – 1.75 (m, 2H). LC/MS: 332.00 (M+H^+^).

**Figure 6.**
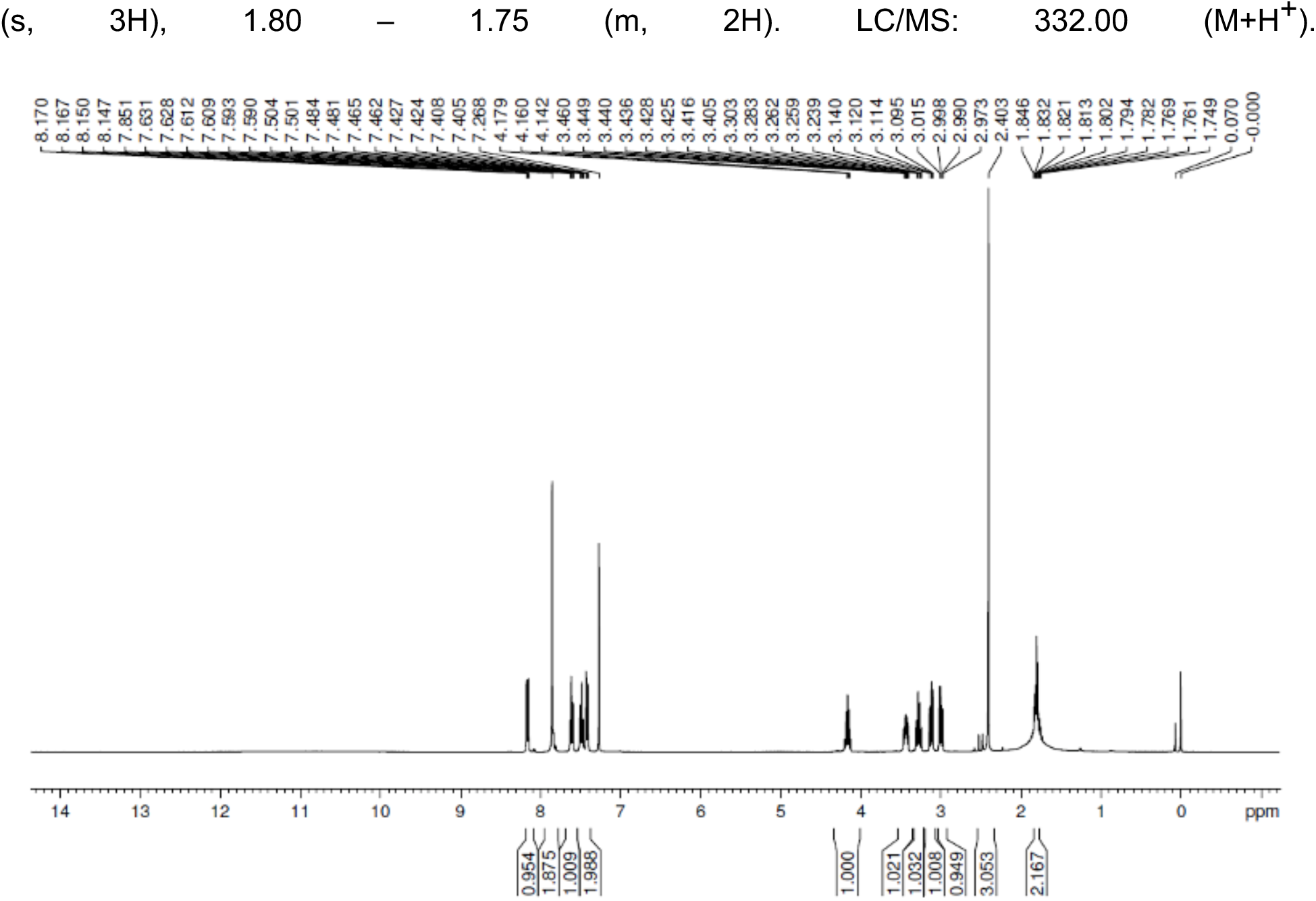
^1^H NMR spectrum of **SH-1688.**

#### SH-0105

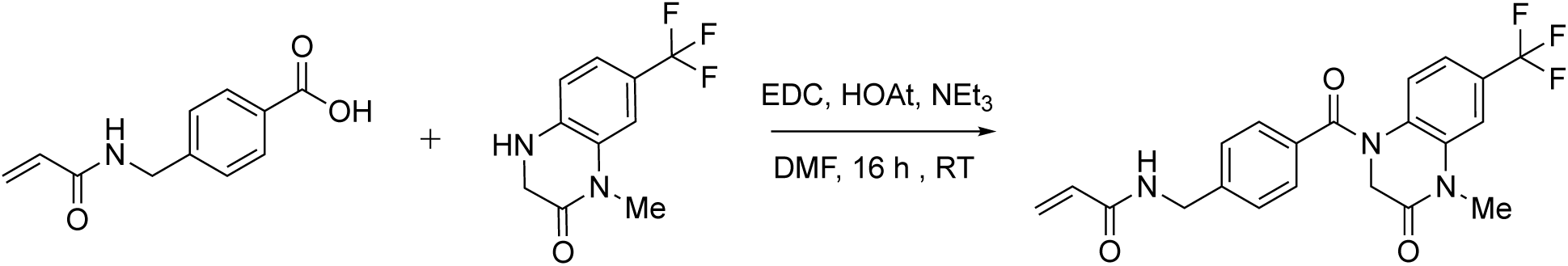

*N*-(4-(4-methyl-3-oxo-6-(trifluoromethyl)-1,2,3,4-tetrahydroquinoxaline-1-carbonyl)benzyl)acrylamide was obtained by using the aforementioned standard coupling conditions with 51 mg (0.22 mmol) of 1-methyl-7-(trifluoromethyl)-3,4-dihydroquinoxalin-2(1*H*)-one hydrochloride, 54 mg (0.263 mmol) of 4-[(prop-2-enamido)methyl]benzoic acid, 54 mg (0.348 mmol) of EDC, 33 mg (0.326 mmol) of Et3N, and 31.4 mg (0.231 mmol) of HOAt. Purified by HPLC procedure 4 (gradient: from A-85%: B-15% to A-35%: B-65%; Rf = 0.81; run time = 6.5 min). Yield: 40.0 mg (40 %). Beige solid. ^1^H NMR (400 MHz, Methanol-D_4_) δ 7.53 (s, 1H), 7.42 (d, *J* = 8.3 Hz, 2H), 7.32 (d, *J* = 8.5 Hz, 2H), 7.14 (d, *J* = 8.6 Hz, 1H), 6.98-6.96 (m, 1H), 5.69 (dd, *J* = 8.3, 3.7 Hz, 1H), 4.57 (s, 2H), 4.48 (s, 2H), 3.48 (s, 3H). EI MS m/z: pos. 418.0 (M+H^+^).

**Figure 7.**
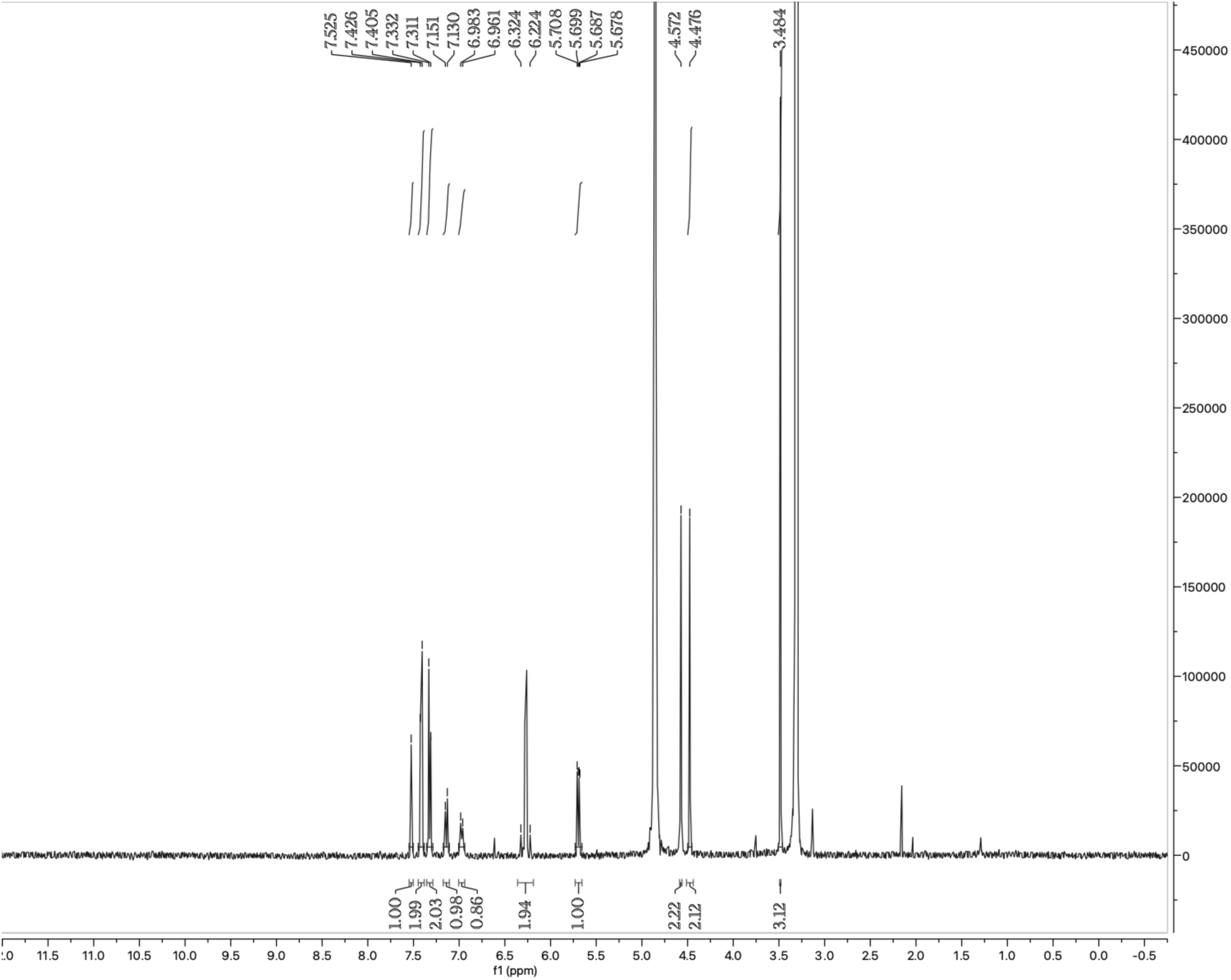
^1^H NMR spectrum of **SH-0105**.

#### SH-0073

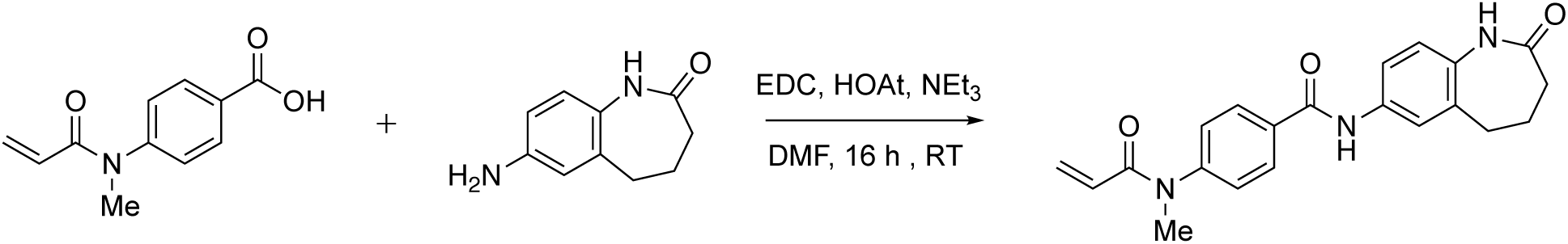

4-(N-methylprop-2-enamido)-N-(2-oxo-2,3,4,5-tetrahydro-1H-1-benzazepin-7-yl)benzamide was obtained by using the aforementioned standard coupling conditions with 53 mg (0.301 mmol) of 7-amino-2,3,4,5-tetrahydro-1H-1-benzazepin-2-one, 57 mg (0.278 mmol) of 4-(N-methylprop-2-enamido)benzoic acid, 49 mg (0.316 mmol) of EDC, and 43 mg (0.316 mmol) of HOAt. Purified by HPLC procedure 1 (gradient: from A-70%: B-30% to A-45%: B-55%). Yield: 56.9 mg (62 %). White powder. LCMS purity: 95.4 % (LCMS procedure 1, Rf = 0.51, run time = 2 min). ^1^H NMR (400 MHz, Methanol-D_4_) δ 8.03 (d, *J* = 8.6 Hz, 2H), 7.62-7.60 (m, 2H), 7.44-7.42 (m, 2H), 7.04 (d, *J* = 8.9 Hz, 1H), 6.31 (dd, *J* = 16.8, 2.1 Hz, 1H), 6.22-6.15 (m, 1H), 5.63 (dd, *J* = 10.2, 2.2 Hz, 1H), 3.39 (s, 3H), 2.81 (t, *J* = 6.9 Hz, 2H), 2.34-2.21 (m, 4H). EI MS m/z: pos. 364.1 (M+H^+^)

**Figure 8.**
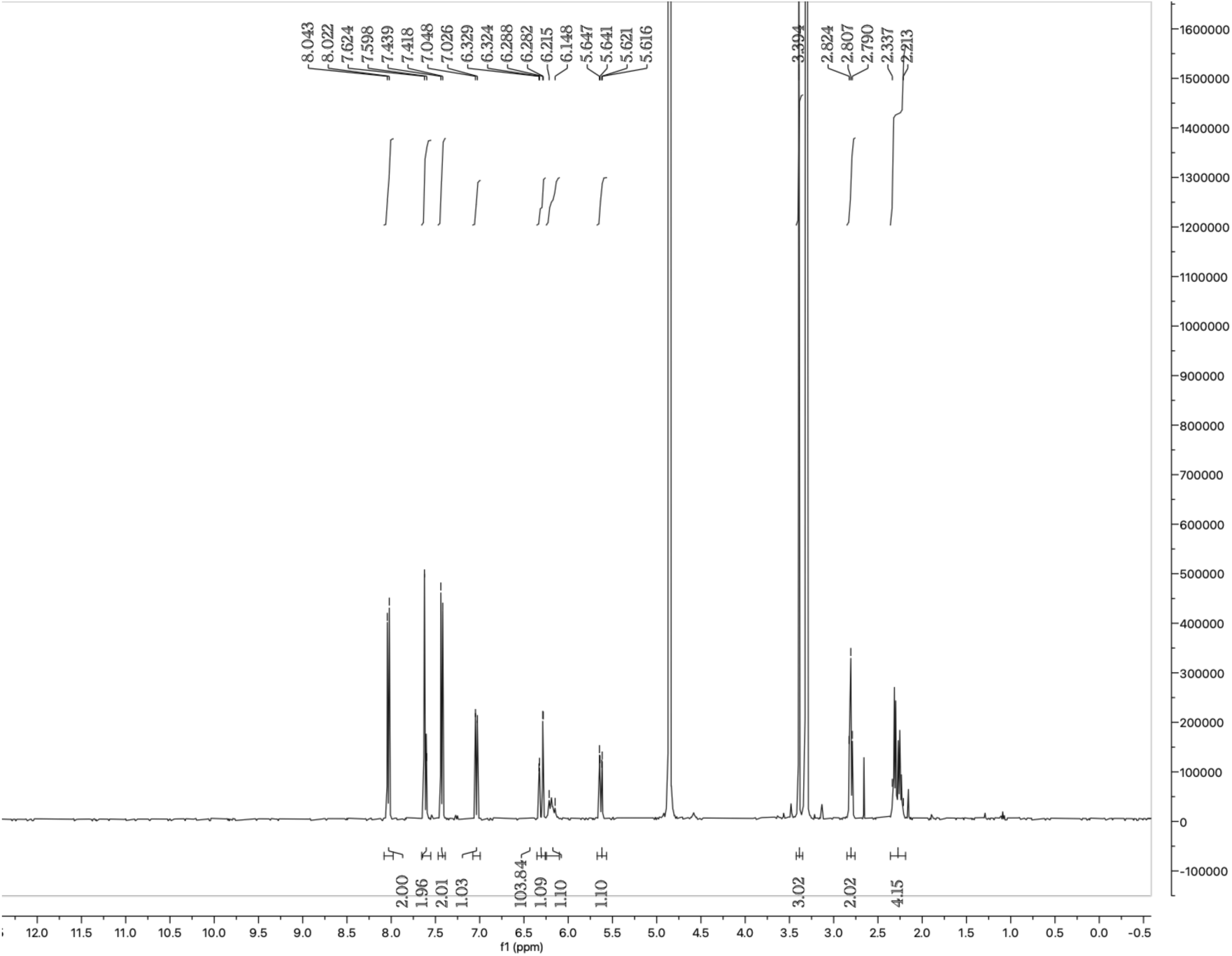
^1^H NMR spectrum of **SH-0073**.

#### SH-0059

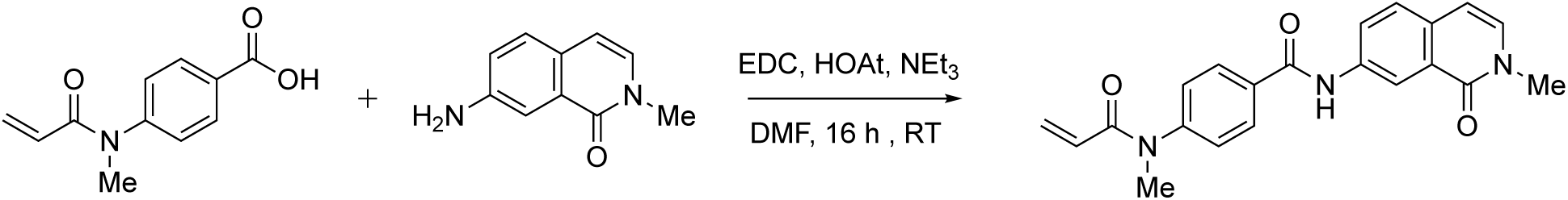

N-(2-methyl-1-oxo-1,2-dihydroisoquinolin-7-yl)-4-(N-methylprop-2-enamido)benzamide was obtained by using the aforementioned standard coupling conditions with 51 mg (0.293 mmol) of 7-amino-2-methyl-1,2-dihydroisoquinolin-1-one, 62 mg (0.302 mmol) of 4-(N-methylprop-2-enamido)benzoic acid, 59 mg (0.38 mmol) of EDC, and 41.9 mg (0.308 mmol) of HOAt. Purified by HPLC procedure 1 (gradient: from A-75%: B- 25% to A-50%: B-50%). Yield: 43.8 mg (44.1 %). Beige powder. LCMS purity: 98.5 % (LCMS procedure 1, Rf = 0.43, run time = 2 min). ^1^H NMR (400 MHz, Methanol-D_4_) δ 8.63 (d, *J* = 2.4 Hz, 1H), 8.14 (dd, *J* = 8.7, 2.3 Hz, 1H), 8.08 (d, *J* = 8.8 Hz, 2H), 7.68 (d, *J* = 8.6 Hz, 1H), 7.45 (d, *J* = 8.6 Hz, 2H), 7.33 (d, *J* = 7.3 Hz, 1H), 6.70 (d, *J* = 7.3 Hz, 1H), 6.31 (dd, *J* = 16.8, 2.2 Hz, 1H), 6.23-6.16 (m, 1H), 5.64 (dd, *J* = 10.1, 2.1 Hz, 1H), 3.63 (s, 3H), 3.40 (s, 3H). EI MS m/z: pos. 362.2 (M+H^+^).

**Figure 9.**
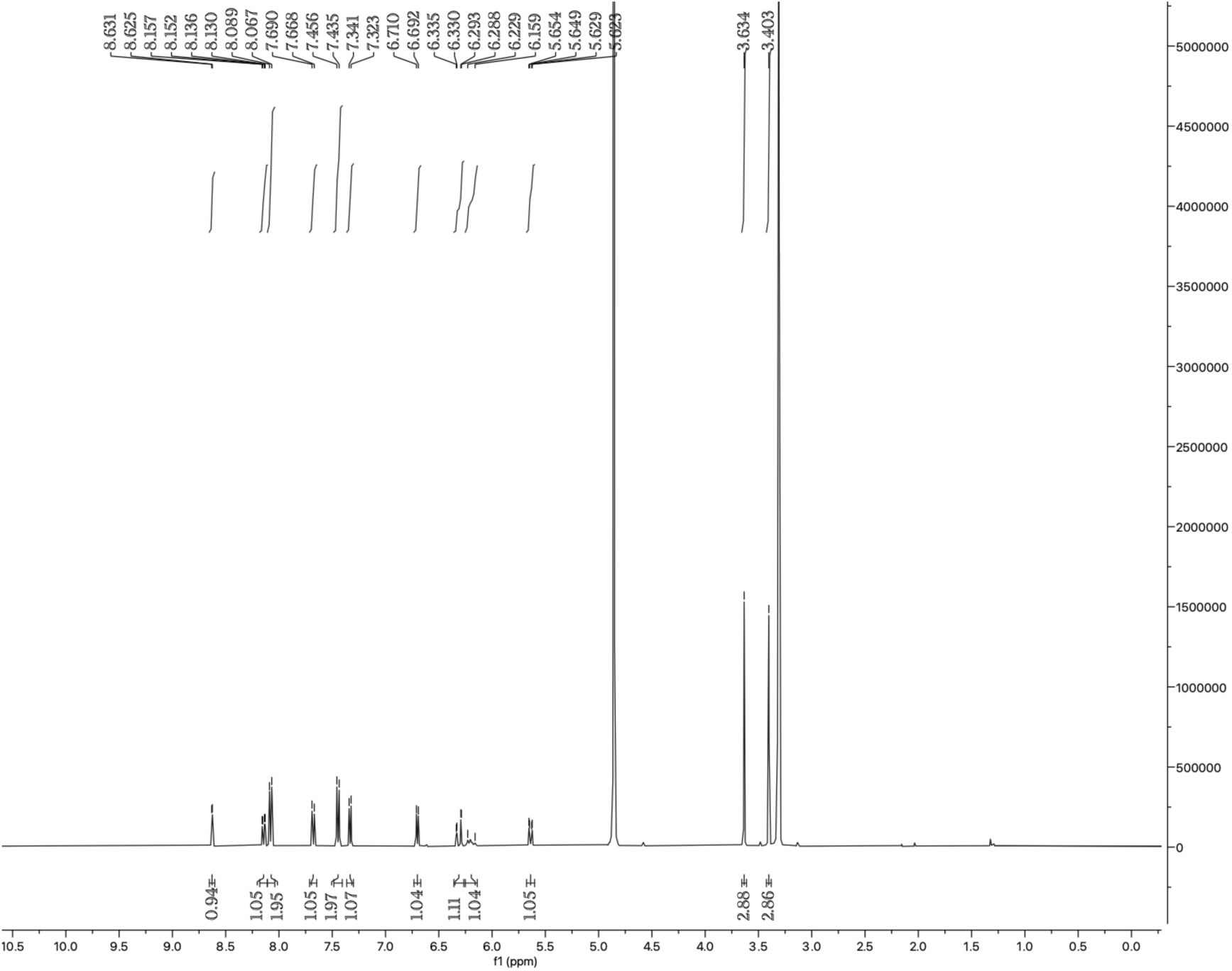
^1^H NMR spectrum of **SH-0059**.

#### SH-0096

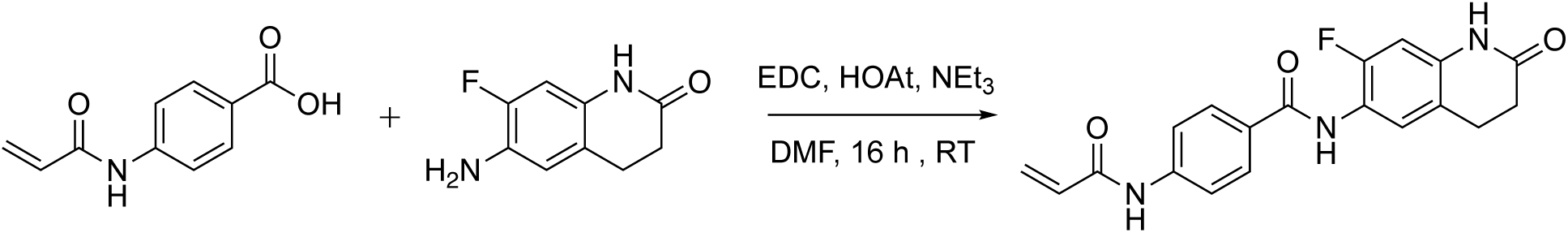

N-(7-fluoro-2-oxo-1,2,3,4-tetrahydroquinolin-6-yl)-4-(prop-2-enamido)benzamide was obtained by using the aforementioned standard coupling conditions with 44 mg (0.244 mmol) of 6-amino-7-fluoro-1,2,3,4-tetrahydroquinolin-2-one, 46 mg (0.241 mmol) of 4-(prop-2-enamido)benzoic acid, 40 mg (0.258 mmol) of EDC, and 34.9 mg (0.256 mmol) of HOAt. Purified by HPLC procedure 1 (gradient: from A-85%: B-15% to A-60%: B-40%). Yield: 24 mg (31.1 %). Beige powder. LCMS purity: 96.9 % (LCMS procedure 1, Rf = 0.51, run time = 2 min). ^1^H NMR (400 MHz, Methanol-D_4_) δ 7.95 (d, *J* = 8.9 Hz, 1H), 7.81 (d, *J* = 9.0 Hz, 1H), 7.47 (d, *J* = 7.9 Hz, 1H), 6.75 (d, *J* = 11.0 Hz, 1H), 6.50-6.38 (m, 1H), 5.82 (dd, *J* = 9.3, 2.6 Hz, 1H), 2.96 (t, *J* = 7.6 Hz, 1H), 2.60 (dd, *J* = 8.5, 6.6 Hz, 1H). EI MS m/z: pos. 354.0 (M+H^+^).

**Figure 10.**
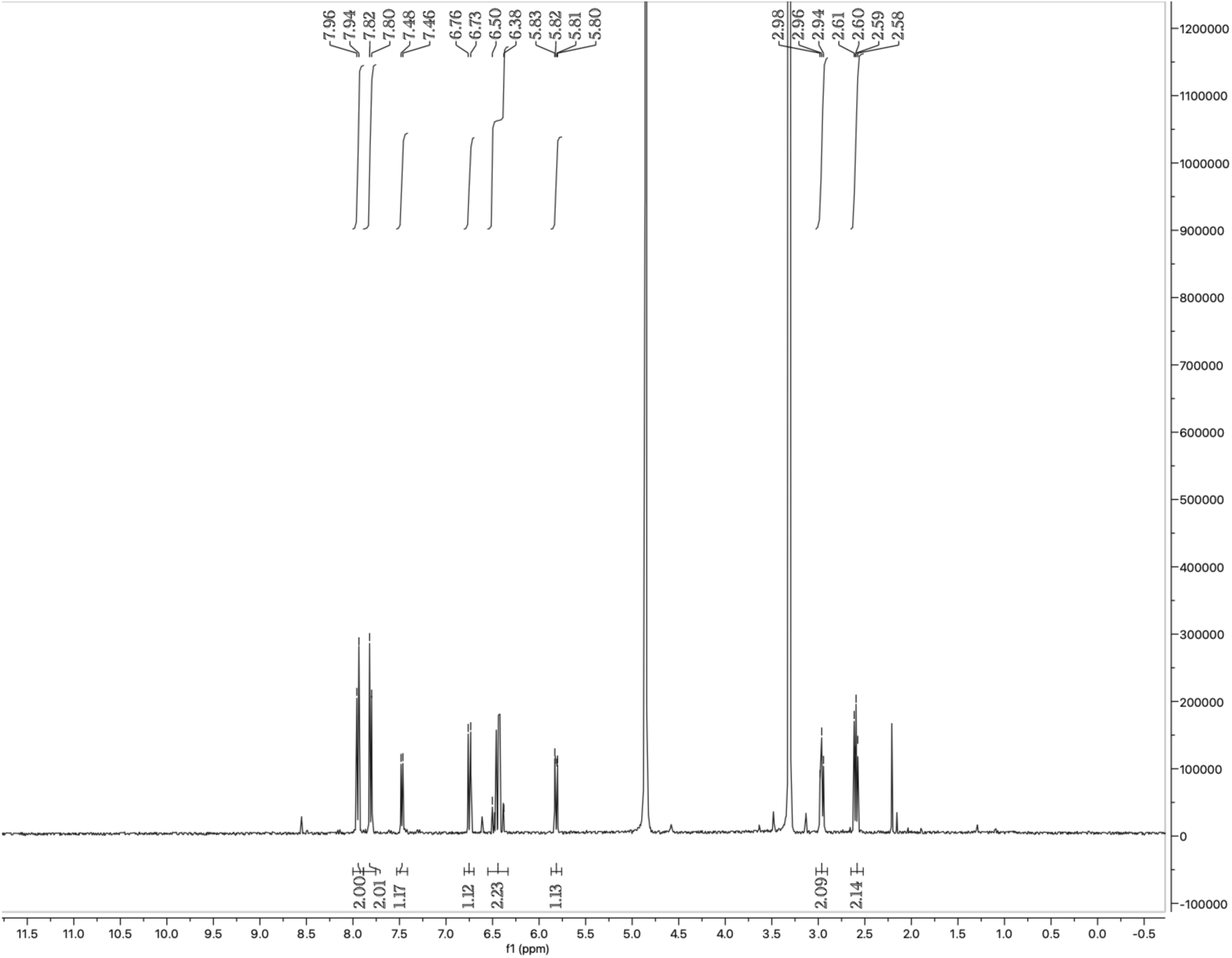
^1^H NMR spectrum of **SH-0096**.

#### SH-0002

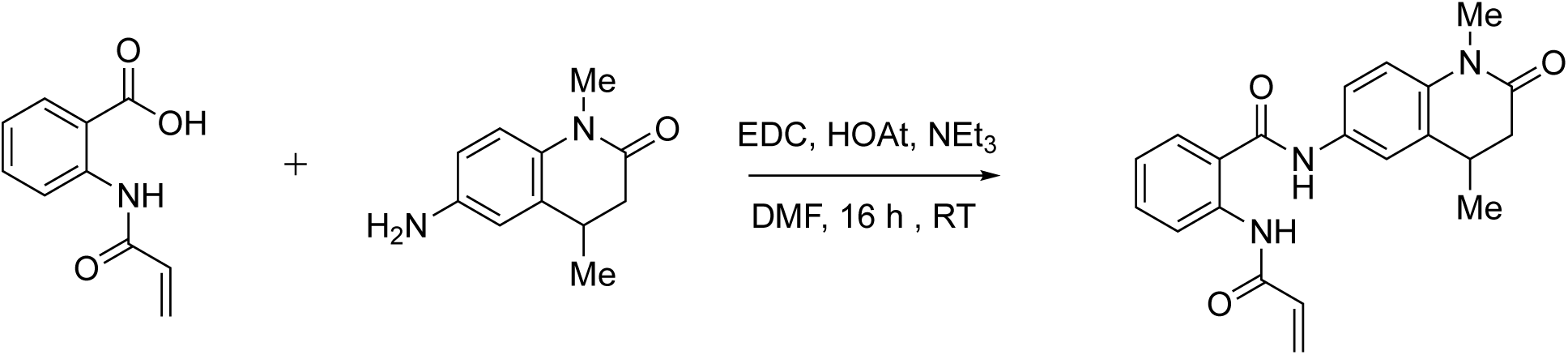

2-acrylamido-*N*-(1,4-dimethyl-2-oxo-1,2,3,4-tetrahydroquinolin-6-yl)benzamide was obtained by using the aforementioned standard coupling conditions with 56 mg (0.298 mmol) of 6-amino-1,4-dimethyl-3,4-dihydroquinolin-2(1*H*)-one, 63 mg (0.33 mmol) of 3-(prop-2-enamido)benzoic acid, 61 mg (0.393 mmol) of EDC, and 42.5 mg (0.312 mmol) of HOAt. Purified by HPLC procedure 2 (gradient: from A-60%: B-40% to A-10%: B-90%). Yield: 20.0 mg (19.0 %). Beige solid. ^1^H NMR (400 MHz, CDCl_3_) δ 8.21 (s, 1H), 8.17 (s, 1H), 7.94 (s, 1H), 7.72 (d, *J* = 8.0 Hz, 1H), 7.62 (d, *J* = 8.3 Hz, 1H), 7.54-7.44 (m, 2H), 7.41 (t, *J* = 7.9 Hz, 1H), 6.96 (d, *J* = 8.7 Hz, 1H), 6.45 (d, *J* = 16.8 Hz, 1H), 6.28 (dd, *J* = 16.9, 10.2 Hz, 1H), 5.79 (d, *J* = 11.5 Hz, 1H), 3.35 (s, 3H), 3.08-2.98 (m, 1H), 2.72 (dd, *J* = 15.8, 5.4 Hz, 1H), 2.44 (dd, *J* = 15.8, 7.5 Hz, 1H), 1.27 (d, *J* = 7.0 Hz, 3H). EI MS m/z: pos. 364.2 (M+H^+^).

**Figure 11.**
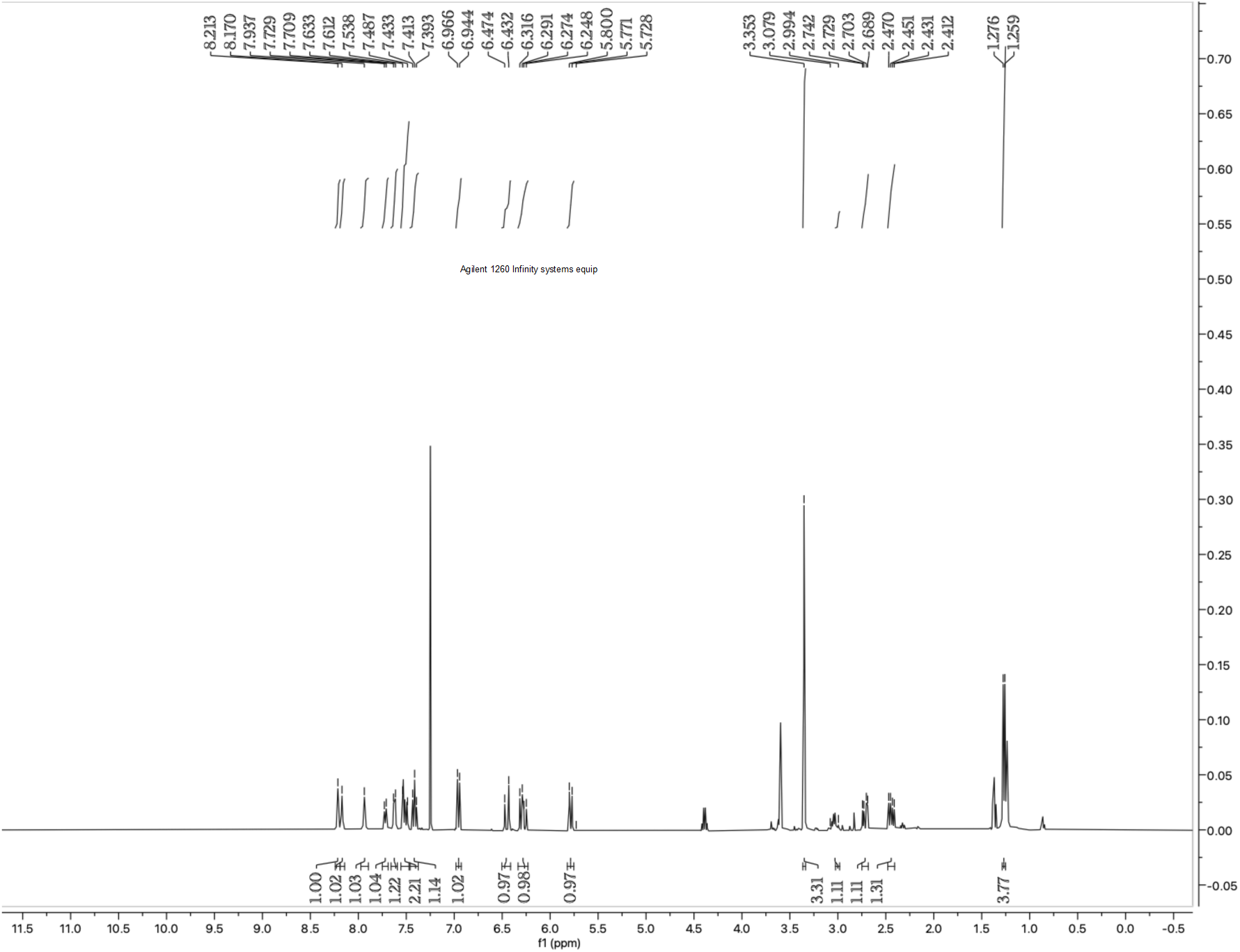
^1^H NMR spectrum of **SH-0002**.

#### SH-0042

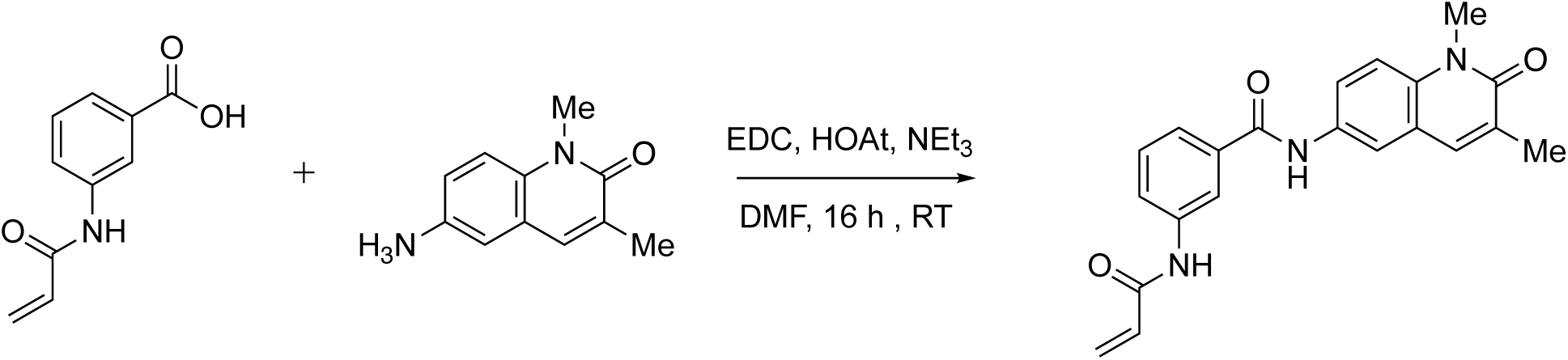

N-(1,3-dimethyl-2-oxo-1,2-dihydroquinolin-6-yl)-3-(prop-2-enamido)benzamide was obtained by using the aforementioned standard coupling conditions with 56 mg (0.298 mmol) of 6-amino-1,3-dimethyl-1,2-dihydroquinolin-2-one, 63 mg (0.33 mmol) of 3-(prop-2-enamido)benzoic acid, 61 mg (0.393 mmol) of EDC, and 42.5 mg (0.312 mmol) of HOAt. Purified by HPLC procedure 2 (gradient: from A-60%: B-40% to A-10%: B-90%). Yield: 9.6 mg (8.9 %). Beige powder. LCMS purity: 100 % (LCMS procedure 1, Rf = 0.6, run time = 2 min). ^1^H NMR (400 MHz, Methanol-D_4_) δ 8.24 (s, 1H), 8.06 (d, *J* = 2.4 Hz, 1H), 7.88 (dd, *J* = 9.1, 2.4 Hz, 1H), 7.82 (d, *J* = 8.1 Hz, 1H), 7.77 (s, 1H), 7.70 (d, *J* = 8.1 Hz, 1H), 7.57 (d, *J* = 9.1 Hz, 1H), 7.50 (t, *J* = 7.9 Hz, 1H), 6.50-6.38 (m, 2H), 5.82 (dd, *J* = 9.2, 2.7 Hz, 1H), 3.79 (s, 4H), 2.24 (s, 3H). EI MS m/z: pos. 362.2 (M+H^+^)

**Figure 12.**
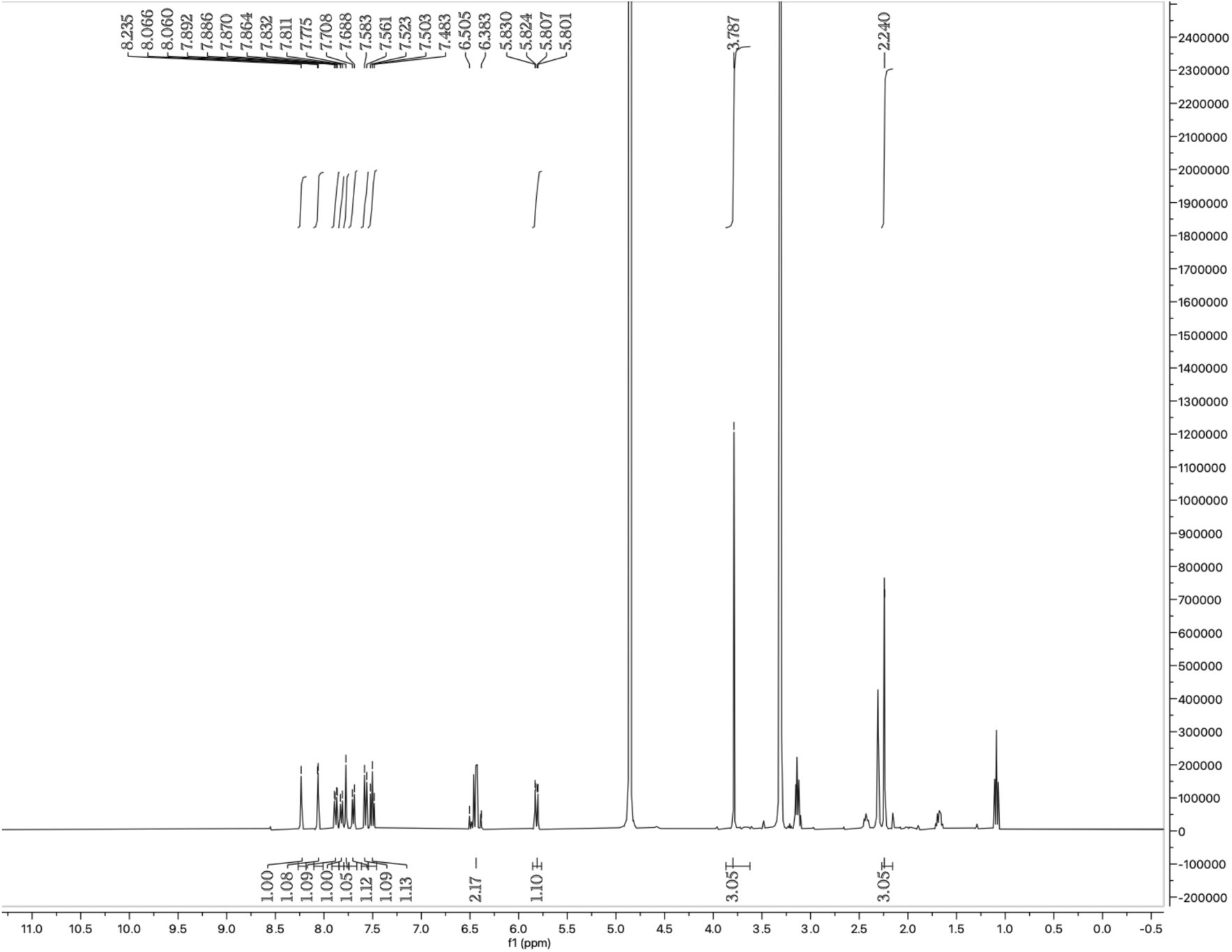
^1^H NMR spectrum of **SH-0042**.

#### SH-0029

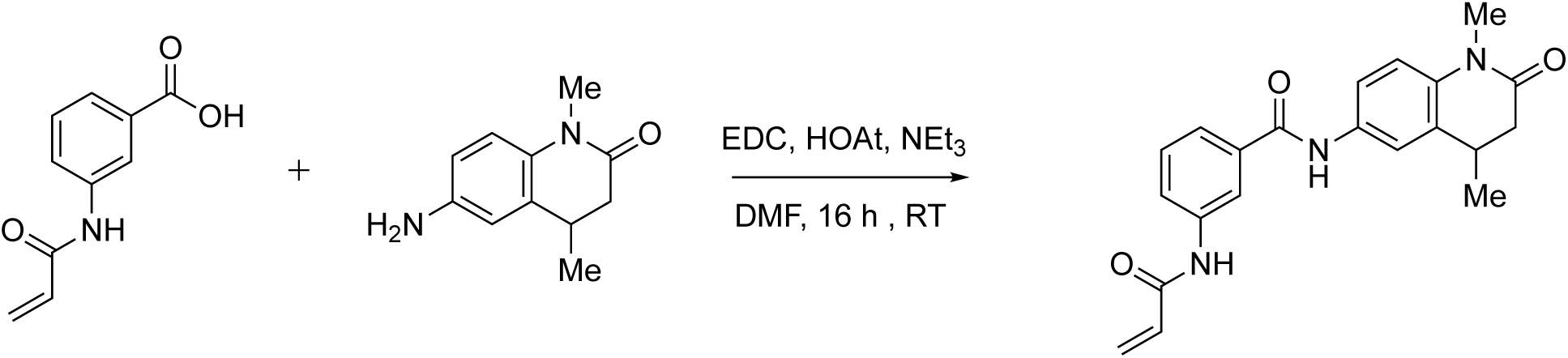

3-acrylamido-*N*-(1,4-dimethyl-2-oxo-1,2,3,4-tetrahydroquinolin-6-yl)benzamide was obtained by using the aforementioned standard coupling conditions with 56 mg (0.298 mmol) of 6-amino-1,4-dimethyl-3,4-dihydroquinolin-2(1*H*)-one, 63 mg (0.33 mmol) of 3-(prop-2-enamido)benzoic acid, 61 mg (0.393 mmol) of EDC, and 42.5 mg (0.312 mmol) of HOAt. Purified by HPLC procedure 2 (gradient: from A-60%: B-40% to A-10%: B-90%). Yield: 48.0 mg (45.0 %). Beige solid. ^1^H NMR (400 MHz, Methanol-*D*_4_) δ 8.17 (s, 1H), 7.78 (dm, *J* = 8.1 Hz, 1H), 7.64-7.59 (m, 3H), 7.46 (t, *J* = 7.9 Hz, 1H), 7.12 (d, *J* = 9.4 Hz, 1H), 6.47-6.35 (m, 2H), 5.78 (dd, *J* = 9.2, 2.7 Hz, 1H), 3.35 (s, 3H), 3.35 (s, 3H), 3.06 (h, *J* = 7.0 Hz, 1H), 2.72 (dd, *J* = 15.9, 5.5 Hz, 1H), 2.42 (dd, *J* = 15.9, 7.0 Hz, 1H). EI MS m/z: pos. 364.2 (M+H^+^).

**Figure 13.**
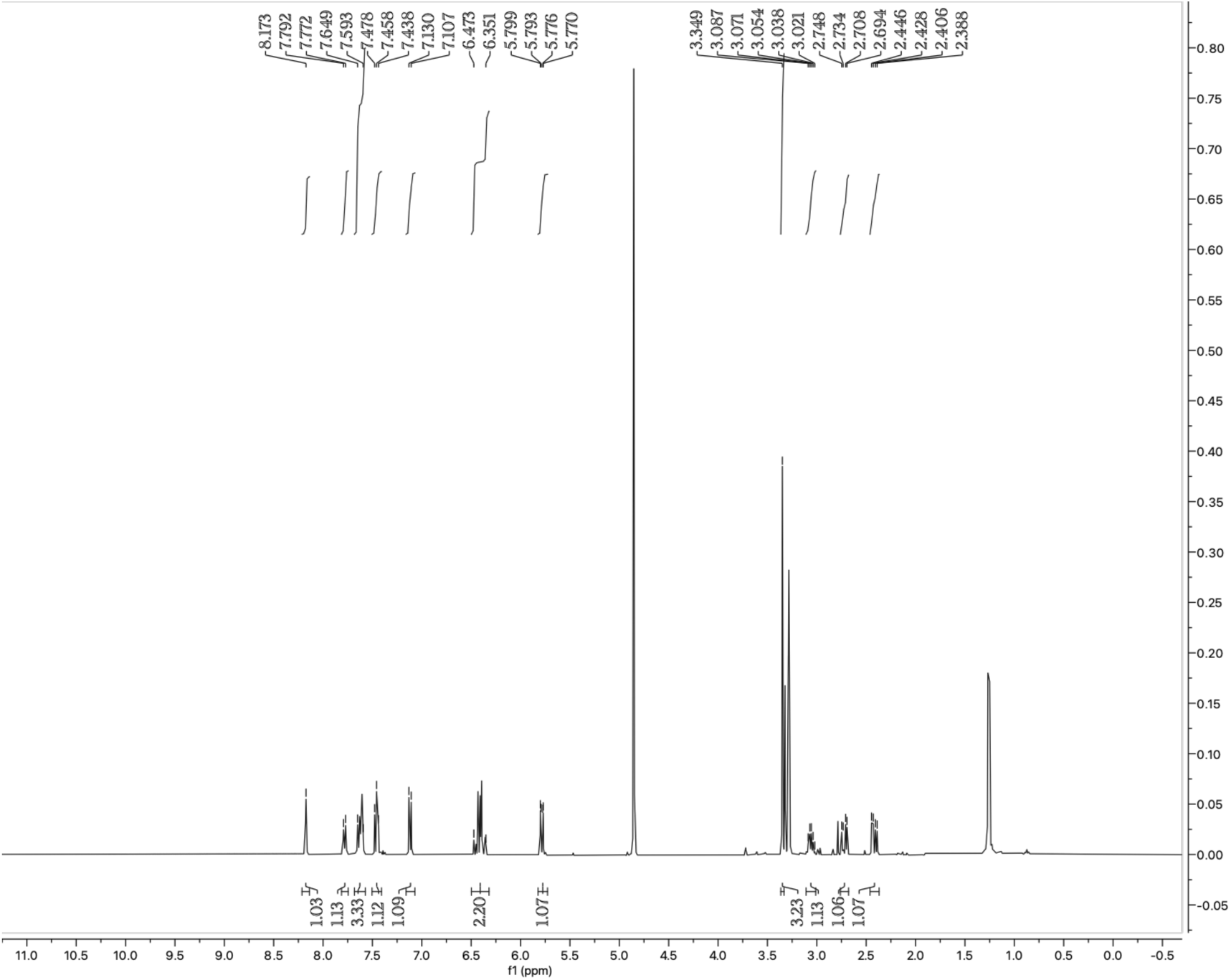
^1^H NMR spectrum of **SH-0029**.

#### SH-0029-DTB

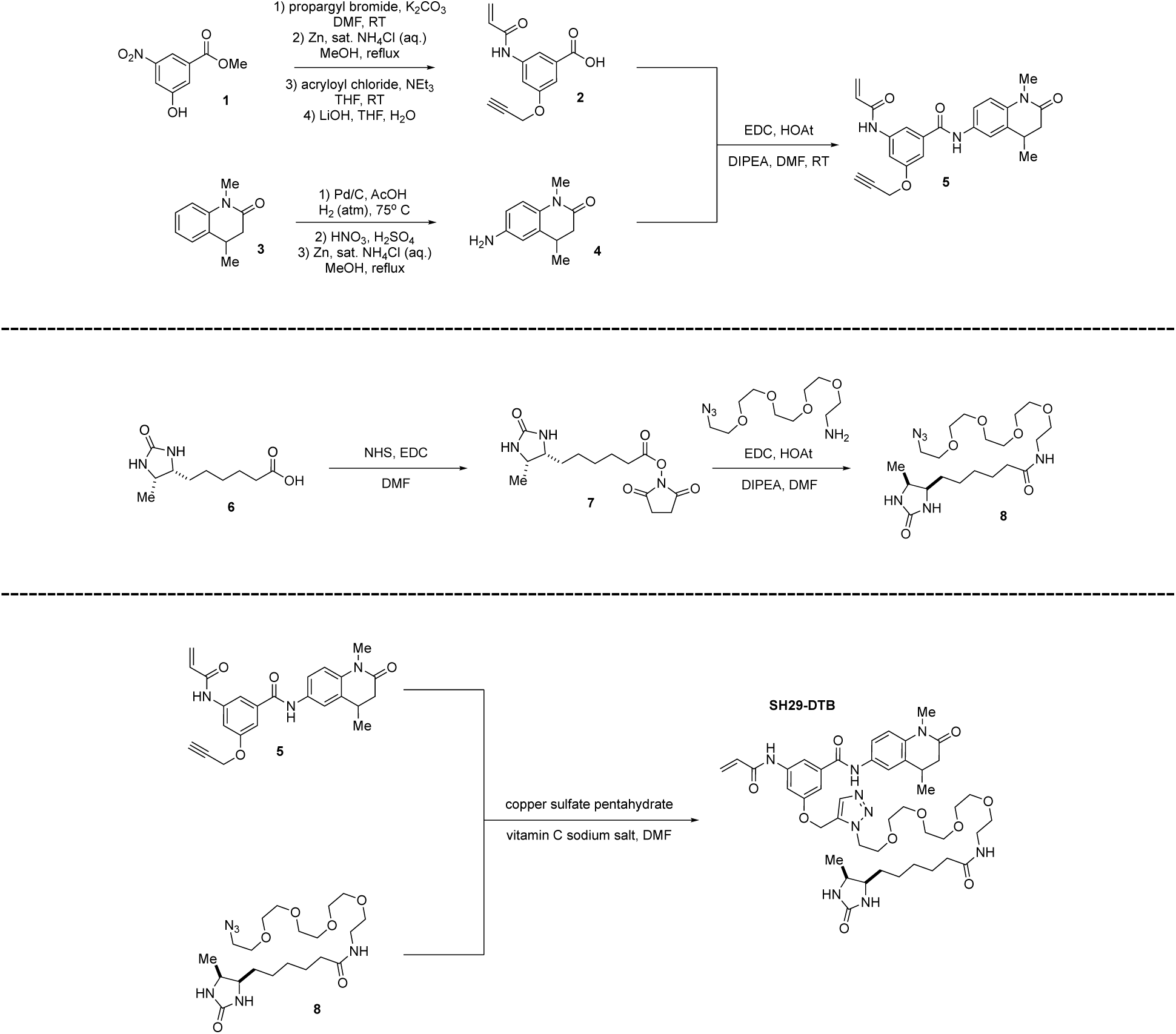

##### Synthesis of 2

To a flame-dried round bottom flask, methyl 3-hydroxy-5-nitrobenzoate (5.00 g, 25.4 mmol), potassium carbonate (4.20 g, 30.5 mmol), and propargyl bromide (1.68 g, 30.5 mmol) in DMF (20 mL) stirred at RT for 16 h. The mixture was diluted with diethyl ether (100 mL) and water (100 mL) and then transferred to a separatory funnel. The organic layer was separated, dried with magnesium sulfate, concentrated, and used in the next step without further purification to afford methyl 3-nitro-5-(prop-2-yn-1-yloxy)benzoate.

The isolated alkyne intermediate (1.00 g, 4.25 mmol) was suspended in MeOH (30 mL) and stirred at RT, then aqueous sat. NH_4_Cl (10 mL) and Zn metal (1.00 g, 15.4 mmol) were added portion-wise. The mixture was raised to reflux for 1 h, cooled to RT, and filtered through celite. The filtrate was concentrated to remove methanol then redissolved in EtOAc. The mixture was transferred to a separatory funnel and washed with sat. NaHCO_3_, dried with magnesium sulfate, concentrated, and purified via column chromatography to afford methyl 3-amino-5-(prop-2-yn-1-yloxy)benzoate.

In a flame-dried round bottom flask, the amine intermediate (1.00 g, 4.88 mmol) was dissolved in THF and cooled to 0° C. Acryloyl chloride (0.530 g, 5.86 mmol) and NEt_3_ (0.591 g, 5.86 mmol) were added dropwise. The reaction mixture was stirred for 3 h, quenched with 10 mL of DI water, and then transferred to a separatory funnel. The aqueous layer was extracted three times with EtOAc, dried with magnesium sulfate, concentrated, and purified via column chromatography to afford methyl 3-acrylamido-5-(prop-2-yn-1-yloxy)benzoate.

Methyl 3-acrylamido-5-(prop-2-yn-1-yloxy)benzoate (1.00 g, 3.86 mmol) was dissolved in THF/H_2_O (10:1, 11 mL), then LiOH monohydrate (0.195 g, 4.63 mmol) was added portion wise. The reaction mixture was stirred for 12 h at RT, then quenched with 20 mL of 1M HCl. The solution was transferred to a separatory funnel, extracted 3xEtOAc, dried with magnesium sulfate, concentrated, and purified via column chromatography to afford compound **2**.

##### Synthesis of 4

1,4-dimethylquinolin-2(1*H*)-one (**2**) (2.00 g, 11.7 mmol) was dissolved in 50.0 mL of AcOH, then Pd/C (cat.) was added, and the atmosphere was replaced with hydrogen gas. The reaction mixture was raised to 75° C, and then stirred for 16 h. The crude mixture was cooled to RT and filtered through celite. The filtrate was then poured into ice-water, and the product was collected via filtration and used in the next step without further purification.

1,4-dimethyl-3,4-dihydroquinolin-2(1*H*)-one (1.00 g, 5.71 mmol) was dissolved in H_2_SO_4_ (10 mL) and cooled to 0° C. Fuming nitric acid (5.71 mmol) was added dropwise, then the mixture was allowed to warm to RT and stirred for 3 h. The crude mixture was poured into a 2:1 ice-water mixture and extracted with EtOAc, dried with magnesium sulfate, concentrated, and purified via column chromatography to afford 1,4-dimethyl-6-nitro-3,4-dihydroquinolin-2(1H)-one.

1,4-dimethyl-6-nitro-3,4-dihydroquinolin-2(1*H*)-one (1.00 g, 4.59 mmol) was suspended in MeOH (70 mL). Then, aqueous sat. NH_4_Cl (20 mL) and Zn metal (1.00 g, 15.4 mmol) were added portion-wise. The mixture was raised to reflux for 1 h, cooled to RT, and filtered through celite. The filtrate was concentrated to remove methanol and then redissolved in EtOAc. The mixture was transferred to a separatory funnel, washed with sat. NaHCO_3_, dried with magnesium sulfate, concentrated, and purified via column chromatography.

##### Synthesis of 5

Compound 5 was obtained by using the aforementioned standard coupling conditions with 49 mg (0.203 mmol) of compound **2**, 45 mg (0.229 mmol) of compound **4**, 43 mg (0.277 mmol) of EDC, and 29 mg (0.214 mmol) of HOAt. The crude reaction mixture was purified by preparatory HPLC.

##### Synthesis of 8

Desthiobiotin (**6**, 200 mg, 0.93 mmol), N-hydroxysuccinimide (241 mg, 1.11 mmol), and EDC-HCl (213 mg, 1.11 mmol) were stirred in DMF under a nitrogen atmosphere for 12 hours. The crude mixture was diluted with EtOAc, washed with water, dried with magnesium sulfate, and concentrated (used without further purification).

To a solution of desthiobiotin NHS-ester (100 mg, 0.32 mmol) in dichloromethane, N,N-diisopropylethylamine (42 mg, 0.32 mmol) and 14-azido-3,6,9,12-tetraoxatetradecan-1-amine (83 mg, 0.32 mmol) were added. The reaction mixture was stirred overnight at room temperature under a nitrogen atmosphere. The solvent was then removed in vacuo, and the resulting oil was subjected to flash chromatography to isolate compound **8** (*N*-(14-azido-3,6,9,12-tetraoxatetradecyl)-6-((4*R*,5*S*)-5-methyl-2-oxoimidazolidin-4-yl)hexanamide).

##### Synthesis of SH-0029-DTB

In a round bottom flask, compound **5** (0.100 g, 0.240 mmol), (+)-sodium l-ascorbate (8.5 mg, 0.048 mmol), copper(II) sulfate powder (7.6 mg, 0.048 mmol), and compound **8** (110 mg, 0.240 mmol) were added. The atmosphere was purged with argon gas, then DMF (3 mL) was added. The reaction mixture was stirred at RT for 16 h. The solvent was removed in vacuo and directly purified via column chromatography to isolate **SH-0029-DTB**. ^1^H NMR (400 MHz, Methanol-D_4_) δ 8.16 (s, 1H), 7.73 (s, 1H), 7.63-7.61 (m, 3H), 7.32 (s, 1H), 7.12 (d, J = 9.5 Hz, 1H), 6.47-6.36 (m, J = 42.9 Hz, 2H), 5.79 (dd, J = 8.8, 3.0 Hz, 1H), 5.25 (s, 2H), 4.59 (t, J = 4.9 Hz, 2H), 3.87 (t, J = 5.0 Hz, 2H), 3.78-3.52 (m, 12H), 3.45 (t, J = 5.5 Hz, 2H), 3.36-3.32 (m, 6H), 3.11-3.04 (m, 1H), 2.72 (dd, J = 16.0, 5.5 Hz, 1H), 2.42 (dd, J = 15.8, 7.0 Hz, 1H), 2.13 (t, J = 7.4 Hz, 2H), 1.59-1.53 (m, J = 7.5 Hz, 2H), 1.51-1.26 (m, 12H), 1.04 (d, J = 6.5 Hz, 3H). EI MS m/z: pos 876.4 (MH^+^).

**Figure 14.**
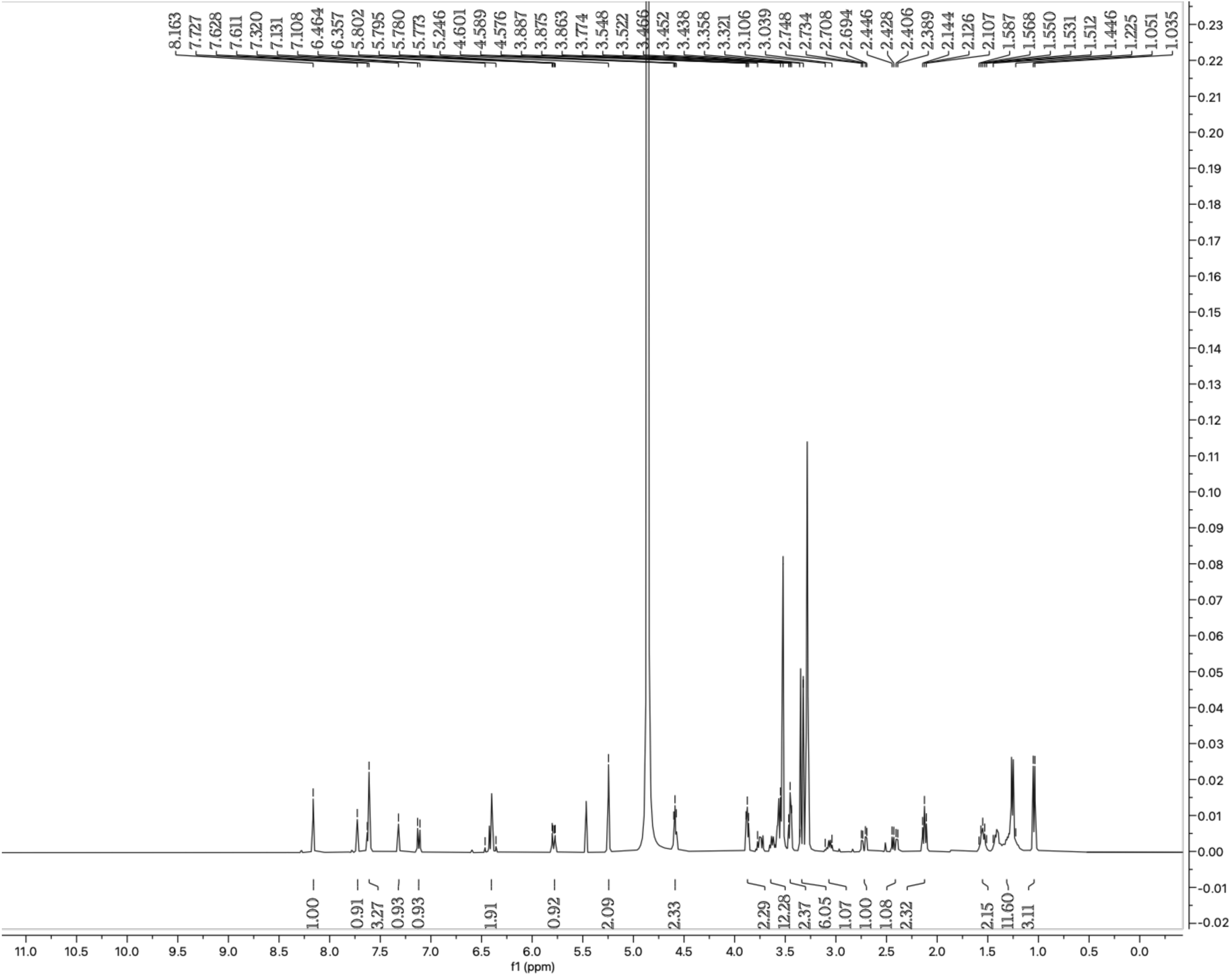
^1^H NMR spectrum of **SH-0029-DTB**.

#### SH-0017

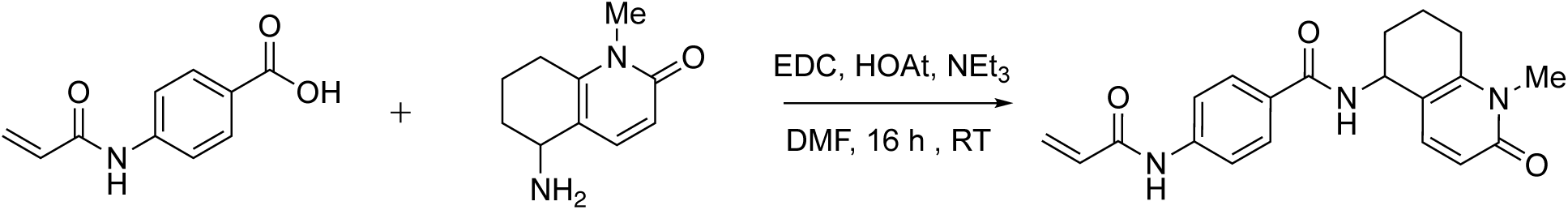

N-(1-methyl-2-oxo-1,2,5,6,7,8-hexahydroquinolin-5-yl)-4-(prop-2-enamido)benzamide was obtained by using the aforementioned standard coupling conditions with 49 mg (0.275 mmol) of 5-amino-1-methyl-1,2,5,6,7,8-hexahydroquinolin-2-one, 57mg (0.298mmol) of 4-(prop-2-enamido)benzoic acid, 57 mg (0.367 mmol) of EDC, and 39.3 mg (0.289 mmol) of HOAt. Purified by HPLC procedure 2 (gradient: from A-70%: B-30% to A-20%: B-80%; Rf = 0.56; run time = 6.5 min). Yield: 52.9 mg (55.5 %). Yellow powder. LCMS purity: 100 % (LCMS procedure 1, Rf = 0.39, run time = 2 min). ^1^H NMR (400 MHz, Methanol-D_4_) δ 7.83 (d, *J* = 8.9 Hz, 2H), 7.74 (d, *J* = 9.0 Hz, 2H), 7.42 (d, *J* = 9.3 Hz, 1H), 6.49-6.36 (m, 3H), 5.80 (dd, *J* = 9.3, 2.6 Hz, 1H), 5.16-5.15 (m, 1H), 3.56 (s, 3H), 2.87-2.73 (m, 2H), 2.1-1.91 (m, 4H). EI MS m/z: pos. 352.2 (M+H^+^).

**Figure 15.**
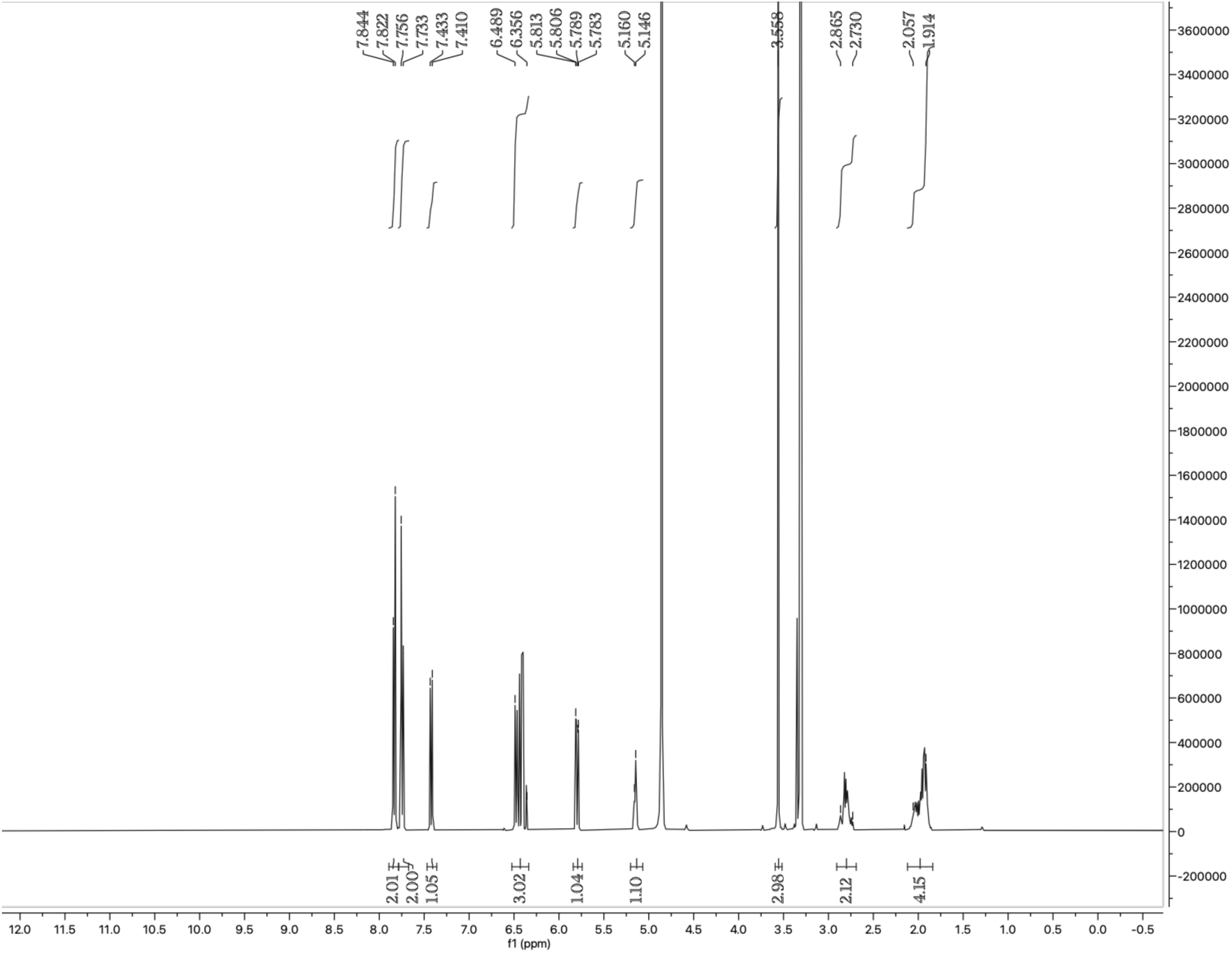
^1^H NMR spectrum of **SH-0017**.

#### SH-0087

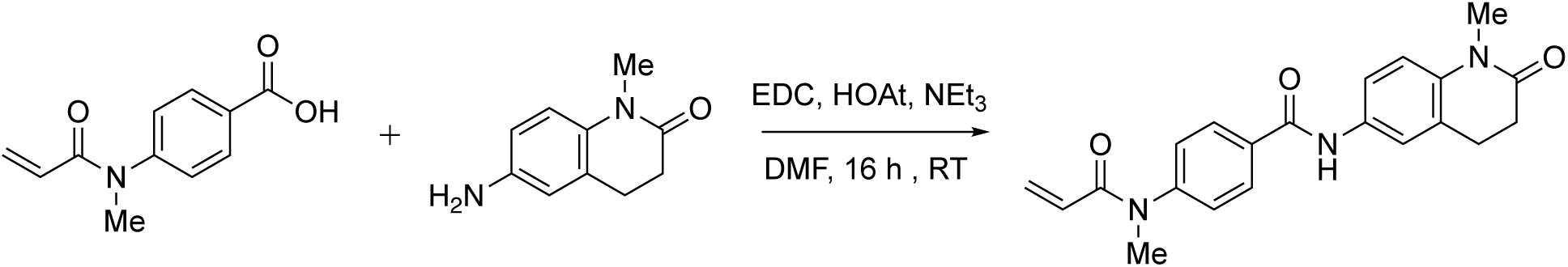

N-(1-methyl-2-oxo-1,2,3,4-tetrahydroquinolin-6-yl)-4-(N-methylprop-2-enamido)benzamide was obtained by using the aforementioned standard coupling conditions with 59 mg (0.335 mmol) of 6-amino-1-methyl-1,2,3,4-tetrahydroquinolin-2-one, 69 mg (0.336 mmol) of 4-(N-methylprop-2-enamido)benzoic acid, 52 mg (0.335 mmol) of EDC, and 47.9 mg (0.352 mmol) of HOAt. Purified by HPLC procedure 1 (gradient: from A-75%: B-25% to A-50%: B-50%). Yield: 69.4 mg (62.5 %). Beige powder. LCMS purity: 100 % (LCMS procedure 1, Rf = 0.58, run time = 2 min). ^1^H NMR (400 MHz, Methanol-D_4_) δ 8.04-8.02 (m, 2H), 7.64-7.60 (m, 2H), 7.44-7.41 (m, 2H), 7.14 (d, *J* = 8.5 Hz, 1H), 6.31 (dd, *J* = 16.9, 2.1 Hz, 1H), 6.21-6.15 (m, 1H), 5.63 (dd, *J* = 10.2, 2.2 Hz, 1H), 3.38 (d, *J* = 8.6 Hz, 6H), 2.96-2.93 (m, 2H), 2.66-2.63 (m, 2H). EI MS m/z: pos. 364.2 (M+H^+^)

**Figure 16.**
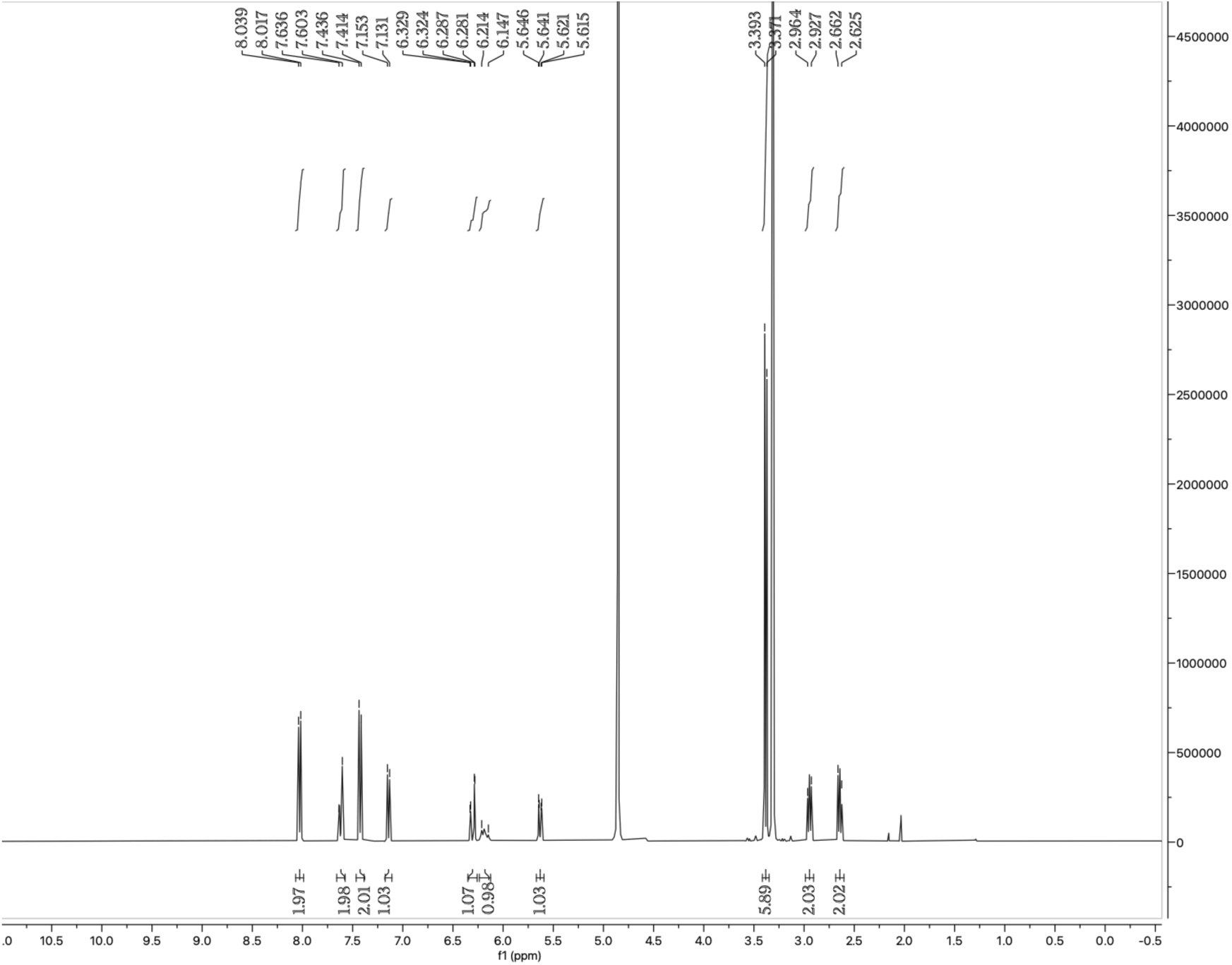
^1^H NMR spectrum of **SH-0087**.

#### SH-0077

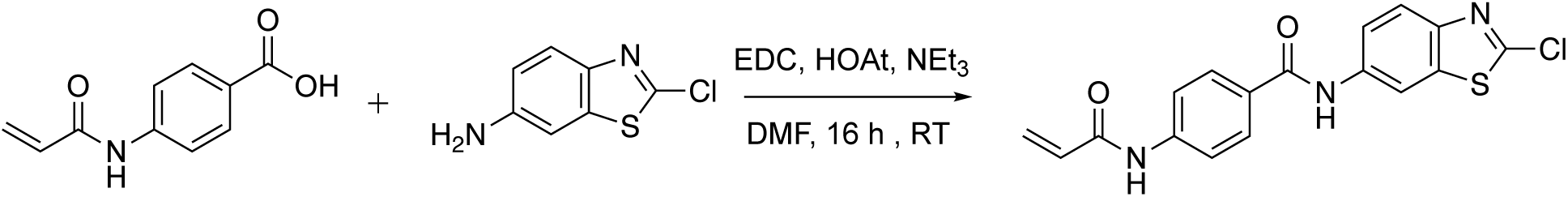

N-(2-chloro-1,3-benzothiazol-6-yl)-4-(prop-2-enamido)benzamide was obtained by using the aforementioned standard coupling conditions with 52 mg (0.283 mmol) of 2-chloro-1,3-benzothiazol-6-amine, 59 mg (0.309 mmol) of 4-(prop-2-enamido)benzoic acid, 52 mg (0.335 mmol) of EDC, and 40.4 mg (0.297mmol) of HOAt. Purified by HPLC procedure1 (gradient: from A-60%: B-40% to A-35%: B-65%). Yield: 13.1 mg (13 %). Yellow powder. LCMS purity: 100 % (LCMS procedure 1, Rf = 0.65, run time = 2 min). ^1^H NMR (400 MHz, Methanol-D_4_) δ 8.50 (d, *J* = 2.1 Hz, 1H), 7.97 (dm, *J* = 8.9 Hz, 2H), 7.89 (d, *J* = 8.8 Hz, 1H), 7.83 (dm, *J* = 8.9 Hz, 2H), 7.75 (dd, *J* = 8.9, 2.1 Hz, 1H), 6.51-6.38 (m, 2H), 5.82 (dd, *J* = 9.3, 2.6 Hz, 1H). EI MS m/z: pos. 358.0 (M+H^+^).

**Figure 17.**
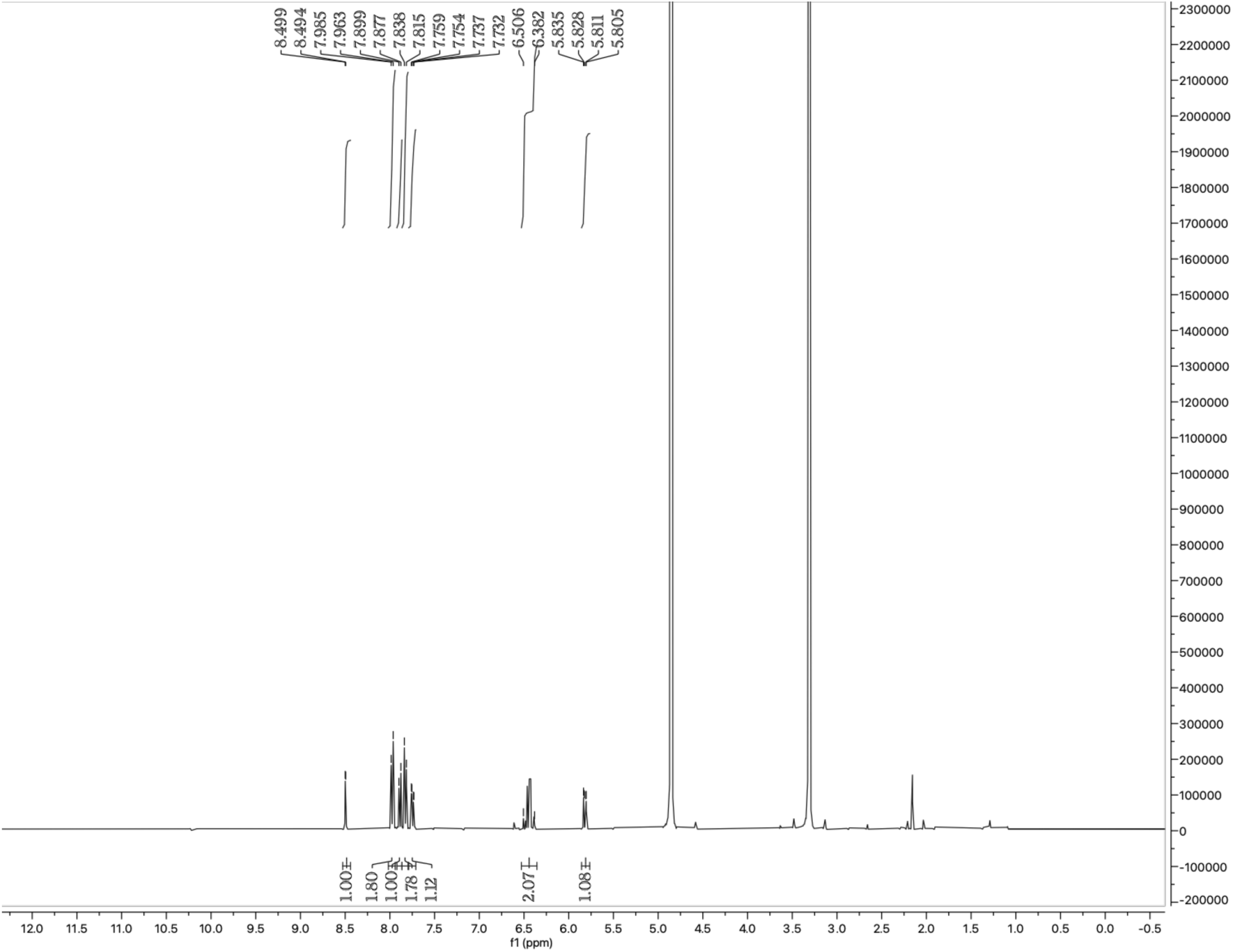
^1^H NMR spectrum of **SH-0077**.

#### SH-0013

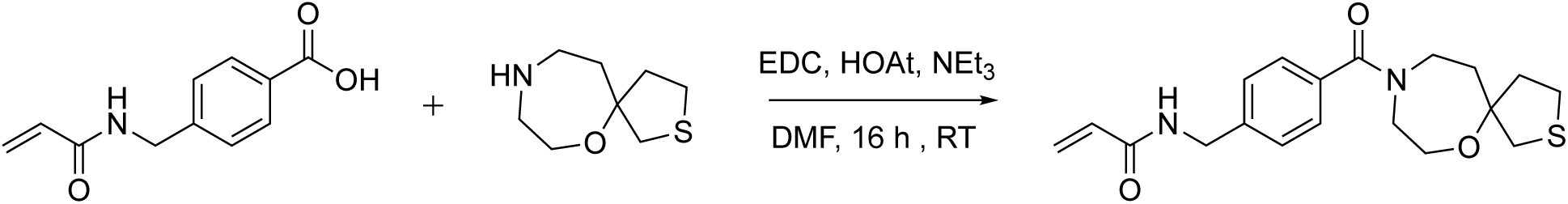

N-[(4-{6-oxa-2-thia-9-azaspiro[4.6]undecane-9-carbonyl}phenyl)methyl]prop-2-enamide was obtained by using the aforementioned standard coupling conditions with 46 mg (0.22 mmol) of 6-oxa-2-thia-9-azaspiro[4.6]undecane hydrochloride, 54 mg (0.263 mmol) of 4-[(prop-2-enamido)methyl]benzoic acid, 54 mg (0.348 mmol) of EDC, 33 mg (0.326 mmol) of Et3N, and 31.4 mg (0.231 mmol) of HOAt. Purified by HPLC procedure 4 (gradient: from A-85%: B-15% to A-35%: B-65%; Rf = 0.81; run time = 6.5 min). Yield: 50 mg (58 %). Pink powder. LCMS purity: 100 % (LCMS procedure 1, Rf = 0.52, run time = 2 min). ^1^H NMR (400 MHz, Methanol-D_4_) δ 7.40-7.39 (m, 4H), 6.33-6.23 (m, 2H), 5.69 (dd, *J* = 8.5, 3.5 Hz, 1H), 4.48 (s, 2H), 3.94-3.54 (m, 6H), 3.00-2.90 (m, 2H), 2.83-2.63 (d, *J* = 57.7 Hz, 2H), 2.27-2.08 (m, 3H), 1.84-1.68 (m, 1H). EI MS m/z: pos. 361.2 (M+H^+^).

**Figure 18.**
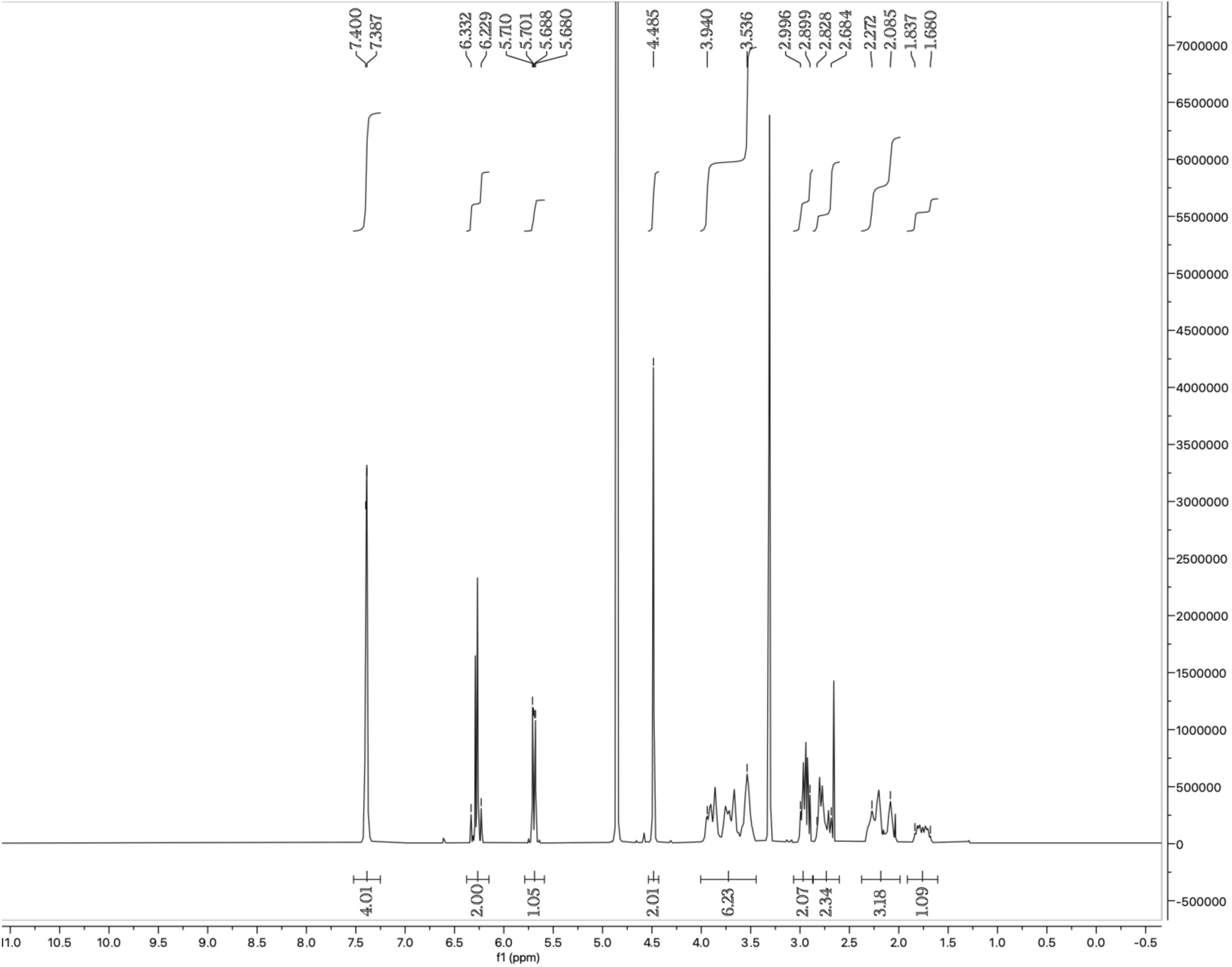
^1^H NMR spectrum of **SH-0013**.

#### SH-0035

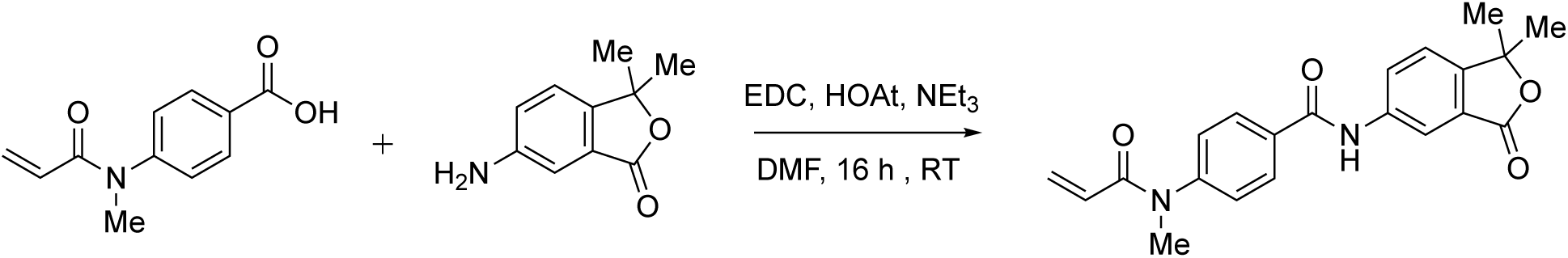

N-(1,1-dimethyl-3-oxo-1,3-dihydro-2-benzofuran-5-yl)-4-(N-methylprop-2-enamido)benzamide was obtained by using the aforementioned standard coupling conditions with 36 mg (0.203 mmol) of 6-amino-3,3-dimethyl-1,3-dihydro-2-benzofuran-1-one, 47 mg (0.229 mmol) of 4-(N-methylprop-2-enamido)benzoic acid, 43 mg (0.277 mmol) of EDC, and 29.1 mg (0.214 mmol) of HOAt. Purified by HPLC procedure 3 (gradient: from A-65%: B-35% to A-40%: B-60%). Yield: 35.8 mg (47.2 %). Beige powder. LCMS purity: 100 % (LCMS procedure 1, Rf = 0.55, run time = 2 min). ^1^H NMR (400 MHz, MeOD-D_4_) δ 8.26 (d, *J* = 2.0 Hz, 1H), 8.07-8.04 (m, 3H), 7.62 (d, *J* = 8.4 Hz, 1H), 7.45 (d, *J* = 8.6 Hz, 2H), 6.31 (dd, *J* = 16.8, 2.1 Hz, 1H), 6.22-6.15 (m, 1H), 5.64 (dd, *J* = 10.2, 2.2 Hz, 1H), 3.40 (s, 3H), 1.68 (s, 6H). EI MS m/z: pos. 365.0 (M+H^+^).

**Figure 19.**
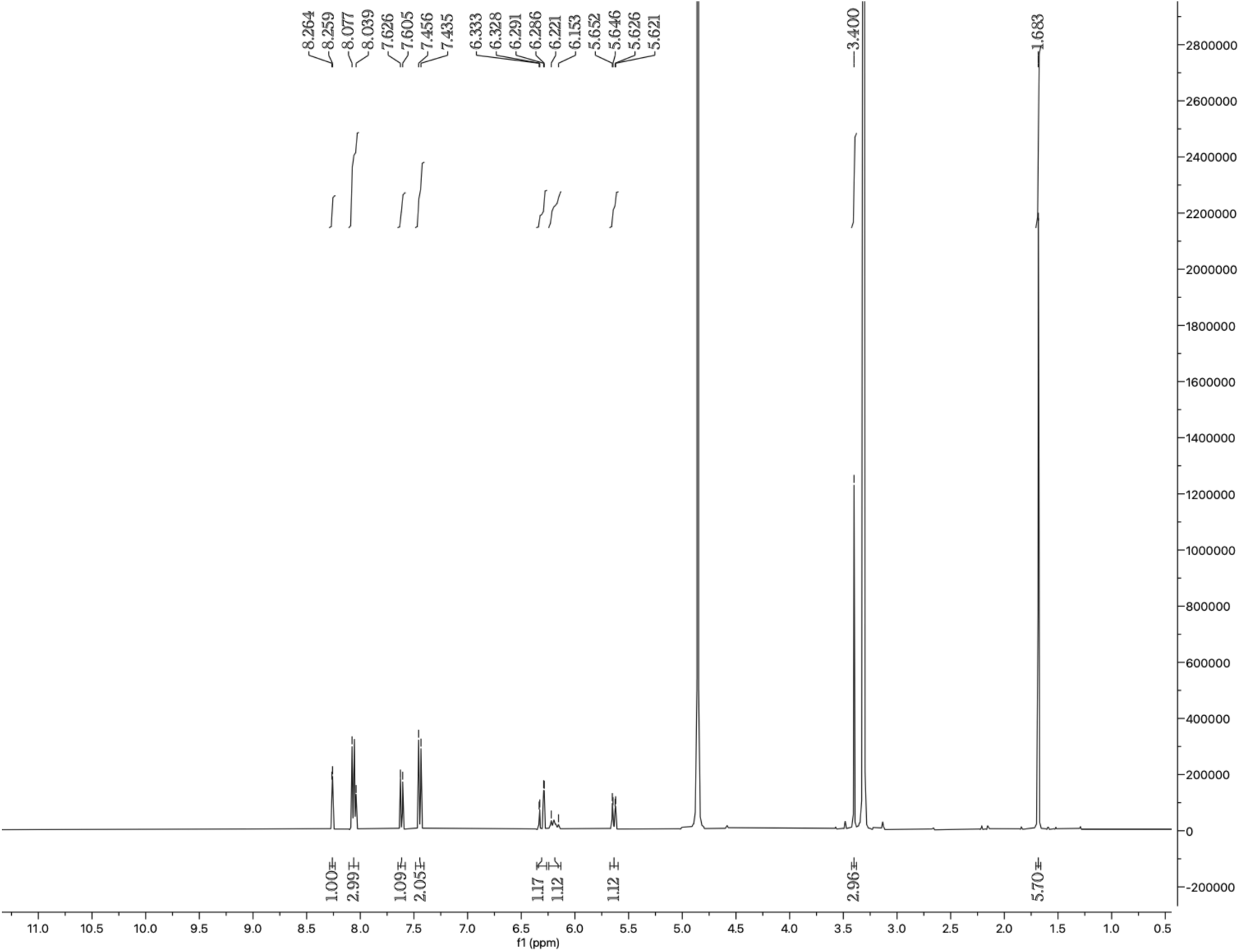
^1^H NMR spectrum of **SH-0035**.

#### 2-A01

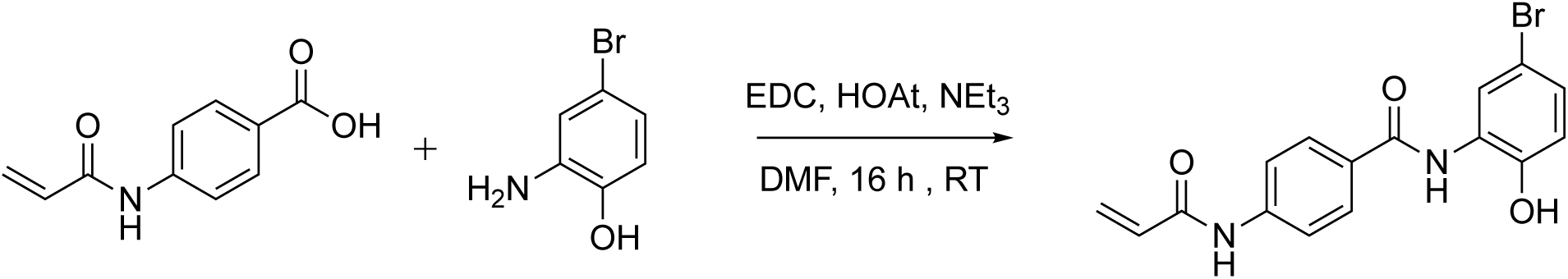

4-acrylamido-N-(5-bromo-2-hydroxyphenyl)benzamide was obtained by using the aforementioned standard coupling conditions with 38 mg (0.203 mmol) of 4-acrylamidobenzoic acid, 43 mg (0.229 mmol) of 2-amino-4-bromophenol, 43 mg (0.277 mmol) of EDC, and 29 mg (0.214 mmol) of HOAt. Purified by preparatory HPLC. Beige powder. ^1^H NMR (400 MHz, DMSO) δ 10.44 (s, 1H), 10.21 (bs, 1H), 9.39 (s, 1H), 7.97-7.93 (m, 2H), 7.82-7.80 (m, 2H), 7.18 (dd, *J* = 8.6, 2.5 Hz, 1H), 6.88 (d, *J* = 8.6 Hz, 1H), 6.47 (dd, *J* = 17.0, 10.1 Hz, 1H), 6.31 (dd, *J* = 17.0, 2.0 Hz, 1H), 5.81 (dd, *J* = 10.0, 2.0 Hz, 1H). EI MS m/z: pos. 361.0 (MH^+^).

**Figure 20.**
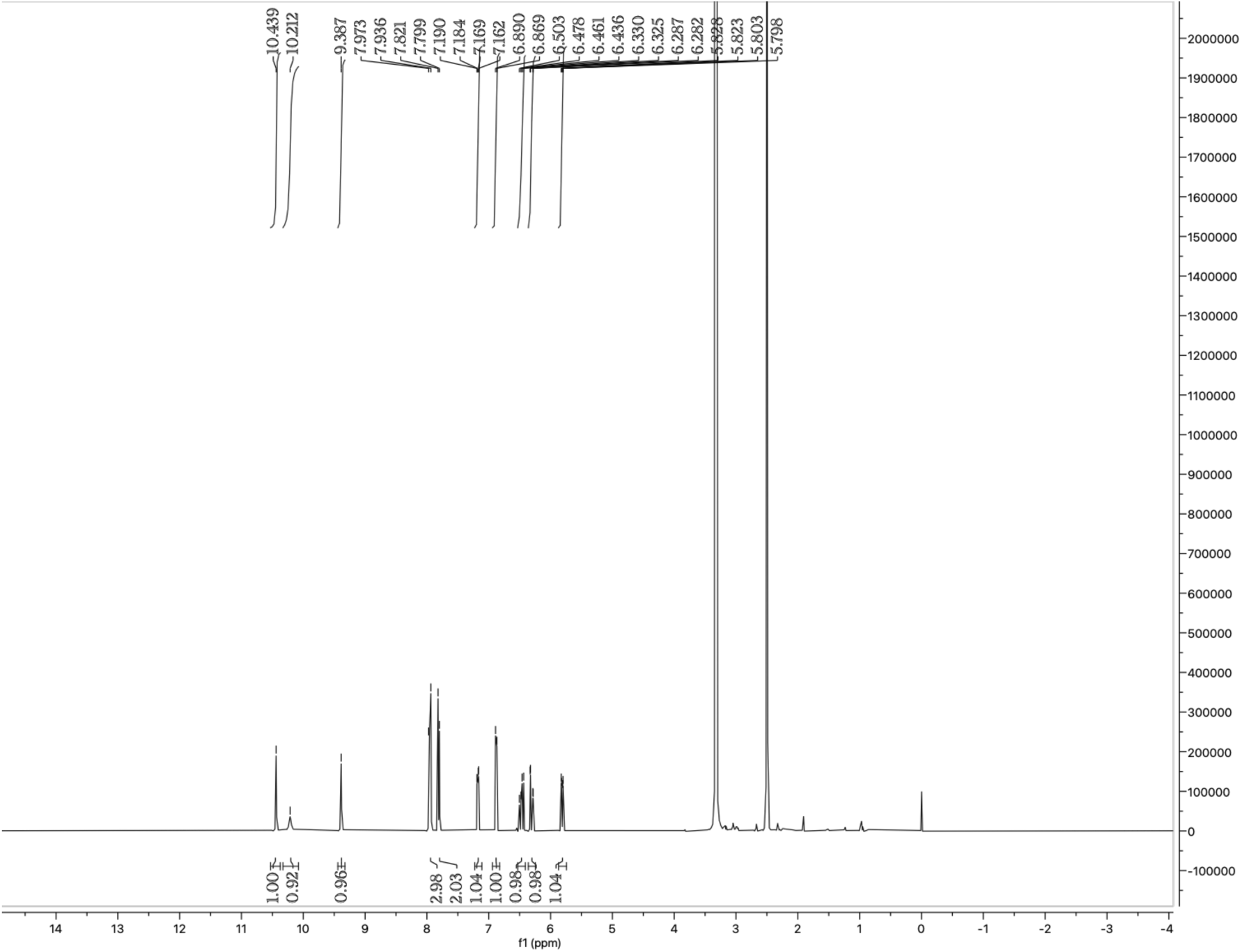
^1^H NMR spectrum of **2-A01.**

#### SH-0081

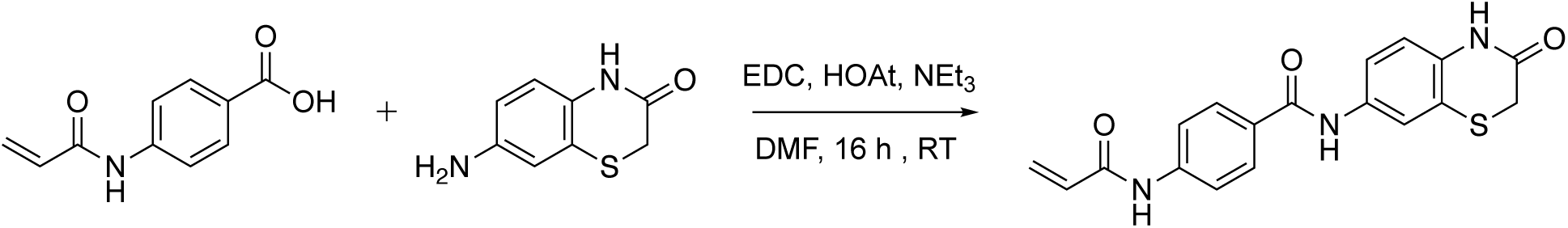

4-acrylamido-N-(3-oxo-3,4-dihydro-2H-benzo[b][1,4]thiazin-7-yl)benzamide was obtained by using the aforementioned standard coupling conditions with 38 mg (0.203 mmol) of 4-acrylamidobenzoic acid, 41 mg (0.229 mmol) of 7-amino-2*H*-benzo[*b*][1,4]thiazin-3(4*H*)-one, 43 mg (0.277 mmol) of EDC, and 29 mg (0.214 mmol) of HOAt. Purified by preparatory HPLC. Beige powder. EI MS m/z: pos. 354.1 (MH^+^).

**Figure 21.**
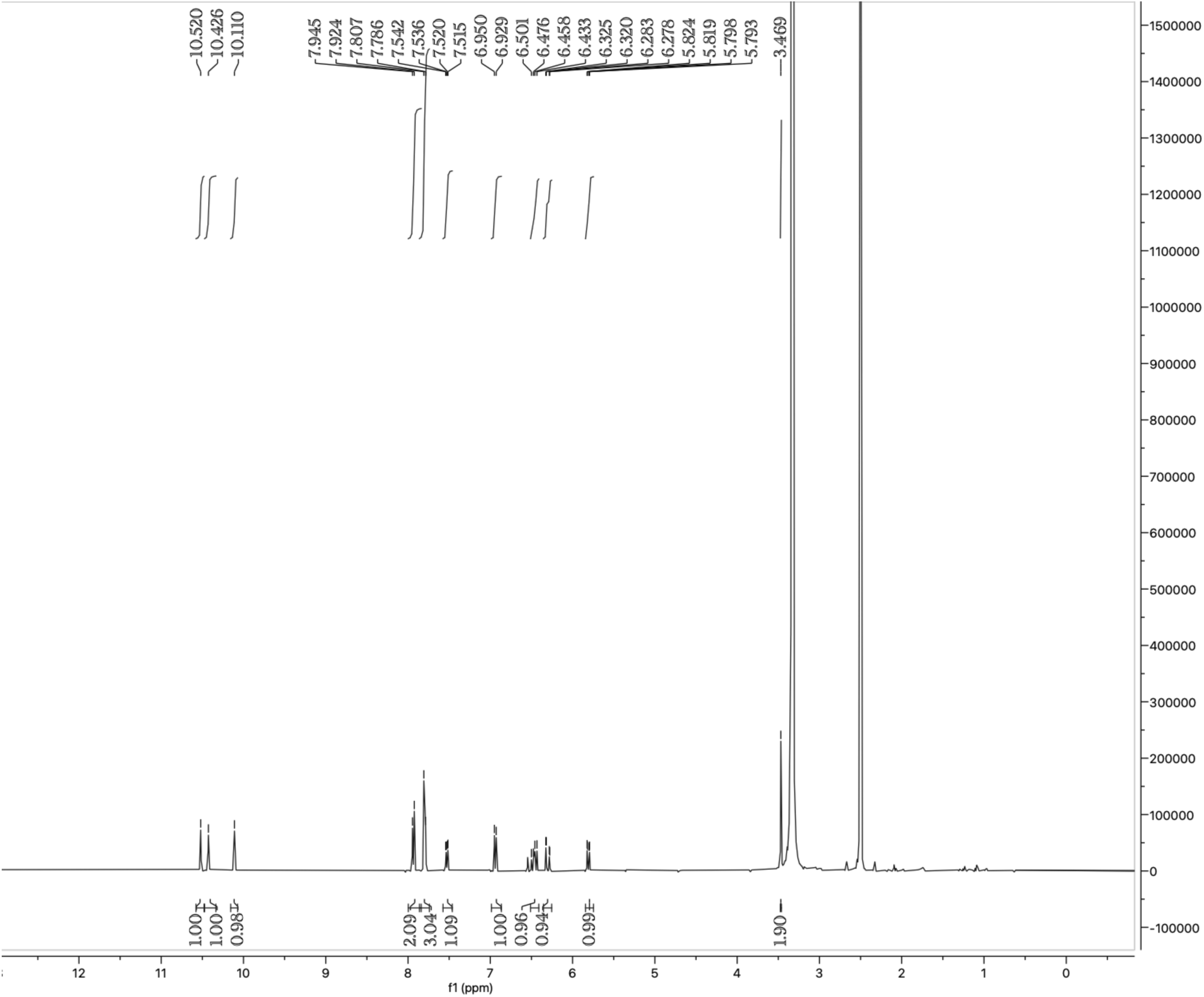
^1^H NMR spectrum of **SH-0081**.

#### *N*-(2-(2-iodoacetamido)ethyl)-6-((4*R*,5*S*)-5-methyl-2-oxoimidazolidin-4-yl)hexanamide (DBIA)

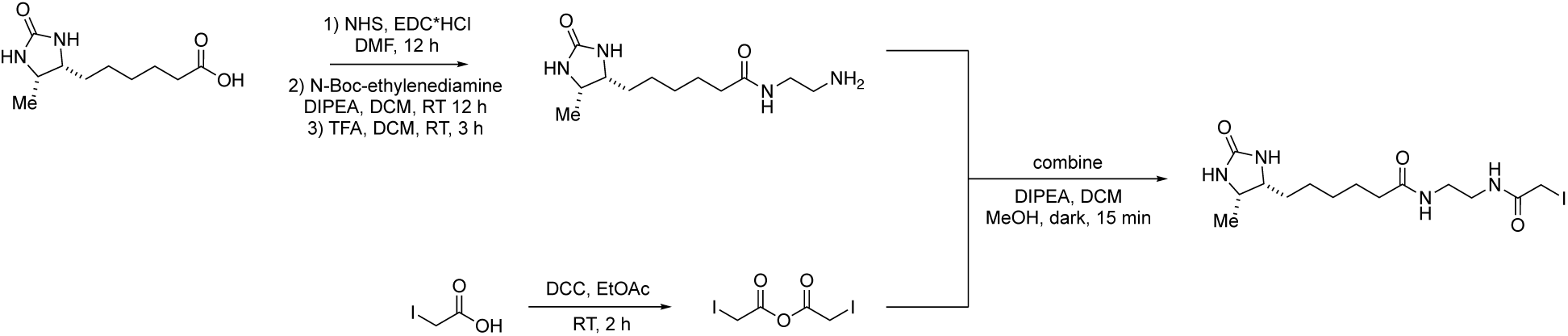

Desthiobiotin (200 mg, 0.93 mmol), N-hydroxysuccinimide (241 mg, 1.11 mmol), and EDC-HCl (213 mg, 1.11 mmol) were stirred in DMF under a nitrogen atmosphere for 12 hrs^158^. The crude mixture was diluted with EtOAc, washed with water, dried with magnesium sulfate, and concentrated (used without further purification).

To a solution of desthiobiotin, NHS-ester (100 mg, 0.32 mmol) in dichloromethane, N,N-diisopropylethylamine (0.32 mmol) and N-boc ethylenediamine (0.32 mmol) were added^165^. The reaction mixture was stirred overnight at room temperature under a nitrogen atmosphere. The solvent was then removed in vacuo, and the resulting oil was dispersed in 20% TFA in dichloromethane (DCM) and stirred for 3 hrs at room temperaturee. The TFA and DCM were removed in vacuo to yield a yellow oil, which was washed repeatedly with diethyl ether to remove impurities.

Iodoacetic acid^159^ (8.0 g, 43.0 mmol) and DCC (4.2 g, 20.5 mmol) were stirred in EtOAc (120 mL) for 2 hrs under a nitrogen atmosphere. The crude mixture was filtered and concentrated for direct use in the next reaction.

N-(2-aminoethyl)-6-((4R,5S)-5-methyl-2-oxoimidazolidin-4-yl)hexanamide^166^ (0.182 g, 0.71 mmol) dispersed in 10 mL of methanol/DCM (1:9 v/v), to which was added N,N-diisopropylethylamine (0.71 mmol) and iodoacetic anhydride (378 mg, 1.07 mmol). The reaction was stirred in the dark for 1 h at room temperature. The reaction mixture was concentrated, then triturated with acetone to isolated DBIA as a white solid. Note: the compound is light sensitive. Thus, work was done in low light conditions and the flask was wrapped in foil. ^1^H NMR (400 MHz, DMSO-*D*_6_) δ 8.23 (s, 1H), 7.76 (s, 1H), 6.27 (s, 1H), 6.08 (s, 1H), 3.57-3.53 (m, 3H), 3.46-3.39 (m, 1H), 3.03 (bs, 4H), 2.00 (t, *J* = 7.4 Hz, 2H), 1.46-1.40 (m, 2H), 1.29-1.17 (m 6H), 0.91 (d, *J* = 6.4 Hz, 3H). EI MS m/z: pos. 425.1 (M+H^+^).

**Figure 22.**
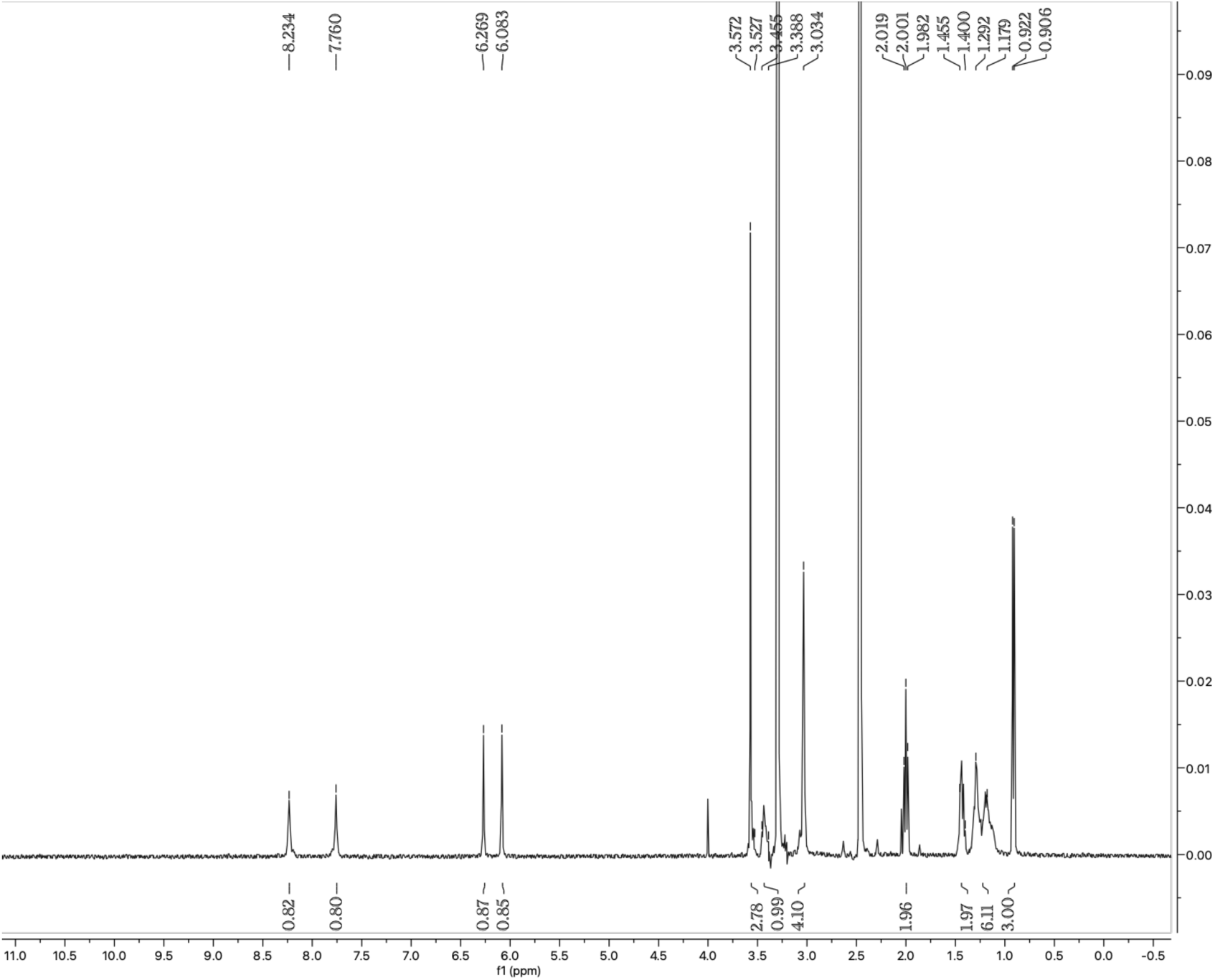
^1^H NMR spectrum of **DBIA**.

## Supplemental Data Tables

Table S1. Culture methods of 416 cancer cell lines analyzed in DrugMap, related to Figure 1.

Table S2. Cysteine ligandability analyzed in DrugMap, related to Figure 1.

Table S3. List of heterogeneous cysteines, related to Figure 2.

Table S4. Cysteine ligandability analyzed in iso-TMT experiments of K562 treated with TMCEP (A) or KI696 (B), related to Figure 2.

Table S5. Cysteine ligandability analyzed in iso-TMT experiments of HEK-293T overexpressing FLAG-UGDH treated with 1 mM UGDH, related to Figure 2.

Table S6. Cysteine ligandability analyzed in iso-TMT experiments of PC9 or K562 treated with 1 μM Osimertinib or Selinexor, respectively, related to Figure 2.

Table S7. Cysteine ligandability analyzed in iso-TMT experiments of HEK-293T overexpressing FLAG-PRDX5 or FLAG-PRDX5•F157L, related to Figure S6.

Table S8. Cysteine ligandability analyzed in iso-TMT experiments of MM1S treated with 50 uM SH-7971 or SH-1696, related to Figure 3.

Table S9. Cysteine ligandability analyzed in iso-TMT experiments of SKMEL5 treated with 10 uM SH-0105 or SH-0029, related to Figure 4.

Table S10. RNAseq analysis of 7 melanoma transfected with siSOX10 or siCTRL, related to Figure S10.

Table S11. RNAseq analysis of COLO679, IGR1, and U257 treated with 3 µM of SH-0105, SH-0029 for 48 hrs, related to Figure S10.

Table S12. RNAseq analysis of SKMEL5 transfected with siSOX10 or siCTRL treated with 2.5 µM of SH-0105 or SH-0029 for 48 hrs, related to Figure S10.

Table S13. Structural parameters, related to Figure 1.

Table S14. sgRNA plasmid, siRNA and primers used in this study, related to STAR Methods.

Table S15. Crystallographic Data and Refinement Statistics, related to Figure 3 and S7.

Table S16. CSEA cysteine sets, related to Figure 1.

